# CK2 inhibition suppresses glial inflammation in the brain

**DOI:** 10.1101/2025.08.05.668554

**Authors:** Ioana I. N. Da Silva, Desiree Ramirez, Sarah L. Parylak, Leslie A. Wallace, James K. Tucker, Joel M. Erberich, Rutvi Katariya, Aidan H. McDonald, Iryna S. Gallina, Ariana L. Tucker, Jillybeth Burgado, Baptiste N. Jaeger, Jerika J. Barron, Joshua M. Pratt, Monique Pena, Vipula Racha, Christina K. Lim, Sarah Fernandes, Simone Benassi, Lynne Randolph-Moore, Krishna C. Vadodaria, Maria C. Marchetto, Nicola J. Allen, Fred H. Gage

**Affiliations:** Laboratory of Genetics, The Salk Institute for Biological Studies, 10010 North Torrey Pines Road, La Jolla, CA 92037-1002, USA; Molecular Neurobiology Laboratory, The Salk Institute for Biological Studies, 10010 North Torrey Pines Road, La Jolla, CA 92037-1002, USA; Department of Anthropology, University of California, San Diego, 9500 Gilman Drive MC 0532, La Jolla, CA 92093-0532, USA

## Abstract

Neuroinflammation plays a key role in Alzheimer’s disease (AD) and related neurodegenerative disorders. Chronic activation of astrocytes and microglia fuels neuronal damage via cytokine secretion, oxidative stress, and proteolysis. However, glial inflammatory regulation remains poorly understood. Using chemoproteomics, we identified CK2, particularly the brain-enriched catalytic subunit CK2α2, as a key driver of astrocytic inflammation. CK2 enhances NF-κB activity by phosphorylating NF-κB S529 and IκBα S32, promoting pro-inflammatory gene expression. CK2 inhibition via genetic or chemical approaches dampens inflammation, including IL-6 and IL-8 expression in an acute neuroinflammation mouse model. CK2α2 is upregulated in AD postmortem tissues and patient-derived astrocytes. AD astrocytes exhibit a hyperinflammatory state that can be attenuated by CK2 inhibition. Overexpression of CK2α2 in cortical organoids mimics AD pathology, whereas CK2 inhibition using the potent, selective, and brain-penetrant probe TAL606 rescues inflammatory markers in transgenic AD mice. These findings position CK2 as a central regulator of neuroinflammation and a promising therapeutic target for AD and related disorders.

## Introduction

Neuroinflammation has emerged as a significant etiological factor for numerous neurological and psychiatric diseases, including Alzheimer’s disease (AD)^1–3^. During age-related neurodegeneration, microglia and astrocytes, the brain’s resident immune cells, become chronically activated and perpetuate a destructive loop of pro-inflammatory cytokine release^4–6^. Chronic inflammation in AD is evidenced by the presence of reactive glia in postmortem studies and inflammatory biomarkers in serum and CSF^7,8^. Despite the emergence of neuroinflammation as a therapeutic target, the mechanisms regulating inflammation in glia remain poorly understood.

Phenotype-guided studies have been a rich source of novel targets with therapeutic applications in human diseases^9^. The flavonoid natural product family has been shown to possess varying levels of anti-inflammatory activity in human and mouse-derived immune cells, including monocytes and microglia^10,11^. However, efforts to elucidate flavonoids’ direct cellular targets and to ascribe specific flavonoid structural features to anti-inflammatory activity have been lacking.

Using a chemoproteomics approach in IL1β-activated astrocytes, we identified the kinase CK2 as a target of natural product flavones known for their anti-inflammatory properties^12–14^. CK2 is a protein kinase that is catalytically active either as a heterotetrameric complex composed of 2 alpha and 2 beta subunits or as an alpha monomer^15^. Catalytic alpha subunits CSNK2A1/CK2α1 and CSNK2A3/CK2α1P are nearly identical (99% cDNA homology^16^), whereas CSNK2A2/CK2α2 shares 77% identity with CSNK2A1. CK2β, the regulatory subunit, can regulate CK2 activity both positively and negatively^17^. Both CK2α1 and CK2α2 are highly expressed in the brain, but CK2α2 is enriched in the brain and testes relative to other tissues^18^.

We show that CK2 is an upstream regulator of glial inflammation and that pharmacological inhibition or genetic perturbation of CK2, especially brain-enriched catalytic subunit isoform CK2α2, reduces biochemical and transcriptional programs driving inflammation in glia. NF-κB is a major transcriptional driver of inflammation. We find that CK2 modulates NF-κB activity via direct phosphorylation of NF-κB S529 and IκBα S32, leading to upregulation of NF-κB transcriptional signatures. Furthermore, CK2 inhibition blocks IL-6 and IL-8 expression in an acute neuroinflammation mouse model. We show that CK2 activity is dysregulated in AD patients, with elevated CK2α2 expression in AD astrocytes and microglia. In AD patient-derived astrocytes, CK2 overactivation is correlated with a hyperinflammatory state that can be blocked by CK2 inhibition. Physiological overexpression of CK2α2 promotes AD-like phenotypes in glia-enriched cortical organoids. Using the potent, selective, and brain-penetrant chemical probe TAL606, we show that inhibition of CK2 in APP/PS1 transgenic AD mice leads to suppression of neuroinflammation indicators after one month of treatment. These findings highlight CK2 as a potent regulator of neuroinflammation, positioning CK2 inhibition as a putative therapeutic avenue for the treatment of neurodegenerative diseases.

## RESULTS

### Anti-inflammatory activity of flavones in human microglia and astrocytes

We used a flow cytometry-based assay to assess the anti-inflammatory activity of several natural flavonoid compounds (Extended Data Fig. 1A) and to identify corresponding structure-activity relationships. Primary human cerebellar astrocytes (HCA) were co-treated with 20 µM of each compound or vehicle (DMSO), 10 ng/mL IL1-β, and a protein transport inhibitor for 6 hours, and were then processed for intracellular staining of pro-inflammatory cytokines IL-6, IL-8, TNF-α, and MCP-1. We found an unsaturated C2-C3 bond was the predominant determining factor for anti-inflammatory activity. For example, apigenin (API) strongly blocked IL1-β-induced IL-6, IL-8, TNF-α, and MCP-1 cytokine upregulation, unlike its flavanone analogue naringenin (NAR; Fig. 1A and Extended Data Fig. 1B). A similar divergence in activity was observed with trimethylated apigenin (4′,5,7-trimethoxyflavone, TMF) and its corresponding inactive flavanone analogue 4′,5,7-trimethoxyflavanone (dTMF, Extended Data Fig. 1C) as well as chrysoeriol (CHR) versus homoeriodictyol (dCHR; Extended Data Fig. 1A,D). For subsequent phenotypic testing, we used active flavones API, TMF, or CHR; API and CHR were initially identified as among the most potent flavones in our SAR studies, and we continued to characterize TMF given its potential as a more bioavailable prodrug of API. We broadly assessed the effects of CHR treatment on cytokine release in lipopolysaccharide (LPS)-stimulated human induced pluripotent stem cell (iPSC)-derived microglia-like cells (iMGL)^19^ using a panel of 36 cytokines (Fig. 1B). CHR significantly blocked the release of multiple LPS-induced pro-inflammatory cytokines, including interleukins (IL-6, IL-8, IL1ra), chemokines (CXCL1, MCP-1, MIP-1), complement (C5a), and TNF-α (p < 0.05, unpaired two-tailed t-tests). Altered phagocytic function of microglia is associated with neurodegenerative disease.^20^ CHR blocked microglial phagocytosis of pHrodo E. coli beads (P = 0.0015, Fig. 1C; compare to phagocytosis inhibitor cytochalasin D^21^). The anti-inflammatory effects of CHR (Fig. 1D) and TMF (Extended Data Fig. 1C) were dose-dependent in astrocytes and microglia. There was no cytotoxicity associated with flavone treatment (Extended Data Fig. 1E).

**Figure 1.**
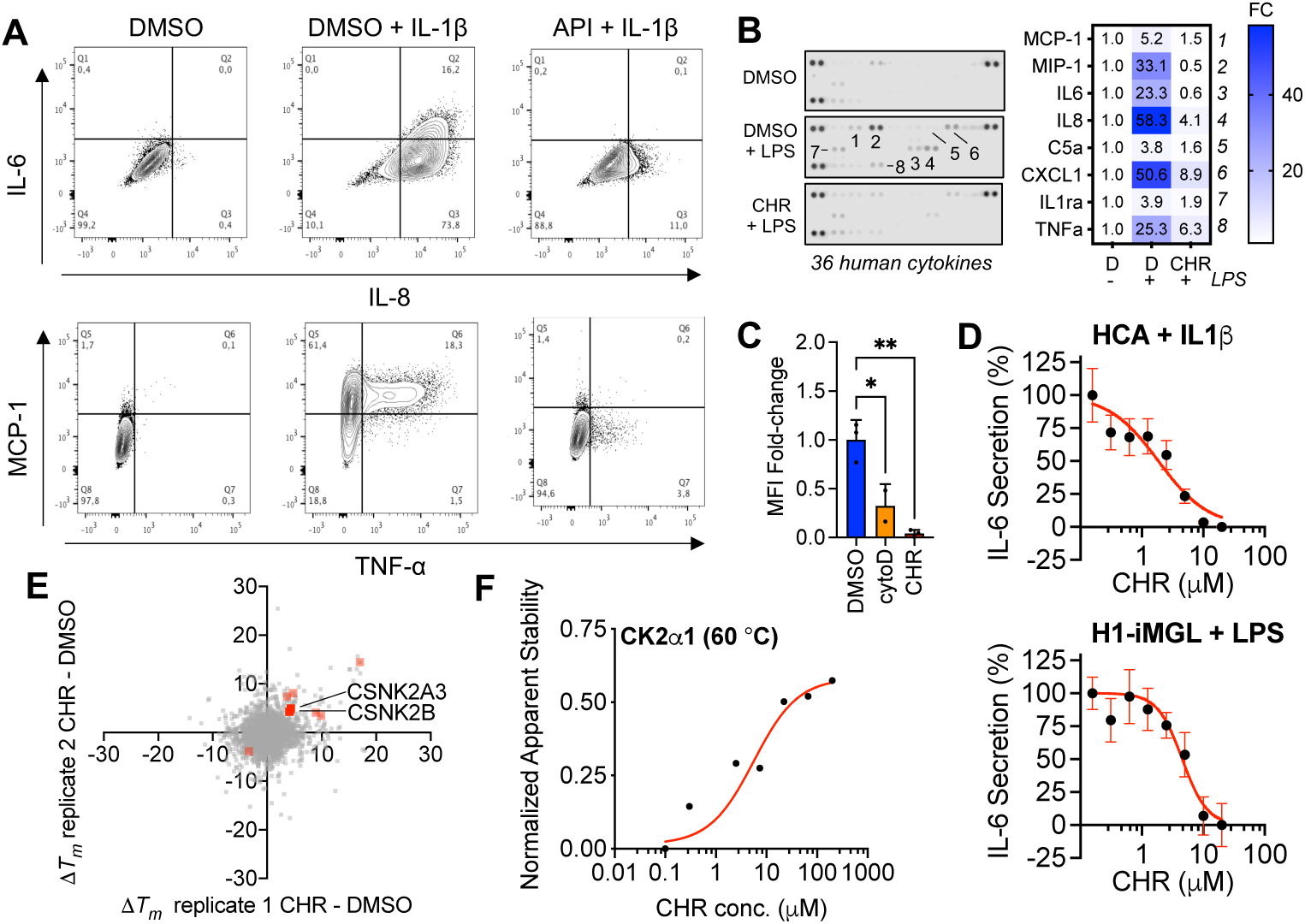
Anti-inflammatory activity and unbiased target identification of flavones in microglia and astrocytes. A. Apigenin (API) 20 µM blocks IL-6, IL-8, MCP-1, and TNF-α cytokine production in human primary astrocytes (HCA). Representative flow cytometry traces from 12 independent experiments (see Extended Data Fig. 1A). B. Chrysoeriol (CHR) 20 µM suppresses LPS-induced pro-inflammatory cytokine release in human iPSC-derived microglia (n = 2, mean fold-change (FC); only cytokines with significant FC, P < 0.05, shown). Representative of 3 independent experiments. C. CHR 20 µM blocks microglial phagocytosis of E. coli-FITC conjugated beads (5-hour treatment, n = 2 for “cytoD” group, n = 3 for the other groups, mean ± SD, one-way ANOVA with Dunnett’s post-hoc test). Mean fluorescence intensity (MFI) fold-change relative to DMSO. Representative of 3 independent experiments. D. CHR exhibits dose-dependent anti-inflammatory activity in human astrocytes (top) or iMGL (bottom) by blocking IL1-β or LPS-induced IL-6 secretion, respectively (n = 9, mean ± s.e.m.). Normalized to DMSO vehicle ± stimulation; 3 independent experiments pooled for analysis. E. Thermal shift proteome profiling (TPP) in human iPSC-derived astrocytes. Red denotes proteins exhibiting significant and reproducible thermal shifts after CHR treatment in IL1-β-activated astrocytes. F. Isothermal dose response thermal shift assay validates CK2 as a target of CHR. Each dot represents normalized band intensity (Western blot in Extended Data Fig. 2A). Representative of 3 independent experiments. * P < 0.05; ** P < 0.01

### Target identification by thermal shift proteome profiling (TPP) reveals CK2 as a flavone target

Next, we undertook an unbiased proteomics approach to identify cellular targets of flavones in inflamed astrocytes. TPP identifies drug target engagement in cells by measuring ligand stabilization of bound proteins against heat denaturation^22,23^. In this assay, stabilized proteins can be either direct or indirect targets, whereas destabilized proteins are indirect targets. We extracted lysates from IL1-β-treated human iPSC-derived astrocytes, treated them with CHR, TMF or DMSO for 30 minutes, and performed TPP using LC-MS/MS with TMT10 labeling.

In the CHR vs. DMSO comparison, we identified 8 proteins with a significant thermal shift (ΔT_m_) reproducible in 2 independent experiments (Fig. 1E; Supplementary Table 1). Two of the stabilized hits, CSNK2A3 (CK2α1P) and CSNK2B (CK2β), constitute components of the CK2 holoenzyme, suggesting that the CK2 complex could be a direct target of CHR. CSNK2A1 and CSNK2A2 peptides did not pass mass spectrometry stringency filters in the TPP analysis. We confirmed that one of the alpha subunits of the CK2 holoenzyme, CK2α1, is dose-dependently stabilized by CHR in an isothermal dose response thermal shift assay followed by Western blotting (Fig. 1F, Extended Data Fig. 2A).

Network analysis of the other identified TPP hits (ADK, PTGR1, TP53BP1, CALU, RCN2, ARF3) showed high connectivity with each other and CK2 (Extended Data Fig. 2B). ARF3 showed thermal destabilization indicative of an indirect hit. ADK and TP53BP1 physically interact with CK2^24,25^, CALU^26^ is a known CK2 substrate, and RCN2 is a likely CK2 substrate^26,27^. We performed co-immunoprecipitation with CK2α1 and showed that it associates with PTGR1 (Extended Data Fig. 2C).

In the TMF vs. DMSO comparison, we found 11 proteins with a significant thermal shift reproducible in 2 independent experiments (Extended Data Fig. 2D). CSNK2A3 and CSNK2B were stabilized by TMF but not enough to pass the stringent statistical significance requirements. Network analysis of the 11 hits and all 4 CK2 subunits, however, showed high connectivity (Extended Data Fig. 2E). Among the 8 stabilized proteins, 6 are known CK2 substrates^26,28^ (RCN1, RCN3, SET and its paralogue SETSIP, CALU, AKAP12) and ACTR2 is a known interactor^24,25^. We validated that CK2α1 was indeed a target of TMF (but not inactive flavanone dTMF) in astrocytes in a thermal shift assay followed by Western blotting (Extended Data Fig. 2F).

Flavones have previously been shown to inhibit CK2 *in vitro*^29–34^ and in cancer cells^35,36^, with existing co-crystal structures of flavone derivatives and CK2 reflecting protein binding as ATP-competitive inhibitors^30,37^. Furthermore, CK2 is known to have roles in inflammation^38–42^. Overall, these data strongly suggested that CK2 is a relevant target to pursue for further validation.

### Validation of CK2 as a potent anti-inflammatory target in astrocytes

We verified that flavones directly inhibited CK2 in cells via additional orthogonal methods. In astrocytes, we found that API was able to outcompete binding of a desthiobiotin-ATP probe to both CK2α1 and CK2α2 (Extended Data Fig. 3A). We also confirmed CK2 target engagement of API (IC_50_ = 14.9 µM CK2α2) and CHR (IC_50_ = 4.2 µM CK2α2, 10.5 µM CK2α1) in transiently transfected HEK293T cells via NanoBRET^4343^ (Extended Data Fig. 3B). The NanoBRET assay works by measuring bioluminescent resonance energy transfer (BRET) between CK2α1- or CK2α2-Nanoluciferase and a tracer, which is suppressed when a CK2 inhibitor is present.

Revisiting our structure-activity studies in the context of CK2 inhibition, we found that anti-inflammatory activity in astrocytes correlated well with CK2 inhibitory activity (Fig. 2A). API, quercetin, fisetin, luteolin, kaempferol, and CHR possessed IC_50_ values ranging from 80-800 nM^30,33^ as well as strong suppressive activity against at least 3 of 4 cytokines measured (Extended Data Fig. 1A,C) and strong activity (IC_50_ < 4 µM) in THP-1 monocyte NF-κB-Luciferase reporter cells (Fig 2B). Chrysin possessed a CK2 IC_50_ value of 9 µM^30^ and showed partial CK2α1/CK2α2 inhibition in cells (Extended Data Fig. 3C), with intermediate anti-inflammatory activity exhibiting as variable suppression of 2 cytokines and no activity in THP-1 monocyte NF-κB-Luciferase reporter cells (Extended Data Fig. 1A, 3D). In contrast with their flavone analogues, flavanones missing the key C2-C3 double bond identified earlier were inactive both as CK2 inhibitors and anti-inflammatory agents (Extended Data Fig. 3C-E, NAR, dTMF, dCHR, IC_50_ > 10 µM).

**Figure 2.**
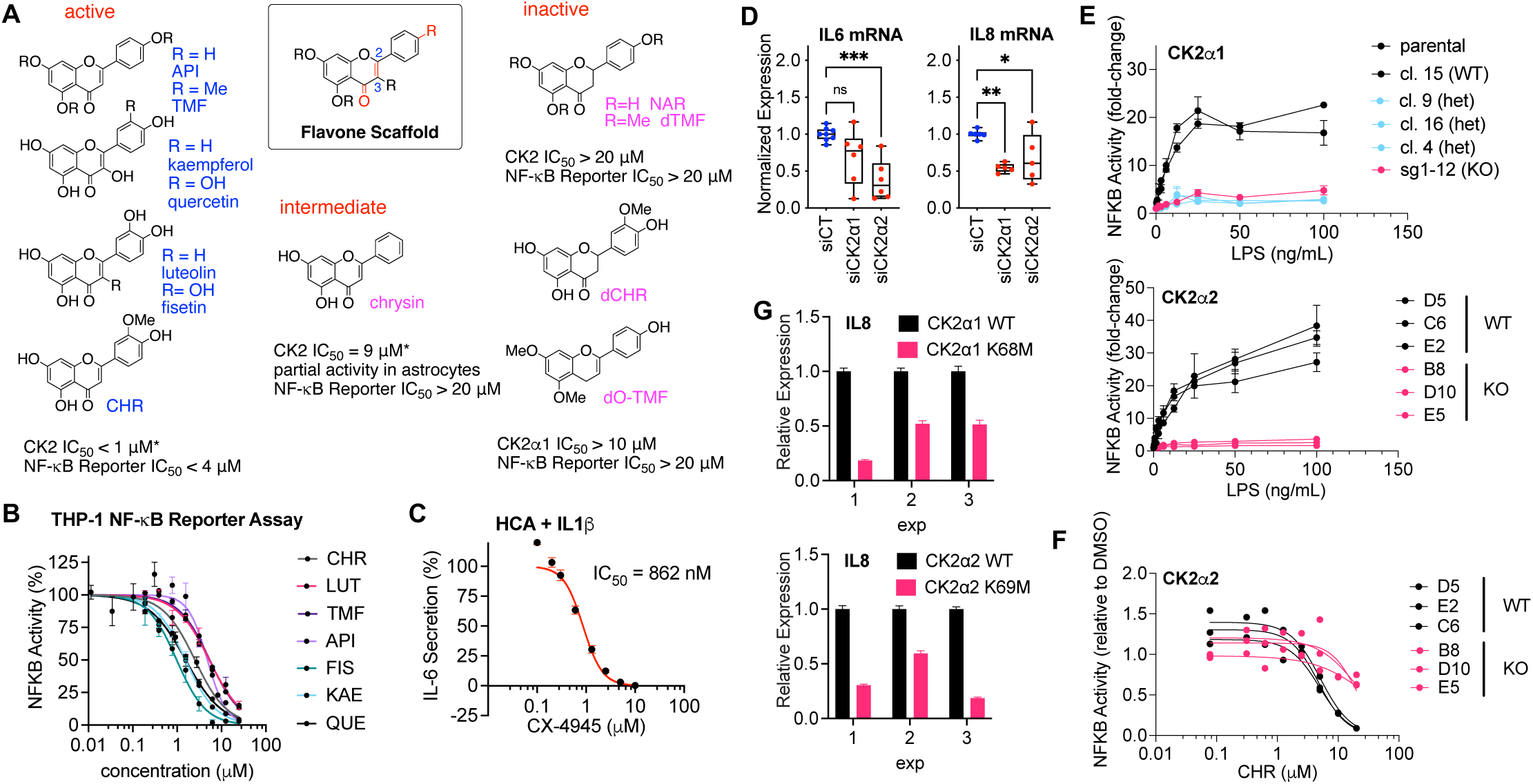
Inhibition of CK2 kinase activity reduces astrocyte inflammation and NF-κB activity. A. Summary of SAR data shows that API, CHR, luteolin, quercetin, kaempferol, fisetin and TMF are active, chrysin is intermediately active, and dTMF, dCHR, NAR, and dO-TMF are inactive at 20 µM both in an NF-κB reporter assay and for CK2 kinase inhibition. Asterisk denotes values sourced from Baier et al. (2017) and Lolli et al. (2012). Highlighted in red are the most important structural features for activity. B. Most-active flavones dose-dependently reduce NF-κB activity in THP-1 macrophages (LUT = luteolin, QUE = quercetin, FIS = fisetin, KAE = kaempferol, n = 3, mean ± s.e.m.). Representative of at least 2 independent experiments (3 for CHR, 8 for TMF, 10 for API). C. Structurally unrelated CK2 inhibitor CX-4945 blocks IL1-β-induced IL6 upregulation (mean ± s.e.m., n = 3). Normalized to DMSO, data are representative of 6 independent experiments. D. KD of CK2α1 and CK2α2 significantly blocks IL1-β-induced IL6 and IL8 upregulation. Box plot with median values and 25-75% CI of n = 8 (siCT), n = 6 (siCK2α1), n = 6 (siCK2α2) biological replicates for IL6 or n = 6 (siCT), n = 5 (siCK2α1), n = 5 (siCK2α2) biological replicates for IL8, pooled from 4 independent experiments, one-way ANOVA with Dunnett’s post-hoc test). E. Quantification of NF-κB reporter luciferase activity in CK2α1 (top) or CK2α2 (bottom) heterozygous (het) or homozygous (KO) knockout THP-1 NF-κB Lucia cell clones stimulated with various doses of LPS (mean ± sem, n = 3). Representative of 3 independent experiments. F. Quantification of NF-κB reporter luciferxase activity in CK2α2 homozygous (KO) knockout THP-1 NF-κB Lucia cell clones stimulated with 20 ng/mL LPS and CHR dilution series (n = 3 clones for each group; individual dose-response curve fits per clone shown). Representative of 3 independent experiments. G. Overexpression of CK2 kinase-dead mutants significantly blocks IL1-β-induced IL-8 upregulation in hES-derived astrocytes (3 independent experiments shown with Poisson error); each value normalized within experiments to WT. ns = not significant; * P < 0.05; ** P < 0.01; *** P < 0.001

We wondered if a structurally unrelated CK2 inhibitor could also block inflammation in astrocytes stimulated with IL1-β. Treatment with CX-4945^44^, a potent synthetic CK2 inhibitor currently in clinical trials for cancer, effectively blocked secretion of IL-6 after 5 hours (IC_50_ = 315 nM, Fig. 2C). CX-4945 also successfully blocked expression of *IL6* and *IL8* mRNA in astrocytes to a similar extent as TMF (Extended Data Fig. 3F). CX-4945 and CK2 inhibitor TBB have been previously shown to reduce secretion of IL-6 and MCP1 in primary astrocytes^45^. Recently, Mishra et al. also reported that CX-4945 and SGC-CK2-1, a selective chemical probe for CK2, blocked IL-6 and IL-8 expression in iPSC-derived microglia^46^.

We asked whether genetic perturbation of CK2 could phenocopy the effects of pharmacological inhibition that we observed with flavones. We performed siRNA knockdown of individual CK2 catalytic subunit isoforms CK2α1 or CK2α2 in human primary astrocytes. After 3 days, cells were treated with IL-1β for 5 hours and RNA was extracted to assess cytokine levels. We observed that knockdown of either CK2α1 or CK2α2 reduced the ability of IL-1β to induce upregulation of pro-inflammatory cytokines *IL6* and *IL8* (Fig. 2D, Extended Data Fig. 4A).

To further dissect the individual roles of the 2 catalytic forms, we performed CRISPR-Cas9-mediated knockout of *CSNK2Α1* or *CSNK2Α2* in THP-1 monocyte NF-κB-Luciferase reporter cells and isolated clonal lines to measure their response to inflammatory stimulation and anti-inflammatory activity of CHR. Isolating homozygous CK2α1 clones proved difficult, highlighting the importance of this isoform for THP-1 cancer cell survival.^47^ Overall, we observed that knockout of CK2α1 or CK2α2 reduced the ability of IL1-β to induce inflammation in these cells (Fig. 2E and Extended Data Fig. 4B-C). However, only CK2α2 knockouts became significantly resistant to the anti-inflammatory effects of CHR, highlighting CK2α2 as the primary driver of inflammation in this model system (Fig. 2F and Extended Data Fig. 4D).

Correspondingly, we wondered if overexpression of a kinase-dead form of CK2 could affect inflammatory induction in astrocytes. We transduced glial progenitor cells (GPCs) with an LV-CK2α1 or CK2α2 wild-type (WT) or single point mutant (K68M and K69M, respectively)^48^, differentiated them for 4 weeks to produce mature astrocytes, and then stimulated them with IL1-β for 5 hours. We found that CK2α1-K68M (P = 0.033) or CK2α2-K69M (P = 0.034) significantly abrogated inflammation relative to WT (paired t-tests, Fig. 2G, Extended Data Fig. 4E-F).

Overall, these data strongly suggest that CK2 is the relevant cellular target of anti-inflammatory flavones and that it is an important upstream pro-inflammatory regulator in astrocytes.

### CK2 activates NF-κB in response to inflammatory stimuli in astrocytes

Others have reported that CK2 protein levels increase in macrophages and certain cancer cell lines in response to pro-inflammatory stimuli like LPS and TNF-α^38,40^. We observed a similar increase in CK2 levels in human primary astrocytes stimulated with IL1-β, with CK2α1 and CK2α2 peaking at 1 hour after induction and returning to baseline after 5 hours (Fig. 3A). Astrocytes co-treated with CK2 inhibitors (CHR or CX-4945) and IL1-β exhibited a compensatory elevation of CK2α1 and CK2α2 levels and a slower return to baseline. At this time point, we also observed CHR- and CX-4945-mediated reduction of CK2 pY255 (a marker of CK2 activation^49^) relative to total CK2α1 in astrocytes by immunofluorescence (Fig. 3B), which indicates that CK2 is an immune-responsive kinase in astrocytes.

**Figure 3.**
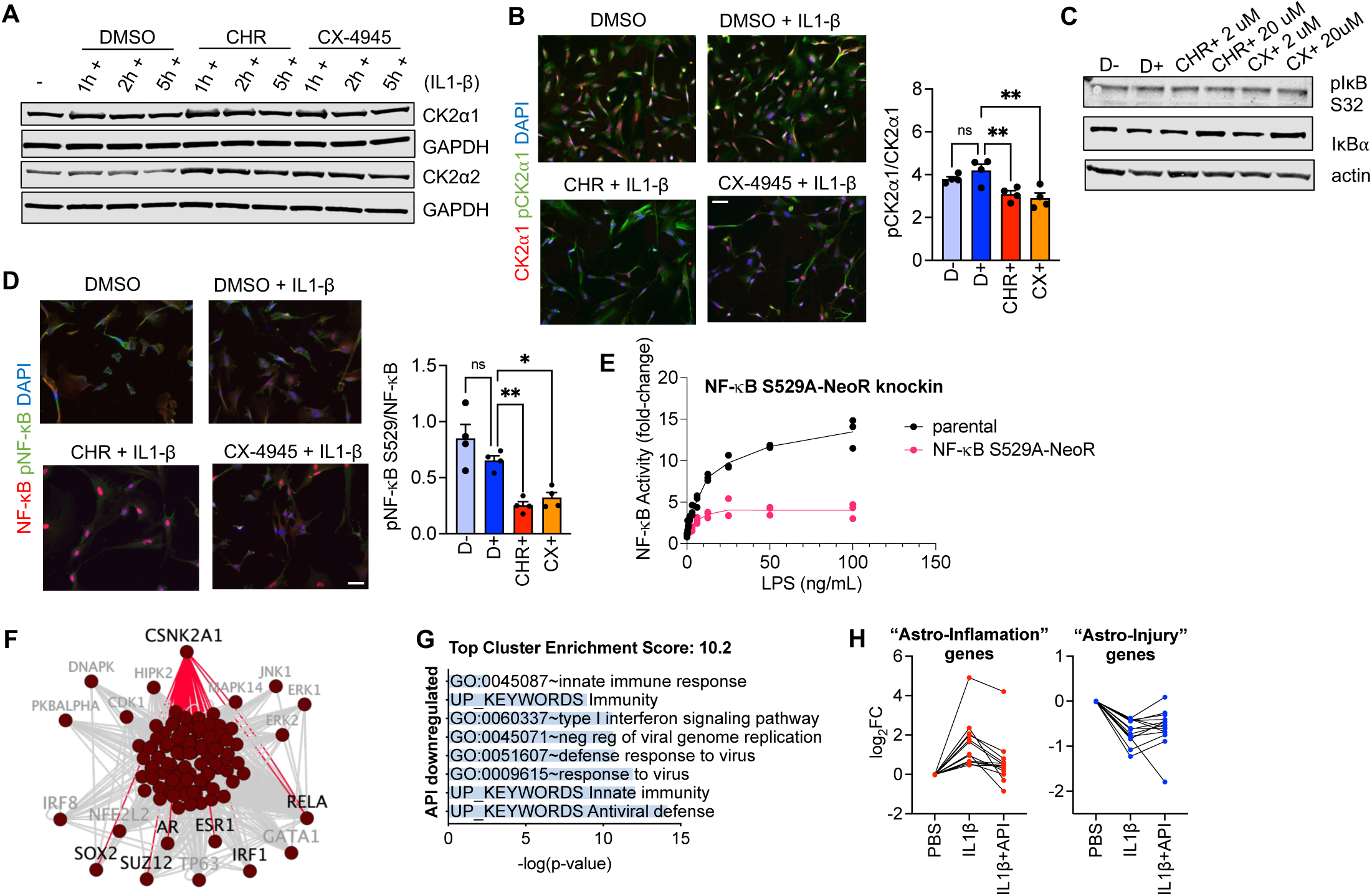
CK2 inhibition attenuates inflammatory biochemical and transcriptional programs via NF-κB. A. CK2 levels increase with inflammation (time course by Western blot). Full blots in Extended Data Figure 15. B. Representative immunofluorescence images and quantification showing reduction of nuclear phospho-CK2α1 Y255/CK2α1 after 5 hours of CK2 inhibitor treatment (mean ± s.e.m., n = 4, one-way ANOVA with Dunnett’s post-hoc test). Scale bar = 100 µm. C. Representative immunoblots showing that pIκB/IκBα levels are reduced with CK2 inhibitors (6-hour treatment). Full blots in Extended Data Figure 15. D. Representative immunofluorescence images and quantification showing reduction of nuclear phospho-NF-κB S529/NF-κB after 5 hours of CK2 inhibitor treatment (mean ± s.e.m., n = 4, one-way ANOVA with Dunnett’s post-hoc test). Scale bar = 100 µm. E. Quantification of NF-κB reporter luciferase activity in a pool of phosphodeficient THP-1 NF-κB Lucia NF-κB-S529A-NeoR knockin cells or parental WT stimulated with various doses of LPS (mean of n = 3 biological replicates shown). F. X2K interaction network showing top enriched kinase modules, intermediate proteins (not labeled), and their downstream TF targets in IL1-β-stimulated astrocytes (gray nodes and edges unrelated to CK2, black nodes connected to CK2 via red edges). G. Immune signatures are significantly enriched in genes downregulated in inflamed astrocytes treated with API. H. CK2 inhibition reduces expression of “Astro-Inflammation” genes and increases expression of “Astro-Injury” genes. A-E: Representative of 3 independent experiments unless otherwise noted. ns = not significant; * P < 0.05; ** P < 0.01

Several proteins involved in the innate immune response have been characterized as substrates of CK2, including IκB S32^50^ and NF-κB S529^51^. NF-κB is a master transcriptional regulator of the immune response, downstream of a broad repertoire of exogenous stimuli, including Toll-like receptors, IL-1R, and TNFR. IκBα is a negative feedback regulator and acts as an inhibitor of the NF-κB program by sequestering NF-κB in the cytoplasm. IκBα S32 phosphorylation is necessary for degradation of IκBα by pro-inflammatory stimuli.^50^ IκBα was downregulated in inflamed astrocytes after 6 hours, but CK2 inhibitors CHR and CX-4945 dose-dependently boosted IκBα levels whereas phosphorylated IκB S32 remained unchanged (Fig. 3C, Extended Data Fig. 4G). Overall, these findings are consistent with CK2 acting downstream of IL1-β to phosphorylate IκB S32, with CK2 inhibition leading to a reduction in pIκB and subsequent stabilization of total IκBα.

To investigate the potential effects of CK2 activity on NF-κB, we measured NF-κB and NF-κB pS529 by immunofluorescence in activated astrocytes. After 5 hours, we found that both CHR and CX-4945 reduced pNF-κB nuclear intensity relative to total NF-κB (Fig. 3D). At the intermediate time point of 2 hours post-induction with IL1-β, we detected an increase in pNF-κB relative to NF-κB via immunoprecipitation (IP) followed by immunoblotting; this increase in phosphorylated pNF-κB species was abrogated by CHR co-treatment (Extended Data Fig. 4H). To further study the effects of CK2-dependent NF-κB S529 phosphorylation, we knocked in phosphodeficient mutation S529A along with the neomycin resistance gene into the endogenous NF-κB locus in THP-1 NF-κB-Luciferase cells. We found that the S529A mutation diminished NF-κB activation in response to LPS treatment, phenocopying CK2 inhibition, knockdown, or knockout (Fig. 3E).

These findings correlate the upregulation of CK2 catalytic subunits (Fig. 3A) and the anti-inflammatory activity of CK2 inhibition observed at this time point (Fig. 1D, 2C) with NF-κB activation. Overall, these results reflect the dynamic regulation of NF-κB by CK2 and indicate that CK2 might play an important role in sustaining NF-κB-driven immune response.

### CK2 kinase activity sustains an inflammatory transcriptional program

To further investigate the mechanistic role of CK2 in driving inflammation in glia, we performed RNA sequencing (RNA-seq) in human primary astrocytes stimulated with IL1-β and API or vehicle for 5 hours. Differential expression (DE) analysis showed 533 genes downregulated and 938 genes upregulated upon activation (P < 0.05, FC > 2, Supplementary Table 2). This inflammatory signature was strongly enriched in genes regulated by CK2, as identified by Kinase Enrichment Analysis^52^ (KEA, P = 5.8×10^-16^ and P = 1.19×10^-10^, Extended Data Fig. 5A). Indeed, network analysis showed that CK2 played a central role in regulating many of the downstream enriched transcription factor (TF) modules, including RELA/NF-κB, IRF1, and polycomb repressive complex subunit SUZ12 (Fig. 3F, Extended Data Fig. 5B). IRF1 is another key TF of innate immunity^53^ and SUZ12/EZH2 have been shown to regulate NF-κB target gene expression^54^.

Next, we compared gene expression of cells treated with API + IL1-β versus IL1-β alone. We found that 531 genes were upregulated and 1,603 genes were downregulated (P < 0.05, |log2FC| > 1, Supplementary Table 2). Notably, API reversed a significant fraction of the gene expression changes induced in activated astrocytes (58% of the genes upregulated by IL1-β, P = 10^-1037^, Extended Data Fig. 5C; and 40% of the genes downregulated by IL1-β, P = 2.7×10^-226^, Extended Data Fig. 5D). Among all the API-downregulated genes, the top cluster of enriched terms by functional gene annotation analysis was related to the innate immune response (Fig. 3G). Querying DE genes following API treatment in the CMap database^55^, which catalogs transcriptional responses of human cells to chemical and genetic perturbations, we found that multiple CK2 inhibitor and CK2 knockdown signatures exhibited high similarity scores (Extended Data Fig. 5E). In contrast, 9 of the top 20 most dissimilar signatures in the CMap database were related to NF-κB activation. These results indicate that CK2 inhibition effectively attenuates the inflammatory response in astrocytes.

We next examined whether CK2 inhibition modulated the polarization of reactive astrocytes. We observed how API modulated the expression of genes related to an astrocyte state induced by inflammation (“Astro-Inflammation”) versus an astrocyte state induced by an acute injury (“Astro-Injury”) like cerebral ischemia^56,57^ (Supplementary Table 3). Of the 14 “Astro-Inflammation” genes significantly upregulated by IL1-β in primary astrocytes, 10 were significantly downregulated by API, including interferon alpha/beta signaling molecules IFI44, GBP2, and PARP14 (P < 0.05, two-tailed t-tests, Fig. 3H). Type I interferon signaling has been linked to neurotoxic effects under chronic inflammatory conditions^58^. Similarly, of the 13 “Astro-Injury” genes downregulated by IL1-β, 12 were upregulated by API, including neuroprotective factor BNDF and synaptogenic factor THBS1 (P < 0.05, two-tailed t-tests, Fig. 3H).

Taken together, these data provide evidence that CK2 is a potent regulator of inflammation in glia upstream of NF-κB. At the transcriptional level, CK2 maintains the polarization of reactive astrocytes towards pro-inflammatory pathways at the expense of an acute injury-like response. CK2 inhibition suppressed the inflamed astrocyte phenotype (in part, marked by expression of “Astro-Inflammation” genes) and promoted an acute injury-responsive astrocyte state (in part, marked by expression of “Astro-Injury” genes).

### CK2 activity is upregulated in AD astrocytes and contributes to a hyperinflammatory astrocyte phenotype

Having established that CK2 regulates neuroinflammation in human glia, we asked whether neurodegenerative diseases such as AD exhibited dysregulated CK2 activity, as astrogliosis and chronic inflammation are linked to pathogenesis in AD. There is some evidence of dysregulation of CK2 in neurodegenerative diseases. Higher CK2 levels were observed in astrocytes in the cortex and hippocampus of AD patients, with localization near amyloid plaques^45^. Marshall et al. recently reported that CK2 levels in AD hippocampi correlated inversely with MMSE scores, a marker of cognitive decline.^59^

We wondered if CK2 dysregulation could be observed in human AD models. Using a cohort of 7 AD and 4 cognitively normal age-matched control iPSC cell lines (Fig. 4A, Supplementary Table 4), we generated patient-derived astrocytes and asked if these observations could be recapitulated *in vitro* using this model (Extended Data Fig. 6A). RNA-seq followed by gene ontology analysis of significantly AD-upregulated genes using ClueGO and DAVID showed evidence of baseline inflammatory activation in AD astrocytes, with enriched terms including secreted proteins, innate immunity, and interferon signaling (Extended Data Fig. 7A-B; Supplementary Table 5). *CSNK2A2* was significantly upregulated in AD patient-derived astrocytes compared to control (Fig. 4B). We verified that CK2α2 protein levels were also significantly and specifically elevated in AD patient-derived astrocytes, and this elevation was associated with downstream elevated phosphorylated NF-κB that could be reduced with CHR treatment (Fig. 4C-D, Extended Data Fig. 6B-C). Transcriptionally, KEA showed that all 3 CK2 subunits were among the top 20 enriched kinase hubs regulating AD-upregulated genes (Fig. 4E). These data support a role for CK2, especially isoform CK2α2, in driving a chronic inflammatory transcriptional program in AD astrocytes.

**Figure 4.**
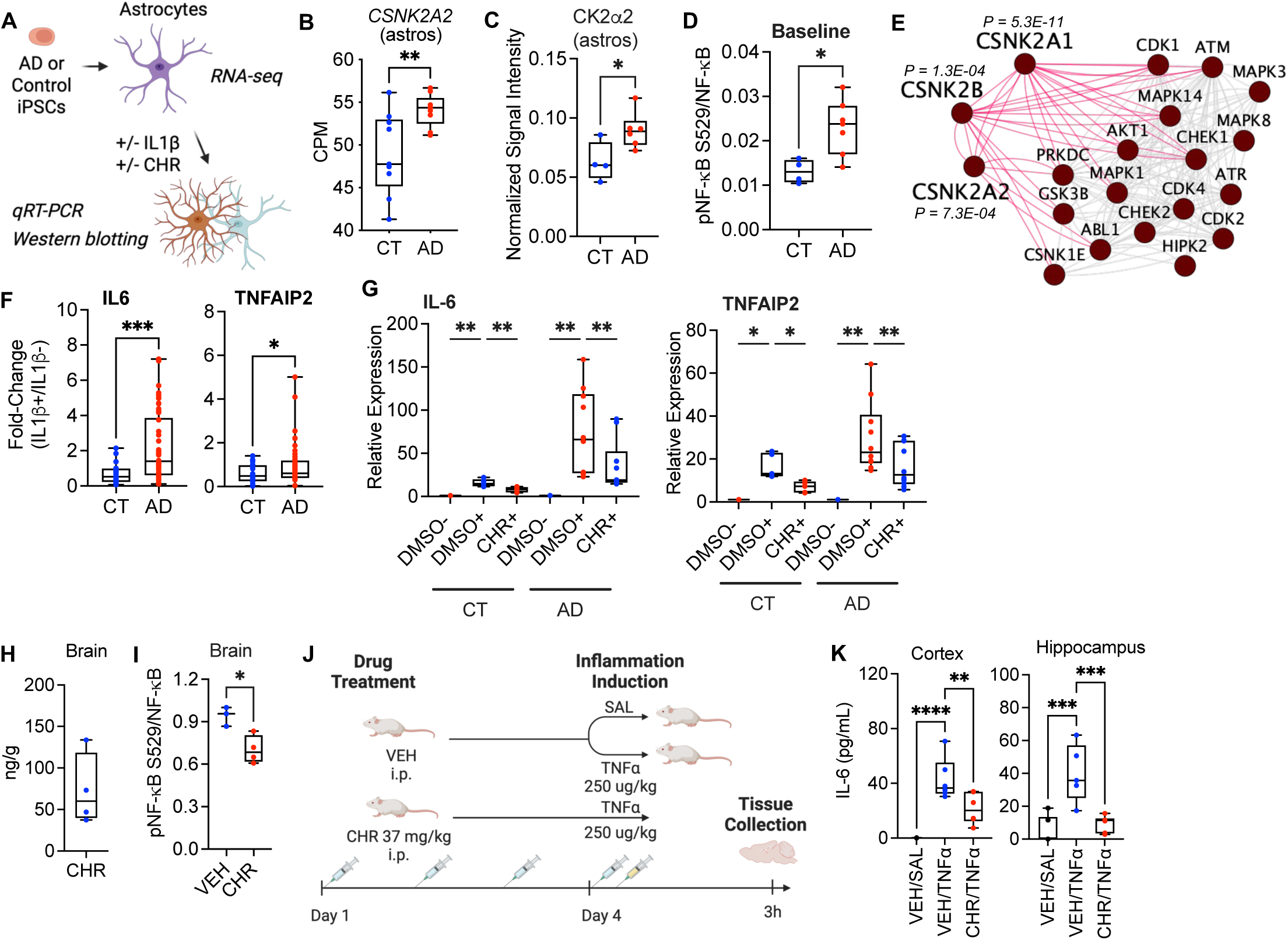
CK2 is dysregulated in Alzheimer’s disease (AD) patients and models, and CK2 inhibition ameliorates AD-associated neuroinflammation *in vitro* and TNF-a-induced neuroinflammation *in vivo*. A. Overview of AD and control astrocyte differentiations and associated studies performed in panels B-G. B. Box plots showing counts per million (CPM) CSNK2A2 mRNA expression by RNA-seq (n = 3 AD and n = 3 CT samples in triplicate; two-tailed t-test). C. Quantification showing cellular CK2α1 and CK2α2 protein levels normalized to GAPDH in AD vs. control iPSC-derived astrocytes (n = 6 AD and n = 4 CT samples in duplicate, two-tailed t-test, P = 0.027). Representative immunoblots shown in Extended Data Fig. 6B. D. Quantification of pNF-κB/NF-κB baseline levels (n = 7 AD and n = 4 CT samples; t-test). Immunoblots shown in Extended Data Fig. 6C. E. STRING network of top 20 significantly enriched kinases by KEA of significantly AD-upregulated genes (p-values shown for CK2 isoforms). F. Box plots showing IL-6 and TNFΑIP2 fold-change (two sets of n = 5 and n = 6 AD and n = 3 and n = 4 CT cell lines in duplicate; two-tailed t-test). Pool of 4 independent experiments. G. Box plots showing IL-6 and TNFΑIP2 mRNA expression relative to GAPDH (n = 5 AD and n = 3 CT samples in duplicate; mixed-effects ANOVA with Sidak’s post-hoc test). H, I. Box plots showing CHR levels measured by LC-MS/MS (I) and pNF-κB/NF-κB levels (J) in mouse brain homogenate 1 hour following single i.p. injection of 40 mg/kg CHR (n = 4 mice) or VEH (n = 3 mice). J. Schematic of *in vivo* experimental design for acute neuroinflammation efficacy study. K. Box plots showing IL-6 cytokine expression in the cortex and hippocampus in mice pre-treated with 37 mg/kg CHR or VEH daily for 4 days prior to exposure to an acute systemic dose of TNFα or saline (SAL) for 2 hours (n = 6 mice per group, one-way ANOVA with adjusted p-values via Benjamini, Krieger, Yekutieli FDR method). C-D, F-G: Representative of 3 independent experiments unless otherwise noted. All box plots show median and 25-75% CI. Panels A and J created with Biorender (https://BioRender.com/xcrzfm6, https://BioRender.com/nsluzrv). * P < 0.05; ** P < 0.01; *** P < 0.001; **** P < 0.0001

We next studied whether AD astrocytes could have a dysregulated response to inflammatory stimuli in addition to their baseline inflamed state. Indeed, AD astrocytes exhibited a hyperinflammatory response to IL1-β treatment, as demonstrated by the higher magnitude of IL-6 and TNFΑIP2 induction compared to controls (Fig. 4F). This hyperinflammatory phenotype was mitigated by CK2 inhibition with CHR, which also reduced NF-κB phosphorylation (Fig. 4G; Extended Data Fig. 6C).

Taken together, these data indicate that CK2 activity is upregulated in astrocytes derived from patients with neurodegenerative diseases like AD and that AD astrocytes exhibit a hyperinflammatory state due in part to CK2 activation. In AD patient-derived astrocytes, CK2 levels correlated with higher induced secretion of inflammatory cytokines like IL-6 that could be abrogated via small molecule CK2 inhibition.

Therefore, we sought to test whether CK2 inhibition could block neuroinflammation *in vivo* using an acute TNFα systemic administration model^60^. Since natural product flavonoids are known to have poor bioavailability and limited drug-like properties, we hypothesized that CHR would likely be cleared rapidly and have trouble crossing the blood-brain-barrier due to the presence of multiple hydrogen bond donors. Given these limitations, we chose to administer the compound intraperitoneally to avoid first-pass metabolism in the gut. One hour after a single-dose intraperitoneal (i.p.) injection at a dose of 40 mg/kg, CHR was detected in mouse brain in appreciable quantities, engaging CK2 and reducing NF-κB phosphorylation (Fig 4H,I, Extended Data Fig. 8A). For the acute neuroinflammation challenge, we chose to administer the compound in multiple doses to allow build-up in the brain before stimulation (Fig. 4J). We treated mice daily for 3 days with 37 mg/kg CHR i.p.; 1 hour after the last dose, mice were injected i.p. with 250 µg/kg recombinant mouse TNFα and sacrificed to assess pro-inflammatory cytokine levels 3 hours later. We detected reduced levels of IL-6, MCP-1, IP10, KC, and TNFα coupled with downregulation of TNFα-induced IκB S32 phosphorylation downstream of CK2 in the cortex and hippocampus (Fig 4K; Extended Data Fig. 8B-C). Overall, these results reflect the potential of CK2 small molecule inhibitors to rescue neuroinflammatory phenotypes *in vivo*.

### CK2 activity is upregulated in AD postmortem samples, and overexpression of CK2α2 induces AD-like phenotypes in glia-enriched cortical organoids

To assess the *in vivo* translatability of our AD astrocyte *in vitro* model, we replicated our findings in postmortem brain tissue from AD patients and cognitively normal controls. First, we analyzed transcriptome profiling data from the Aging, Dementia, and TBI RNA-seq study^61,62^ and found that *CSNK2A2,* but not *CSNK2A1* or *CSNK2B*, was significantly upregulated in dementia patients compared to healthy controls (P = 0.024, two-tailed t-test, Extended Data Fig. 9A). We then analyzed protein expression data from the UPP Proteomics Study^63^ and confirmed that CK2α2 protein was also upregulated in AD patients compared to controls, whereas CK2α1 and CK2β were expressed at similar levels (P = 0.047, one-way ANOVA with Sidak’s post-hoc test, Extended Data Fig. 9B). Interestingly, CK2α2 expression is enriched in the brain relative to other tissues, unlike CK2α1, which is uniformly expressed throughout tissues^64^. CK2α2 was significantly elevated relative to CK2α1 (P < 0.0001) and CK2β (P = 0.0056) in AD patients but not in controls (one-way ANOVA with Sidak’s post-hoc test, Extended Data Fig. 9B).

Given these results, we obtained fresh frozen postmortem brain tissues for analysis of CK2 enzymatic activity and CK2 expression (n = 10 per group, Supplementary Table 6) and fixed brain tissues for analysis of cell type-specific CK2 isoform expression (n = 10 per group, Supplementary Table 7). CK2α1 levels appeared to be lower in AD patients by immunoblotting, but total protein staining by Amido Black showed that the AD samples had less protein overall despite similar loading amounts, rendering this observation a likely artifact of higher protein degradation in the AD samples (Extended Data Fig. 9C). Indeed, a multiple banding pattern for CK2α1 can be observed in the AD patient lanes, indicating degradation. CK2α2 levels appeared similar across AD and cognitively normal sex- and age-matched controls. To account for these differences in protein integrity, we quantified CK2 activity normalized to total CK2 expression and found that it was significantly elevated in AD patients versus controls (n = 10 per group, P = 0.026, two-tailed t-test, Fig. 5A, Extended Data Fig. 9C).

**Figure 5.**
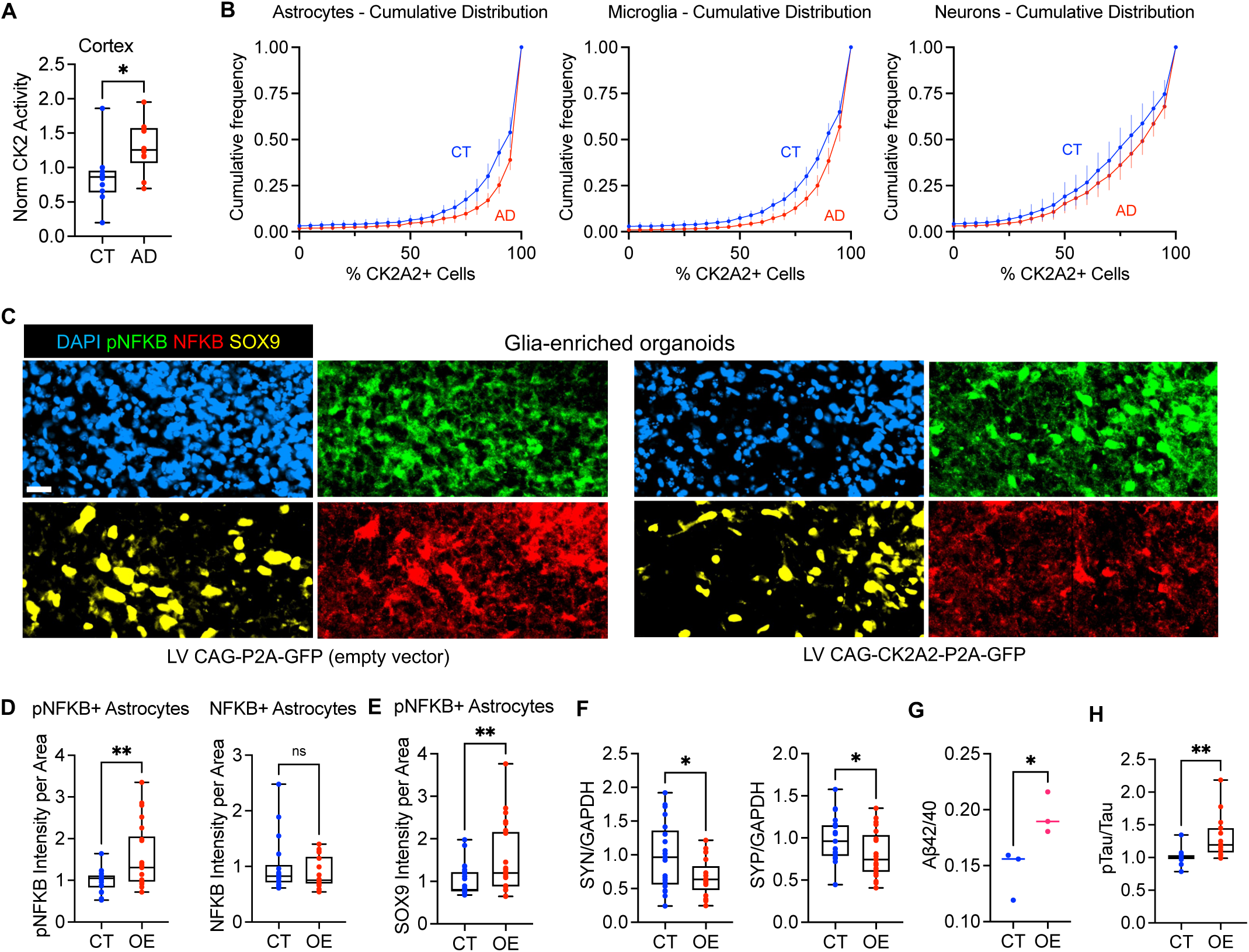
Brain-enriched isoform CK2α2 is specifically elevated in AD postmortem tissues and its activity causes upregulation of inflammatory and synaptic dysfunction markers in a glial organoid model. A. Box plots showing CK2 enzymatic activity in postmortem cortical brain AD or control samples from the NIH Neurobiobank normalized to total CK2 protein levels (n = 10 individuals per group, two-tailed t-test). Representative of 2 independent experiments. See Extended Data Fig. 9C for accompanying blots. B. Cumulative distribution plots of the percent CK2A2 positivity of each respective cell population across regions in control or AD groups in postmortem cortical brain AD or control samples from the UCSD ADRC as measured by immunofluorescence staining (mean ± sem from n = 9 CT individuals, n = 10 AD individuals). See Extended Data Fig. 10A-D for associated analysis, representative images and similar analyses of CK2α1 and CK2β. C. Representative immunofluorescence images showing pNF-κB and NF-κB expression of astrocytes (SOX9), counterstained with DAPI, from glia-enriched organoids transduced with LV CAG-P2A-GFP (empty vector) or LV CAG-CK2A2-P2A-GFP (CK2A2 overexpression). For Figures 6D-H, CT = empty vector and OE = CK2α2 overexpression. Scale bar = 20 µm. D. Quantification of pNF-κB intensity per area of pNF-κB+ astrocytes (left) and NF-κB intensity per area of NF-κB+ astrocytes (right); related to Figure 5C. Median of pooled data from 2 independent experiments shown (n = 18 CT and n = 22 OE organoids; two-tailed t-test). E. Quantification of SOX9 intensity per area of NF-κB+ astrocytes; related to Figure 5C. Median of pooled data from 2 independent experiments shown (n = 24 CT and n = 25 OE organoids; two-tailed t-test). F. Box plots showing quantification of expression of synaptic proteins SYN and SYP normalized to GAPDH expression by immunoblotting (data pooled from 3 independent experiments, n = 21 organoids per group; two-tailed t-test). See Extended Data Figure 11F for representative immunoblots. G. Amyloid-B peptide 42/40 ratio of media collected from CT or OE organoids at 4 weeks post-infection (median, data pooled from 3 independent experiments; two-tailed t-test). H. Box plots showing pTau/Tau expression by ELISA (data pooled from 2 independent experiments, n = 17 organoids per group; two-tailed t-test). All box plots show median and 25-75% CI. * P < 0.05; ** P < 0.01; **** P < 0.0001

We next performed immunofluorescence staining for CK2 isoform expression in frontal cortex slices of AD patients and cognitively normal sex- and age-matched controls. To account for the observed heterogeneity of staining patterns across regions within each individual, we compared the mean cumulative distributions of CK2α2+ cells. We found that the distributions of CK2α2+ astrocytes and microglia in AD patient cortical tissue were right-shifted relative to controls (Fig. 5B; Extended Data Fig. 10A). Supporting this observation, 79% of cortical regions in AD patients exhibited >90% CK2α2 positivity in astrocytes, compared to 63% of regions in control individuals (n = 10 per group, P_adj_ = 0.0024, t-test adjusted for multiple comparisons, Extended Data Fig. 10B). In microglia, 68% of regions in AD patients exhibited >90% CK2α2 positivity, compared to 52% of regions in control individuals (n = 10 per group, P_adj_ = 0.0016, t-test adjusted for multiple comparisons, Extended Data Fig. 10B). In neurons, similar analyses did not show significant differences between AD and controls in the mean cumulative distribution of percent CK2α2+ cells or subject-level cumulative frequencies of CK2α2 positivity in cortical regions (Fig. 5B; Extended Data Fig. 10B). We also compared mean values of CK2α2+ cells and found that AD patients expressed more CK2α2+ microglia (P = 0.043, two-tailed t-test, Extended Data Fig. 10D); similar analyses were not significant in astrocytes or neurons. Similar analyses for CK2α1 and CK2β did not show significant differences (Extended Data Fig. 10A,C,D). Overall, CK2α2 showed high expression across cell types; in contrast, CK2α1 was moderately expressed in microglia and neurons and highly expressed in astrocytes, whereas CK2β had low expression across cell types (Extended Data Fig. 10A-B). Taken together, these data extend our *in vitro* findings, showing that CK2α2 shows dysregulated expression in astrocytes and microglia.

CK2α1 overexpression has been previously shown to cause synaptic dysfunction and cognitive decline in WT mice^65^. To investigate the potential for CK2α2 causality of neurodegenerative pathology, we employed CK2α2 overexpression in a glia-enriched cortical organoid model (GEO) that primarily contains neurons and astrocytes^66^. We transduced 6- to 9-month-old WT GEOs with an LV encoding CK2α2 and GFP or GFP alone; following 1 month, we collected organoids for immunofluorescence staining or protein lysate preparation (Extended Data Fig. 11A-B). Astrocytes and neurons exhibited similar levels of GFP expression across groups (Extended Data Fig. 11C). GEOs exhibited approximately 1.2-fold CK2α2 overexpression (Extended Data Fig. 11D-E), similar to mean fold-changes in expression of CK2α2 observed in AD patient postmortem brains (e.g. Extended Data Fig. 9B). This low level of overexpression was sufficient to induce inflammatory-like changes in astrocytes, including increased NF-κB S529 phosphorylation and SOX9 expression, which are markers that are observed in activated astrocytes^67^ (P = 0.0051, two-tailed t-test, Fig. 5C-E; total NF-κB levels unaffected). We observed significant loss of the synaptic proteins synaptotagmin (SYN, P = 0.016) and synaptophysin (SYP, P = 0.039, two-tailed t-tests, Fig. 5F, Extended Data Fig. 11F), mirroring previously reported findings with CK2α1 overexpression in WT mice^65^. Furthermore, CK2α2 overexpression caused an increase in secreted amyloid β 42/40 ratios (P = 0.039, two-tailed t-test, Fig. 5G) and tau phosphorylation (P = 0.0011, two-tailed t-test, Fig. 5H) in GEOs; both phenotypes are canonically associated with AD. Taken together, these data support the idea that CK2α2 overexpression is causally linked to AD-like phenotypic changes in astrocytes and neurons.

### Validation of chemical probe TAL606 and proof-of-concept efficacy in an AD *in vivo* model

In parallel to this work, we recently published the discovery of a series of potent and selective synthetic CK2 inhibitors with superior drug-like properties compared to natural flavones^68^. Two compounds, TAL177 and TAL606, initially emerged as viable chemical probes due to their single- to double-digit nanomolar potency for CK2α1 and CK2α2 *in vitro*, 5- to 20-fold selectivity for CK2α2 relative to CK2α1 in cells, and promising ADME properties (Fig 6A; Extended Data Fig. 12A-B), all superior to CHR^68^ (Extended Data Fig. 3B). Both compounds showed moderate permeability and low efflux in the MDR1-MDCKII model, with high levels of brain and plasma binding. Importantly, TAL177 and TAL606 demonstrated high kinome selectivity in the DiscoverX scanMAX kinase assay panel, which profiles inhibitory activity against 468 targets in a competitive setting^68^. In cells, the compounds showed dose-dependent anti-inflammatory activity against IL-1β-induced IL-6 and TNF-α secretion in astrocytes, with similar IC_50_ values to the selective CK2 inhibitor SGC-CK2-1 (Extended Data Fig. 12B).

**Figure 6.**
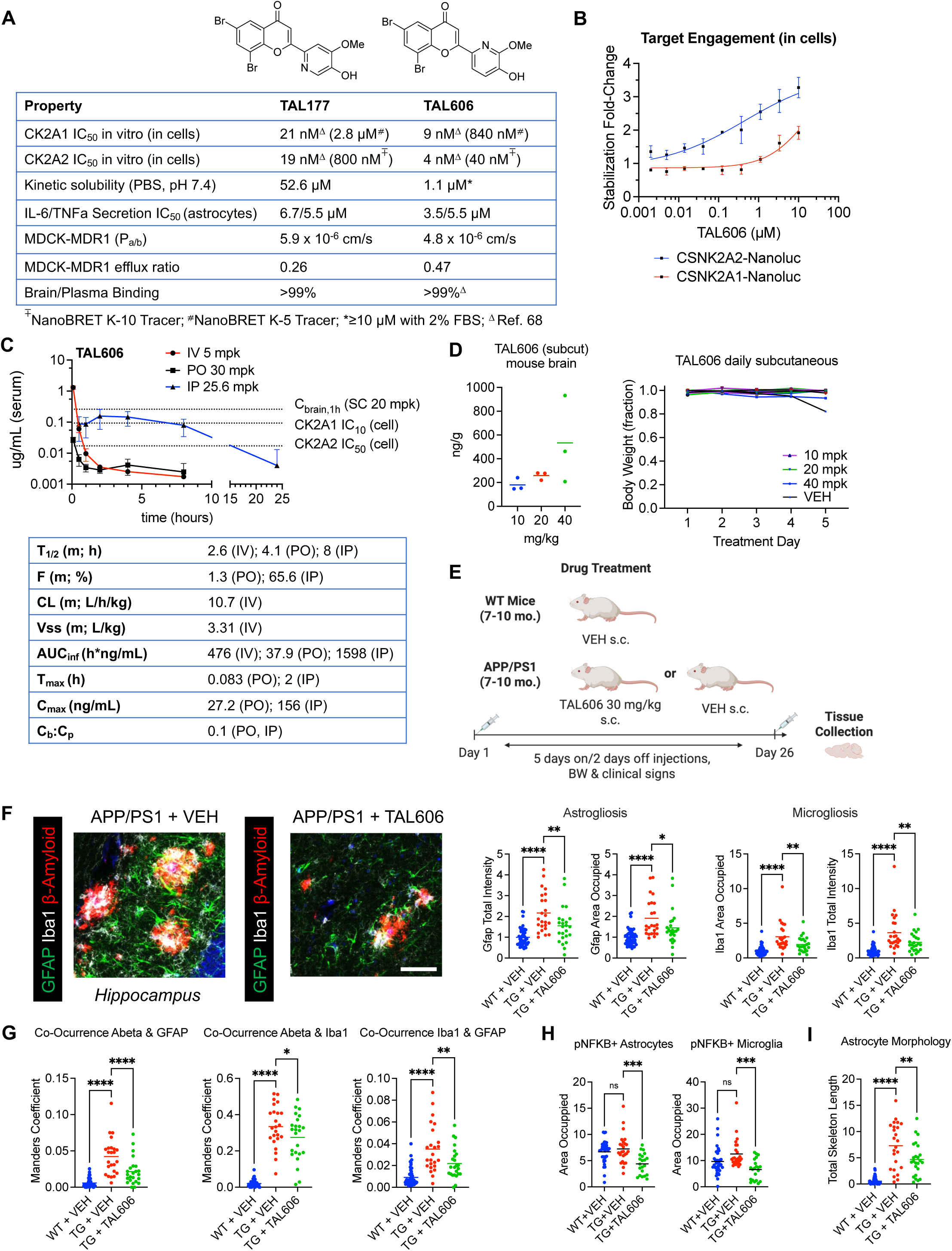
Novel flavonoid-based synthetic chemical probes TAL177 and TAL606 exhibit improved potency, selectivity, and pharmacokinetic properties as CK2 inhibitors *in vitro* and *in vivo*. A. Table showing biochemical^68^ and cellular potency, solubility, cellular anti-inflammatory activity, MDCK-MDR1 permeability, and brain/plasma binding^68^ of synthetic probes TAL177 and TAL606. See Extended Data Figures 12A-B for full dose curves. B. TAL606 shows target engagement in HEK293T cells via isothermal dose-dependent stabilization of CSNK2A1- and CSNK2A2-Nanoluc constructs (n = 3, mean ± s.e.m.). Representative of 3 independent experiments. C. Pharmacokinetic curves and parameters for IV, PO, and IP dosing of TAL606 in mice (n = 3 for IV and PO, n= 6 for IP, mean ± s.e.m.; non-compartment model). D. (Left) TAL606 concentration in mouse brains 1 hour following the fifth dose of a 5-day dosing regimen (mean, n =3). (Right) Individual body weights over time as fraction of starting weight for mice treated during the study. E. Overview of TAL606 proof-of-concept *in vivo* efficacy study in AD APP/PS1 model. Panels F-G: N=8 mice per group TG + VEH and TG + TAL606, N=18 mice WT + VEH, pooled from 2 independent cohorts; for each mouse, 3 total images of the CA1, CA3, and DG regions of the hippocampus were quantified. F. (Left) Representative immunofluorescence images and (Right) quantification of astrogliosis as reflected by GFAP area occupied (P = 0.031) and total intensity (P = 0.0027) and microgliosis as reflected by Iba1 area occupied (P = 0.0035) and total intensity (P = 0.0038) within hippocampus ROIs, normalized to volume of the image stack, with data in each cohort normalized to their WT group average. Scale bar = 50 µm. G. Co-occurrence of amyloid-beta with astrocytes (GFAP, P < 0.0001) or microglia (Iba1, P = 0.031) or astrocytes and microglia with each other (P = 0.0025) within hippocampus ROIs, quantified as Manders coefficients. H. Target engagement was determined by quantifying area occupied by p NF-κB+ astrocytes (GFAP+) or microglia (Iba1+) within hippocampus ROIs (N=8 mice per group TG + VEH and TG + TAL606, N=13 mice WT + VEH; for each mouse, 2-3 total images of the CA1, CA3, and DG regions of the hippocampus were quantified). I. Astrocyte morphology (total skeleton length) was analyzed from GFAP binary images within cortex ROIs (N=8 mice TG + VEH, N=7 mice TG + TAL606, N=18 mice WT + VEH, pooled from 2 independent cohorts; for each mouse, 3 total images of cortex were quantified). Panels F-I: one-way ANOVA with Sidak’s multiple comparisons. Panel E created with Biorender (https://BioRender.com/y7y5uxz). * P < 0.05; ** P < 0.01; *** P < 0.001; **** P < 0.0001.

To test if increased affinity of TAL606 for CK2α2 in cells could be observed in an orthogonal assay, we measured the stabilization of CSNK2A1-Nanoluc or CSNK2A2-Nanoluc in isothermal dose-response assays. We found that the CSNK2A2-Nanoluc construct showed greater stabilization compared to the CSNK2A1-Nanoluc construct in HEK293T cells (fold-change relative to DMSO, Fig. 6B). A possible explanation for the apparent 20-fold cellular selectivity of TAL606 for CK2α2 over CK2α1 despite an observed 2-fold biochemical sensitivity could be that CK2α1 is more sensitive to intracellular ATP concentrations, thus impeding CK2α1 target engagement in cells^69^.

Given its superior potency and selectivity for CK2α2, we decided to characterize TAL606 further. Profiling of TAL606 at 10 µM in a primary binding panel of 45 CNS receptors (see Methods for target list) and at 3 and 10 µM in a functional panel of 320 non-olfactory G-protein coupled receptors (GPCRs) resulted in EC values > 10 µM for 363/365 targets, with 2/365 targets showing showing EC values > 3 µM (Extended Data Fig. 12C). These data demonstrate that TAL606 is highly selective within the CNS-relevant proteome.

Next, we assessed pharmacokinetics (PK), brain exposure, and tolerability in mice (Fig. 6C and D, Extended Data Fig. 12D). TAL606 showed low oral bioavailability but good plasma exposure by i.p. route (Fig. 6C). PK parameters were within acceptable ranges for once-daily dosing (t_1/2_ = 8h i.p.), with brain concentrations reaching 10% of plasma. TAL606 showed target engagement in liver and mouse brains 2 hours after a single dose of 62.5 mg/kg i.p., as assessed by AKT S129 phosphorylation (Extended Data Fig.12D). We found that subcutaneous (s.c.) injection of the vehicle had superior tolerability compared to i.p. injection in vulnerable mouse populations such as aged mice, so subsequent studies were performed via the s.c. route (data not shown). TAL606 was well tolerated after 5 days of daily dosing at 10, 20, and 40 mg/kg s.c. in mice, with high levels of brain exposure (Fig. 6D; see dotted line in Fig. 6C).

Having confirmed acceptable *in vivo* pharmacology, we proceeded with a proof-of-concept efficacy study of TAL606 in an AD mouse model. The APP/PS1 transgenic AD mouse model exhibits elevated CK2 activity at 6 months of age^65^, with concomitant amyloid plaque formation and gliosis onset. We treated 2 independent cohorts of 7- to 10-month-old WT mice with vehicle (n = 26) and APP/PS1 mice with 30 mg/kg TAL606 (n = 19) or vehicle (n = 18) s.c. for 26 days, with a 5 days on/2 days off dosing paradigm (Fig. 6E). The treatment regimen was well tolerated, with no significant weight loss or clinical signs of morbidity found to be enriched in the TAL606-treated group; several animals exhibited skin reactions from s.c. injection in both TAL606 and vehicle-treated groups (Extended Data Fig. 13A). At the conclusion of the studies, serum was collected and cortex and hippocampus were dissected and fixed for immunofluorescence staining of plaques, astrocytes, and microglia. Serum clinical chemistry showed no significant adverse effects on markers of renal or hepatic toxicity (Extended Data Fig. 13B).

We observed variable plaque pathology in heterozygous APP/PS1 mice and limited our analyses to transgenic animals that showed widespread cortical and hippocampal plaque deposition, blindly excluding animals with few or no plaques prior to any further analyses (Extended Data Fig. 13C). In the hippocampus, we found that TAL606 significantly reduced both astrogliosis and microgliosis as well as co-localization of astrocytes and microglia with plaques and with each other in APP/PS1 mice (one-way ANOVAs with Sidak’s multiple comparisons tests, Fig. 6F-G). Concomitantly, hippocampal CK2 target engagement was confirmed via a reduction of NF-κB phospho-S529+ astrocytes (P = 0.0001) and microglia (P = 0.0002, one-way ANOVAs with Sidak’s multiple comparisons test, Fig. 6H, Extended Data Fig. 14A). In the cortex, we found that TAL606 significantly reduced astrogliosis and colocalization of astrocytes with plaques and with microglia, with a trend toward reduction of microgliosis in APP/PS1 mice (Extended Data Fig. 14B-C). Reactive astrocytes exhibit increased ramification in AD brains^70^, which was phenocopied in our APP/PS1 mice, as shown by an increase in total skeleton length (P <0.0001) and branching metrics relative to WT mice (Fig. 6I, Extended Data Fig. 14D). In contrast, cortical astrocytes from TAL606-treated mice showed a significant reversion toward the resting state observed in WT mice (P = 0.0018, one-way ANOVA with Sidak’s multiple comparisons test).

Plaque pathology and gliosis are already established by 7 months, when the youngest animals in our studies underwent treatment. TAL606 did not appear to affect amyloid plaque counts in the cortex or hippocampus (Extended Data Fig. 14E-F). Taken together, these data demonstrate that AD-associated gliosis can be ameliorated by CK2 inhibition *in vivo*.

## DISCUSSION

Although several anti-inflammatory drugs are being currently tested in clinical trials for neurodegenerative disease, specific pathways and central regulators of inflammation in the brain remain to be fully elucidated. Key immune players such as RIPK1 and TNFα are not brain-enriched and strategies involving modulation of these targets could have systemic off-target effects^71–73^. CK2 has been previously shown to regulate inflammatory pathways, with CK2 inhibitors demonstrating efficacy in immune-driven cancer models^40,74,75^. Through phenotype-guided studies of natural product activities, genetic approaches, and *in vitro* human disease modeling, we validated that kinase CK2, especially the CK2α2 isoform, is a potent upstream regulator of NF-κB-driven innate immunity transcriptional programs in astrocytes. We showed that CK2α2 is upregulated in AD cellular models and postmortem brains, and its overexpression in glia-enriched cortical organoids mimics AD phenotypes such as NF-κB activation, loss of synaptic proteins, tau hyperphosphorylation, and elevated amyloid β 42/40. These data are in agreement with other reports of CK2 dysregulation in AD, including CK2 promotion of tau and amyloid-β pathology and AD astrocytes exhibiting elevated CK2 levels associated with amyloid deposits ^45,59,76^.

Of the two catalytic subunit isoforms of CK2, only CK2α2 appeared to be significantly differentially expressed in multiple AD postmortem brain tissue datasets as well as AD patient-derived astrocytes. However, we cannot rule out that concomitant CK2α1 downregulation could be a contributing factor, given that reduced protein quality could offer only a partial explanation for why we observed lower CK2α1 levels in the AD Neurobiobank cohort. Depletion of CK2α1 levels could lead to effectively increased CK2α2 to CK2α1 ratios, which might skew CK2 activity in favor of CK2α2 substrates. Our analysis of CK2 isoform expression in the UPP Proteomics Study of AD postmortem brain tissue supports this hypothesis, as CK2α2 was found to be elevated relative to CK2α1 and CK2β in AD patients, but not controls. In terms of cell-type specificity, AD frontal cortex tended to show more CK2α2+ astrocytes and microglia. Although we did not observe differences in the percentage of CK2α2+ neurons in the ADRC cohort, there is a recent report that dopaminergic neurons in the VTA show CK2-dependent hyperexcitability due to SK channel dysfunction in 3xTg-AD mice^77^. Further studies are needed to characterize the role of CK2α2 in AD neurons, not only in relation to tau and amyloid pathology and SK channel dysfunction but also mitochondrial homeostasis, a pathway in which CK2 has been previously implicated^78,79^.

We validated TAL606 as a potent and highly selective CK2 inhibitor suitable for use as a chemical probe to interrogate efficacy *in vivo*. Importantly, we demonstrated that CK2 inhibition can attenuate inflammation *in vivo*, both in an acute inflammation mouse model and in a disease-relevant AD model at a time point where chronic inflammation is already established. Thus, CK2 inhibition may present a novel therapeutic avenue for patients with neurodegenerative conditions wherein neuroinflammation is a hallmark of the disease. Multiple kinase inhibitor programs in the CNS have progressed to Phase II/III trials, demonstrating the tractability and safety of brain-penetrant kinase inhibitors as a drug class for neurological diseases^80–82^. Many programs have failed due to lack of efficacy rather than toxicology, showing that kinase inhibitors with limited side effect profiles can be designed and implemented for chronic CNS indications.

TAL606 did not affect amyloid plaque pathology, an increasingly controversial marker of disease burden in AD given the high prevalence of plaques in asymptomatic, cognitively normal individuals^83^. Nevertheless, CK2 has been reported to regulate BACE1 expression upstream of amyloid processing^76^, so future studies could investigate whether earlier exposure of APP/PS1 mice to CK2 inhibition, before 6 months, could affect plaque formation.

Recently, components of the CK2 holoenzyme have been characterized as genetic risk factors in AD as well as other neurodegenerative and psychiatric diseases, including amyotrophic lateral sclerosis, multiple sclerosis, and schizophrenia^84,85^. Prior studies show that CK2 activity is abnormally elevated in the brains of Huntington’s disease (HD) patients as well as HD preclinical models^86^. CK2α2-haploinsufficient zQ175 mice exhibit reductions in disease burden, such as improvement of motor function^87^. In Parkinson’s disease, CK2 regulatory subunits were reported to co-localize with Lewy bodies, and elevated CK2 activity was shown to increase α-synuclein S129 phosphorylation^88,89^. Studies of potent and selective brain-penetrant inhibitors like TAL606 in HD and PD models are needed to further elucidate the role of CK2 in these diseases.

CK2 has been implicated in a plethora of biological processes, but recent studies have challenged the pleiotropy of CK2^42,90^. Although CK2α1 is important for development, as reflected by the embryonic lethality of germline knockout mice^64,91,92^, it is unclear if depletion of CK2α1 could be detrimental in adult animals. Our data show that chronic treatment with dual CK2α1/CK2α2 inhibitor TAL606 was well tolerated in APP/PS1 mice, although TAL606 appears to possess higher CK2α2 cellular potency. Human studies with CX-4954 appear to suggest that systemic dual CK2α1/CK2α2 inhibition is well tolerated^93^, with the caveat that CX-4945 is a promiscuous kinase inhibitor. Mouse data show that modulating CK2α2 activity is unlikely to have deleterious effects, as germline knockout mice are viable and healthy besides exhibiting reversible globozoospermia^64,91,92^. Further studies are needed to develop next-generation CK2 inhibitors with increased CK2α2 isoform selectivity, brain to plasma partitioning, and oral bioavailability, which will expand the therapeutic window for CK2 inhibitors for patients with neurodegenerative diseases.

## Acknowledgments

This work was supported in part by the National Institutes of Health/National Institute of Aging (NIH/NIA), award numbers 5 RF1 AG056306, 4 R37 AG072502, 5 R01 AG067556, 5 P01 AG051449, 5 R01 AG074307. It is subject to the NIH Public Access Policy. Through acceptance of this federal funding, NIH has been given a right to make this manuscript publicly available in PubMed Central upon the Official Date of Publication, as defined by NIH. The content is solely the responsibility of the authors and does not necessarily represent the official views of the NIH. Research was supported in part by the Freedom Together Foundation, The Dolby Foundation, and the American Heart Association and the Paul G. Allen Frontiers Group Grant #19PABHI34610000/TEAM LEADER: Fred H. Gage/2019, Milky Way Foundation, Annette C. Merle-Smith, and Lynn and Edward Streim. This work was supported by the Mass Spectrometry Core of the Salk Institute with funding from NIH-NCI CCSG: P30 014195 and the Helmsley Center for Genomic Medicine, by the Flow Cytometry Core Facility of the Salk Institute with funding from NIH-NCI CCSG: P30 014195 and Shared Instrumentation Grant S10-OD023689 (Aria Fusion cell sorter), by the NGS Core Facility of the Salk Institute with funding from NIH-NCI CCSG: P30 014195, the Chapman Foundation and the Helmsley Charitable Trust, by the GT3 Core Facility of the Salk Institute with funding from NIH-NCI CCSG: P30 014195, an NINDS R24 Core Grant and funding from NEI, and by the Stem Cell Core Facility of the Salk Institute with funding from the Helmsley Charitable Trust. The results published here are in part based on data obtained from the AD Knowledge Portal (https://adknowledgeportal.synapse.org); study data were provided through the Accelerating Medicine Partnership for AD (U01AG046161 and U01AG061357), based on specimens provided by the Emory Goizueta Alzheimer’s Research Disease Center (P50AG025688), where data collection was supported through funding by NIA grants R01AG053960, R01AG057911, R01AG061800, RF1AG057471, RF1AG057470, R01AG061800, R01AG057911, R01AG057339. The UCI-ADRC is funded by NIH/NIA Grant P30 AG066519. We thank Jim Moresco, Jolene Diedrich and Antonio Pinto for technical support. GPCRome and primary CNS receptor binding data was generously provided by the National Institute of Mental Health’s Psychoactive Drug Screening Program, Contract # 75N95023C00021 (NIMH PDSP). The NIMH PDSP is Directed by Bryan L. Roth MD, PhD at the University of North Carolina at Chapel Hill and Project Officer Jamie Driscoll at NIMH, Bethesda MD, USA. We thank the UCSD Tissue Technology Shared Resource (Valerie Estrada, Michael Rose) for immunofluorescence staining of ADRC brain sections. The Tissue Technology Shared Resource is supported by a National Cancer Institute Cancer Center Support Grant (CCSG Grant P30CA23100). We thank Jeremiah Moper at the UCSD Drug Development Pipeline for directing the TAL606 IP PK study. We thank Liping Yan at Chempartner for directing the TAL 606 IV and PO PK studies. This work was supported by the Waitt Advanced Biophotonics Core Facility of the Salk Institute (RRID: SCR_014838) with funding from NIH-NCI CCSG P30 CA014195, NIH-NIA San Diego Nathan Shock Center P30 AG068635, The Henry L. Guenther Foundation and the Waitt Foundation (Core Personnel: Daniela Boassa, Sammy Weiser-Novak, Elsie Quansah). Human tissue was obtained from the NIH NeuroBioBank (Special acknowledgement to the University of Maryland, Baltimore, MD and the NIH Brain and Tissue Repository-California, Human Brain & Spinal Fluid Resource Center, VA West Los Angeles Medical Center, Los Angeles, CA, which is supported in part by the NIH and the US Dept of Veterans Affairs) and University of California-San Diego Shiley-Marcos Alzheimer’s Disease Research Center (UCSD Shirley-Marcos ADRC, supported in part by NIH Grant: P30 AG062429). We thank Diogo H. Da Silva and Mary Lynn Gage for manuscript editing support.

## Author Contributions

I.N.D. conceived of the project, generated the data, helped analyze the results, and directed the project in coordination with F.H.G. TPP was performed and analyzed by I.N.D. Knockdown, overexpression, and CRISPR/Cas9 studies were performed by I.N.D., D.R., and J.M.E. NanoBRET and kinase capture studies were performed by I.N.D. qPCR studies were performed by D.R. Immunofluorescence studies were performed by D.R. and L.A.W. In vivo studies were performed by S.L.P., I.S.G., A.H.M, and A.L.T. Immunoprecipitation and Western blotting were performed by I.N.D., D.R., J.M.E. and L.A.W. Cloning was performed by J.M.E., R.K., and L.R.M. Inflammation and viability flow cytometry studies were performed by J.J.B. and B.N.J. NF-κB reporter and Lumit studies were performed by D.R., J.M.E, and I.N.D. Chemical synthesis and characterization were performed by J.K.T. RNA-seq of primary astrocytes was performed by M.C.M. and K.C.V. with additional analysis by I.N.D. RNA-seq of hiPSC-derived astrocytes was performed by J.B. with input from N.J.A. and analysis by I.N.D. Cell lines were generated and characterized by I.N.D., D.R., M.C.M., V.R., and M.P. Organoid studies were performed by C.L., S.F., J.P, L.A.W, and I.N.D. The manuscript was written by I.N.D. and F.H.G. and read and edited by all authors.

## SUPPORTING MATERIALS

## METHODS

### Data and materials availability

Primary RNA-sequencing data are available through GEO; source data for gene expression analyses are shown in Supplementary Tables 2, 3, and 5. Raw proteomics data are available through MassIVE and processed and analyzed data are provided in Supplementary Table 1. All source data for immunoblotting are shown in Extended Data Figures 9-12. The publicly available datasets used in this study are https://aging.brain-map.org/download/index (Aging, Dementia, TBI study) and <u>syn17009177</u> (UPP Proteomics Study). All other data generated and/or analyzed during this study are available from the corresponding authors on reasonable request. All unique and stable reagents generated in this study, of which sufficient quantities exist, are available from the corresponding authors with a complete Materials Transfer Agreement.

### Subjects

Induced human pluripotent stem cell (hiPSC) lines derived from dermal fibroblasts or peripheral blood mononuclear cells (PBMC) from donors between 63 and 87 years of age were obtained from the University of California San Diego Alzheimer’s Disease Research Center (UCSD-ADRC) and the University of California Irvine Alzheimer’s Disease Research Center (UCI-ADRC) and the Institute for Memory Impairments and Neurological Disorders (MIND) iPS Cell Bank. Protocols were previously approved by the Salk Institute Institutional Review Board and other local human subjects committees and informed consent was obtained from all subjects. Both AD and control subjects underwent clinical assessment, including detailed neuropsychological testing, and the subjects selected for biopsies were stratified to show either a clear non-demented clinical picture or clear phenotype of AD (classified as probable AD until pathological confirmation). They were followed up after biopsy with no evidence of cognitive decline (controls) and with evidence of progressive impairment (AD). Control hiPSC line R88 was a kind gift from Jeffrey Jones. Supplementary Tables 4 and 8 contain available subject information.

### Cell culture

Primary fetal human cerebellar astrocytes (HCA, ScienCell) were cultured in Astrocyte Medium (AM, ScienCell). Human embryonic stem cell (hESC) and hiPSC lines were maintained on Matrigel (Cultrex)-coated plates in TeSR medium (made in-house by the Salk Stem Cell Core), fed daily and passaged with dispase every 5-7 days (Gibco). All subjects provided written informed consent and all procedures were approved by local human subjects committees. Mature hESC- or hiPSC-derived astrocytes were cultured in DMEM/F12 Glutamax (Thermo Fisher Scientific) supplemented with N2 and B27 (Thermo Fisher Scientific) and 10% fetal bovine serum (FBS, Biowest) or serum-free medium (see below). Mature hiPSC-derived microglia were cultured in DMEM/F12 Glutamax serum-free medium (see below) supplemented with TGF-b1, IL-34, and M-CSF. THP-1 NF-κB-Lucia reporter cells (InvivoGen) were grown in RPMI 1640 with 10% FBS, Glutamax, 25 mM HEPES pH = 7.5, and penicillin/streptomycin (P/S), and were differentiated into monocytes with 10 ng/mL PMA for 3 days. HEK293T were grown in DMEM with 10% FBS, Glutamax, and P/S. All cell lines were maintained in a humidified incubator (5% CO_2_) at 37 °C and were routinely tested for mycoplasma.

### Generation of glia-enriched cortical organoids

hiPSCs were cultured in accordance with WiCell protocols^1,2^. <u>hiPSC</u> colonies were cultured on Cultrex RGF BME (R&D Systems; 3433-005-01)-coated, 6-well tissue culture treated plates using mTeSR Plus (STEMCELL Technologies, 85850) and were maintained in 5% CO_2_ humidified incubators at 37 °C. hiPSC colonies were passaged at a ratio of 1:6 every 4-6 d using Gentle Cell Dissociation Reagent (STEMCELL Technologies, 100-0485) for 5 min at 37 °C and mechanical scraping with a cell lifter. Prior to GEO generation, hiPSCs were adapted to irradiated mouse embryonic fibroblast feeders (iMEFs; R&D Systems, PSC001) by passaging 1 well of a 6-well plate of hiPSCswhen approximately 80% confluent at a ratio of 1:3 to a 6-well tissue culture-treated plate seeded at 1.5 × 10^5^ iMEFs per well. On 0 d, <u>hiPSCs</u> were seeded in 50% mTeSR Plus and 50% hESC medium containing DMEM/F12 (11330-032), 20% knockout serum replacement (10828-028), 1X MEM non-essential amino acids (MEM NEAA; 11140-050), 1X GlutaMAX (35050-061), 0.1% (55 μM) 2-Mercaptoethanol (21985-023), and 10 ng/ml bFGF (100-18B) all from Thermo Fisher Scientific. From 1 d onwards, hiPSCs were cultured in hESC medium. hiPSCs were passaged at a ratio of 1:2 to a 6-well tissue culture-treated plate seeded at 1.5 × 10^5^ iMEFs per well when colony sizes reached approximately 1.5 mm in diameter using 1 mg/ml collagenase type IV (Thermo Fisher Scientific, 17104019) for 10-40 min (until colony edges began to curl) at 37 °C and mechanical scraping with a 5-ml glass pipette. All <u>hiPSCs</u> were maintained below passage 50 and confirmed negative for mycoplasma. Cells were confirmed to be karyotypically normal. All experiments involving cells from human subjects were performed in compliance with the institutional Embryonic Stem Cell Research Oversight (ESCRO) committee.

GEOs were generated as previously described^3,4^ with some modifications allowing for long-term, serum-free culture. Embryoid bodies (EBs) were generated when the colonies of one entire 6-well tissue-culture plate reached approximately 1.5 mm in diameter. To generate EBs, intact hiPSC colonies (cell lines described in Supplementary Table 8^5^) were treated with 1 mg/ml Collagenase type IV (Thermo Fisher Scientific, 17104019) for 20–45 min (until colony edges began to curl) at 37 °C and were mechanically detached using a cell lifter. Lifted colonies were maintained in a 10 cm ultra-low-attachment dish (Nunc) with hESC medium containing 4 ng/ml bFGF and 10 µM ROCK inhibitor Y-27632 (Tocris Bioscience, 1254). On 1 d, EBs were switched to astrocyte medium (AM; ScienCell, 1800) supplemented with astrocyte growth supplement (AGS; ScienCell, 1800), 2% FBS, 500 ng/ml Noggin (R&D Systems, 6057-NG), and 10 ng/ml PDGF-AA (Proteintech, HZ-1215) until 14 d. EBs were cultured under stationary conditions from 0-4 d and were transferred to an orbital shaker set to 75 rpm from 4 d onwards. On day 15, noggin was excluded from the medium and organoids were cultured in AM supplemented with AGS, 2% FBS, and 10 ng/ml PDGF-AA. On day 21, organoids were maintained in medium consisting of DMEM/F-12 (Thermo Fisher Scientific, 11330-032), 1X N2 supplement (Thermo Fisher Scientific, 17502–048), 1X B27 (without RA; Thermo Fisher Scientific, 12587-010), 1X MEM NEAA, 1X GlutaMAX, 1X insulin-transferrin-selenium (ITS; Thermo Fisher Scientific, 41400-045), 5 µg/ml insulin solution human (Sigma-Aldrich, I9278), 1X antibiotic antimycotic solution (Sigma-Aldrich, A5955), and 0.1% (55 µM) 2-Mercaptoethanol. Organoids cultured beyond day 75 were cultured in the previously described medium formulation with 50% DMEM/F12 and 50% Neurobasal-A (Thermo Fisher Scientific, 10888-022) and were additionally supplemented with 20 ng/ml brain-derived neurotrophic factor (BDNF; R&D Systems, 248-BDB), 20 ng/ml glial cell line-derived neurotrophic factor (GDNF; R&D Systems, 212-GD), 0.2 µM ascorbic acid (AA; STEMCELL Technologies, 72132) and 245 µg/ml cyclic adenosine monophosphate (cAMP; Tocris Bioscience, 1141). GEO media changes were performed every other day.

### Lentiviral transduction of organoids

Organoids were transduced with approximately 2E+06 lentiviral particles encoding either GFP (pBOB-CAG-P2A-eGFP-PGKpuro) or CK2A2 -P2A-GFP (pBOB-CAG-CK2A2-P2A-eGFP-PGKpuro) under the CAG promoter. Viral particles were added directly to the culture medium and incubated with the organoids for 48 hr at 37°C in a humidified incubator with 5% CO₂. At 2 days post-infection, the medium was replaced with fresh organoid culture medium. Conditioned media were subsequently collected every 7 days and cleared from cell debris by centrifugation at 3,000 g for 10 min at 4°C for downstream analyses. One month following transduction, organoids were either washed twice with PBS and snap frozen in LN2 for downstream analyses or prepared for sectioning as described below.

### Animals and drug treatment

All animal procedures were approved by the Institutional Animal Care and Use Committee of The Salk Institute for Biological Studies. Experiments were conducted in accordance with the National Institutes of Health’s Guide for the Care and Use of Laboratory Animals and the US Public Health Service’s Policy on Humane Care and Use of Laboratory Animals. Mice were group housed under a standard 12-hr light/dark cycle with *ad libitum* access to rodent chow and water. For CHR treatment experiments, male C57Bl6/NHsd mice 8-16 weeks of age were used. For assessing *in vivo* pharmacokinetics, mice received a single i.p. injection of chrysoeriol (CHR, 40 mg/kg) or vehicle (3% wt/vol HPMC). Mice were euthanized for blood and tissue collection 1, 4, or 24 hr later. For assessing inflammation *in vivo*, mice received daily treatments of CHR at 37 mg/kg or vehicle (5% DMSO and 95% corn oil) injected i.p. for 4 consecutive days. On day 4, mice received a single i.p. injection of TNFα (250 ug/kg) or saline (SAL) 1 hr after CHR or vehicle pretreatment. Mice were then euthanized for blood and tissue collection 3 hr later. At terminal time points, mice were deeply anesthetized with a ketamine/xylazine/acepromazine cocktail injected i.p. Cardiac puncture was performed, tissues were dissected and flash frozen on dry ice and then transferred to -80°C storage until analysis.

Hemizygous APP/PS1 mice (Jax stock #005864) and their wildtype littermates were used to assess CK2 inhibition *in vivo*. Mice were 7-10 months old at the beginning of the experiment. Mice were treated chronically with TAL606 (30 mg/kg) or vehicle solution (90% corn oil, 5% DMSO, 5% Tween 20) via subcutaneous injection. Injections were performed daily Monday through Friday for 4 weeks. Mice were euthanized 1hr following the final treatment by overdose with a ketamine, xylazine, and acepromazine anesthetic cocktail. Blood was collected via cardiac puncture, and then mice were transcardially perfused with 0.9% NaCl. Whole blood was allowed to clot for at least 30 min, then spun for 10 min at 1,000g at 4°C to separate serum and stored at -80°C. Half of the brain was dissected, flash frozen over dry ice, and stored at -80°C for RNA and protein analysis. The other half was drop-fixed in 4% PFA for immunofluorescence assays. Frozen serum samples were shipped to IDEXX BioAnalytics (North Grafton, MA) for analysis on the Mouse Standard Tox Panel (62794).

To assess the tolerability of repeated TAL606 treatments *in vivo*, adult (3-month-old) male C57BL6/NHsd mice were treated with TAL606 (10, 20, or 40 mg/kg) or vehicle solution (90% corn oil, 5% DMSO, 5% Tween 20) via subcutaneous injection daily for 5 days. Formulation was prepared as follows: TAL606, DMSO, and Tween 20 were heated with stirring to 90°C for thirty minutes. The mixture was then cooled to RT, diluted with requisite corn oil, and vortexed. Mice were monitored daily by visual inspection and body weight measurement. At 1hr following the final treatment, mice were euthanized for blood and tissue collection under terminal anesthesia. Blood serum was collected as above. Brains and liver samples were extracted and stored at -80°C.

To assess TAL606 pharmacokinetics and pharmacodynamics in brain, adult (2- to 3-month-old) male C57BL6/NHsd mice were given a single TAL606 injection of 62.5 mg/kg i.p. or vehicle and euthanized 1 or 2h later. Blood serum, brain, and liver samples were collected and stored at -80°C. One-fourth of whole brain or liver was dissected on dry ice with a clear razor and transferred to a prechilled snap-top eppendorf tube containing 0.9 to 2.0 mm stainless steel beads (Next Advance) and non-denaturing lysis buffer (20 mM HEPES 7.4 pH, 150 mM NaCl, 1% Triton-X, 1 mM EDTA, 1 mM EGTA, Halt Protease/Phosphatase inhibitors). Samples were then homogenized with Bullet Blender at 4°C for 3 min at level 8 and then sonicated for 3 cycles, each consisting of 5 sec of active sonication, 5 sec off at 25% amplitude (Epishear Sonicator). Homogenates were clarified at 20,000g for 10 min at 4°C. Lysates were processed for Western blotting as described below.

To assess IV and PO pharmacokinetics, adult (6- to 8-week-old) male CD-1 mice were given a single TAL606 dose of 5 mg/kg by tail vein injection (IV) or 30 mg/kg by oral gavage (PO), with serial bleeding sampling at 0.083, 0.5, 1, 2, 4, 8, and 24 hr (N = 3 mice per group). IV dosing formulation was 10% DMSO, 10% Solutol HS 15, and 80% Saline; PO dosing formulation was 90% corn oil, 5% DMSO, and 5% Tween-20 prepared as above. At the designated time points, approximately 30 µL of blood samples were taken from the via facial vein into K2EDTA tubes. The blood sample was put on ice and centrifuged at 2,000 g for 5 min to obtain plasma sample within 15 min then transferred into Eppendorf tubes. Plasma samples were stored at -70℃ until LC-MS/MS analysis. No abnormal clinical symptom was observed during the in-life phase.

To assess IP pharmacokinetics, adult male C57Bl/6J mice (n=6) weighing 20–25 gm were used. Mice were group-housed, five per cage, in a temperature-controlled (22°C) vivarium on a 12 hr/12 hr light/dark cycle with ad libitum access to food and water. Each mouse received a single 25.6 mg/kg TAL606 dose administered (I.P.) in a 3 wt% HPMC DI water solution. Whole blood samples were collected via retro-orbital bleeds at 0.5, 1, 2, 4, 8 and 24 hr post-dose and processed to plasma. Brain and liver samples were collected at the terminal 24 hr time point. Plasma and tissue concentrations of TAL606 were quantitated using liquid chromatography with tandem mass spectrometry (LC-MS/MS). TAL606 standards were prepared from a methanol stock in blank mouse plasma and tissue homogenates and used to generate external calibration curves using linear regression to plot the peak area ratio versus concentration with 1/x weighting (r2 ≥ 0.99) over the analytically reportable range. All animal procedures were conducted in strict adherence to the National Institutes of Health Guide for the Care and Use of Laboratory Animals and approved by the University of California San Diego Institutional Animal Care and Use Committee (IACUC).

### Mass spectrometry for TAL606 PK studies

Plasma samples were analyzed on a Triple Quad 6500+ instrument fitted with a Waters X-Bridge Premier BEH C8 column (2.1×50 mm, 2.5 µm). To generate 9-point calibration standard curves of 1.00-3,000 ng/mL TAL606 in mouse plasma, 5 µL of TAL606 standards in methanol were mixed with 95 µL blank plasma. Non-diluted test plasma samples (5 µL) were transferred to a 96-well plate along with calibration standards and blanks. To single blanks, calibration standards, QCs and test samples, 150 µL of IS (Glipizide, 50 ng/mL) in acetonitrile was added; to double blanks, 150 µL acetonitrile was added. The mixture was shaken for 10 min and centrifuged at 5,800 rpm for 10 min at 4°C. Next 70 µL of supernatant was transferred to a new plate, and 3 µL of solution was injected in the LC-MS/MS instrument. The following HPLC solvents were used: Mobile Phase A: Water-0.025% Formic acid-1mM Ammonium acetate; Mobile Phase B: Acetonitrile:Water(95:5,V/V)-0.025% Formic acid-1mM Ammonium acetate. At a flow rate of 0.50 mL/min and column temperature of 50°C, the following gradient was used: equilibration at 15 % B in 0.2 min, 15-40% B in 0.2 min, 40-90 % B in 1.1 min, 90 % B in 0.9 min, 90-15 % B in 0.01 min, and re-equilibration at 15% for 0.6 min for a total run time of 3 min. MS analyses were performed using electrospray ionization in positive mode. Multiple reaction monitoring (MRM) was performed using Q1/Q3 masses 428.00/396.00 Da, with glipizide as internal standard (Q1/Q3 Masses: 446.20/321.10 Da). Peak area at retention time of 1.53 min was measured and normalized to the internal standard (1.29 min). Compound amount was calculated using the standard curve. PK parameters were estimated by non-compartmental model using WinNonlin 8.2 (Pharsight Corporation). The bioavailability (F%) was calculated as following: AUC_last_-PO/AUC_INF_-PO > 80%: F=(AUC_INF_-PO*DoseIV)/(mean AUC_INF_-IV*DosePO) AUC_last_-PO/AUC_INF_-PO ≤ 80% or AUC_INF_ was not available: F=(AUC_last_-PO*DoseIV)/(mean AUC_last_-IV*DosePO)

### APP/PS1 mouse brain immunofluorescence staining

Saline-perfused APP/PS1 mouse brains were sagittally sectioned in half, drop-fixed in 4% PFA at 4 °C, and transferred to 30% sucrose solution prior to sectioning. Brains were sectioned coronally at 40 µm thickness using a microtome. Sections were stored in tissue cryo-protection solution (25% glycerol, 30% ethylene glycol in PBS) at -20°C until used for immunostaining. Free-floating sections (4-8. 480 µm apart) were washed 3x in 1xTBS for 5 min then blocked in TBS++(3% Normal serum in TBS with 0.25% Triton-X) for 60 min. Sections were incubated in primary antibody overnight for 60-72 hr at 4°C on a shaker, followed by washes in TBS for 15 min and a 30-min wash in TBS++. Secondary antibodies Cy3, AF488, and AF 647 were applied (1:250) protected from light for 2 hr at RT, followed by a TBS++ wash for 15 min and a final wash of TBS for 15 min. A counter stain of DAPI at 1 µg/mL in TBS for 15 min was performed, followed by a final wash in TBS at 15 min. Sections were mounted on slides using Immu-Mount (Epredia: Cat# 9990402) with no. 1.5 coverslips and dried overnight. For initial plaque pathology assessment, imaging was performed on a Zeiss Axioscan 7 slide scanner using a 20x 0.5NA objective. Area occupied by plaques was determined using QuPath (version 0.5.1). An ROI was drawn manually around the cortical tissue from a single hemisphere and 3-4 images were acquired per mouse. For detailed plaque burden and gliosis assessment, imaging was performed using 10-20x magnification objectives on a Zeiss Airyscan 880 confocal microscope, employing Z stacks. Three images were acquired per mouse unless otherwise specified. Primary Antibodies used: Rabbit monoclonal anti-β-Amyloid (D54D2) XP (1:500; CTS: 8243), Chicken polyclonal anti-GFAP (1:5000; EDM Millipore: AB5541), Goat polyclonal anti-Iba1 (1:200; Abcam: ab5076), Mouse monoclonal anti-NeuN (1:500; EMD Millipore: MAB377), Guinea pig polyclonal anti-Doublecortin (1:1000; EDM Millipore: AB2253), Chicken polyclonal anti-MAP2 (1:1000; Abcam: ab5392), Rabbit polyclonal anti-Prox1 (1:500; Abcam: ab101851), Rat monoclonal anti-BrdU (1:250; Abcam: ab6326), Rabbit polyclonal anti-Phospho-NF-κB(Ser52) (1:50; Invitrogen: 44-711G), Chicken polyclonal anti-GFAP (1:5000; EDM Millipore: AB554).

### Postmortem samples

Fresh frozen tissue samples were acquired from the NIH Neurobiobank, from deceased individuals with AD or unaffected controls from Brodmann Area 39 of the parieto-temporal cortex (n=10 per group) (see Supplementary Table 6 for clinical characteristics). Approximately 5 mg of tissue was cut on dry ice with a clear razor and transferred to a prechilled snap-top Eppendorf tube containing 0.5 mm zirconium oxide beads and 300 µL non-denaturing lysis buffer (20 mM HEPES 7.4 pH, 150 mM NaCl, 1% Triton-X, 1 mM EDTA, 1 mM EGTA, Halt Protease/Phosphatase inhibitors). Samples were then homogenized with Bullet Blender at 4°C for 3 min at level 9. Homogenates were clarified at 20,000g for 10 min at 4°C and used for Western blotting. Formalin-fixed paraffin-embedded (FFPE) tissue blocks containing frontal cortex were acquired from the UCSD ADRC, from deceased individuals with early to mid-stage AD or from age- and sex-matched cognitively normal controls (n=10 per group) (see Supplementary Table 7 for clinical characteristics).

### Postmortem samples sectioning and immunofluorescence staining

FFPE tissue blocks were sectioned at 5 µm on a Leica microtome. In preparation for staining, tissues were baked at 60°C for 1 hr. Tissues were cleared and rehydrated through successive alcohol washes (3X Xylene, 2X 100% EtOH, 2x 95% EtOH, 2X 70% EtOH, diH_2_O). Antigen retrieval was performed in Antigen Unmasking Solution (Citrate Based, pH6) (Vector, H-3300) at 95°C for 30 min. Staining was performed on Intellipath Automated IHC Stainer (Biocare). First, peroxidase block Bloxall (Vector, SP-6000) was applied for 10 min, followed by 2X wash in TBST. Then, protein block Blotto (Thermo, PI37530) was applied for 10 min. Slides were incubated with primary antibodies for 1h, followed by 2X TBST washes, 30-min incubation with secondary antibodies, 2X TBST washes, and 2X diH_2_O washes. Slides were counterstained with DAPI (1µg/ml) for 15 min. Slides were mounted and coverslipped with Vectashield Vibrance (Vector, H-1700-10). Slides were scanned using an Akoya Fusion PhenoImager instrument at 10X magnification. The following primary antibodies were used: Anti-CK2A1 (Rabbit, Bethyl, A300-198A, 1:5000), Anti-CK2A2 (Rabbit, ProteinTech, 10606-1-AP, 1:100), Anti-CK2B (Rabbit, Abcam, ab76025, 1:100), GFAP (Chicken, EMD, AB5541, 1:1000), IBA1 (Goat, ab5076, Abcam, 1:100), and MAP2 (Mouse, Sigma, M1406, 1:500). The following secondary antibodies were used: Anti-Rabbit IgG-AF647 (Invitrogen, A-21245, 1:200), Chicken IgG-AF488 (Invitrogen, A78948, 1:200), Goat IgG-AF594 (Invitrogen, A32758, 1:200), Mouse IgG-AF546 (Invitrogen, A10036, 1:200).

### Postmortem tissue imaging analysis

Regions of interest (ROIs) were manually drawn around gray matter (excluding any areas with artifacts) and ROIs were exported as 256×256 µm image tiles in TIF format using Qupath (v0.4.2); the total number of tiles exported per patient ranged from approximately 200 to 2,300. Image tiles were analyzed in CellProfiler (4.2.5). Astrocytes (GFAP+/DAPI+), microglia (IBA1+/DAPI+), neurons (MAP2+/DAPI+), nuclei (DAPI+), and CK2A1-, CK2A2-, or CK2B-positive objects were identified using IdentifyPrimaryObjects and RelateObjects modules with Global Minimum Cross-Entropy (astrocytes, microglia, CK2A1/CK2A2/CK2B), Global Robust Background (nuclei), or Adaptive Minimum Cross-Entropy (neurons) thresholding using appropriate parameters. CK2A1-, CK2A2-, or CK2B-positive astrocytes, microglia, and neurons were identified using RelateObjects modules by filtering only cells overlapping with CK2A1-, CK2A2-, or CK2B-positive objects. Percent positive cells were identified using the CalculateMath module by dividing counts of CK2A1-, CK2A2-, or CK2B-positive astrocytes, microglia, and neurons by total counts of astrocytes, microglia, and neurons and multiplying by 100. For each individual, mean values of percent positive cells as well as cumulative distributions of percent positive cells across ROIs were generated, analyzed and plotted in GraphPad Prism.

### APP/PS1 mice imaging analysis

Single channel maximum projection images (MPIs) were generated from raw images using CellProfiler. ROIs were manually drawn for each image set using the DAPI channel and binary masks delineating each ROI were exported using Fiji. MPIs were analyzed using CellProfiler pipelines by first masking each image using the ROI binary images in MaskImage modules. Then, images were smoothed with sigma 1 using GaussianFilter modules and binary images of plaques (amyloid B), astrocytes (GFAP), microglia (Iba1), and nuclei (DAPI) were generated using Threshold modules with Global Minimum Cross-Entropy as the thresholding strategy, with appropriate parameters. ConvertImageToObjects was used to convert binary images into objects. The area occupied in each channel was determined using MeasureImageAreaOccupied of respective binary images, whereas total intensity was calculated using MeasureImageIntensity in each channel only within objects. The area occupied and total intensity values were normalized to ROI volume (area of mask multiplied by the number of z-stacks used to generate the MPI). Plaque counts were determined using the IdentifyPrimaryObjects module in the amyloid-beta channel and Global Minimum Cross-Entropy as the thresholding strategy, with appropriate parameters. Manders coefficients were calculated using the MeasureColocalization module across the amyloid-beta, GFAP, and IBA1 channels. Astrocyte morphology in the cortex was determined by the MorphologicalSkeleton and MeasureObjectSkeleton modules applied to GFAP binary images (filtered for minimum and maximum sizes to remove artifacts such as clumped cells). For phospho-NFKB analysis in the hippocampus, astrocytes (GFAP), microglia (IBA1), and pNFKB+ objects were identified using IdentifyPrimaryObjects and RelateObjects modules with Global Minimum Cross-Entropy thresholding using appropriate parameters. pNFKB+ astrocytes or microglia were identified using RelateObjects modules by filtering only cells overlapping with pNFKB+ objects. Percent positive areas were identified using the CalculateMath module by dividing counts of pNFKB+ astrocytes or microglia by total counts of astrocytes or microglia and multiplying by 100. Images were visualized in Fiji by applying the same channel settings across treatment groups. Data from all tiles for each mouse were pooled together and plotted in GraphPad Prism.

### Organoid imaging analysis

ROIs were manually drawn around neuron and/or astrocyte-containing areas (excluding any areas with artifacts) and ROIs were exported as 176.3×176.3 µm image tiles in TIF format using Qupath; total number of tiles exported per organoid ranged from approximately 15 to 60. Images were then analyzed using CellProfiler pipelines by first masking each image using the ROI binary images in MaskImage modules. Then, binary images of neurons (NeuN), astrocytes (SOX9), GFP or CK2A2, and nuclei (DAPI) were generated using Threshold modules with adaptive Sauvola as the thresholding strategy, with appropriate parameters. MaskImage modules were used to generate binary images of SOX9+/NeuN-/GFP+, SOX9+/NeuN-/CK2A2+, SOX9-/NeuN+/GFP+, and SOX9-/NeuN+/CK2A2+ areas. Percent area occupied was then calculated by MeasureImageAreaOccupied modules, dividing area covered by GFP or CK2A2 positive astrocytes or neurons by total area covered by astrocytes or neurons. MaskImage modules were used to generate images of SOX9+/NeuN- or SOX9-/NeuN+ areas in GFP or CK2A2 channels. Total intensity within astrocytes or neurons was then calculated using MeasureImageIntensity.

### CK2 activity ELISA

Postmortem brain lysates were assayed for CK2 activity in duplicate according to manufacturer’s instructions (CycLex CK2 Activity Kit CY-1170). Briefly, 10 µL of samples (20 µg protein per well) were transferred to ELISA strips, 90 µl kinase reaction buffer was added, and the plate was incubated at 30°C for 30 min. Plates were washed 5X with wash buffer. Then, 100 µl of HRP conjugated Detection antibody TK-4D4 was added and the plate was incubated at 25°C for 30 mins. Plates were washed 5X with wash buffer. Next, 100 µl of substrate reagent was added and the plate was incubated at 25°C for 15 mins, and 100 µl of Stop Solution was added and absorbance at 450 and 560 nm was measured. Raw CK2 enzymatic activity values were normalized to total CK2 expression divided by GAPDH expression as quantified by Western blotting using ImageStudio software.

### NIMH PDSP GPCRome study

Compound screening in the PRESTO-Tango GPCRome panel was carried out as previously described^6^ by the NIMH Psychoactive Drug Screening Program. HTLA cells were plated in DMEM with 10% (v/v) FBS and 10 U/mL penicillin– streptomycin and transfected using a PEI in-plate method. PRESTO-Tango receptor DNAs were resuspended in OptiMEM, hybridized with PEI, diluted, and aliquoted into 384-well plates containing cells for overnight incubation. Tango assay buffer (20 mM HEPES, 1x HBSS, pH 7.4) or TAL606 in Tango assay buffer was added at a final concentration of 3 µM or 10 µM and plates were incubated overnight. The following day, medium was removed and BrightGlo reagent (Promega) diluted 20-fold with Tango assay buffer was added. The plate was incubated for 20 min at RT in the dark before luminescence measurement.

### Radioligand binding assays

Radioligand binding studies of TAL606 were completed by the NIMH Psychoactive Drug Screening program for 5-HT1A, 5-HT1B, 5-HT1D, 5-HT1E, 5-HT2A, 5-HT2B, 5-HT2C, 5-HT3, 5-HT5A, 5-HT6, 5-HT7A, Alpha1A, Alpha1B, Alpha1D, Alpha2A, Alpha2B, Alpha2C, Beta1, Beta2, Beta3, BZP Rat Brain Site, D1, D2, D3, D4, D5, DAT, DOR, GABAA, H1, H2, H3, H4, KOR, M1, M2, M3, M4, M5, MOR, NET, PBR, SERT, Sigma 1, and Sigma 2. Procedures were previously described in detail at https://pdsp.unc.edu/pdspweb/content/UNC-CH%20Protocol%20Book.pdf. In brief, membrane suspensions containing human recombinant receptors were collected from stable or transiently transfected cell lines. For each receptor, TAL606 was tested at 10 μM in quadruplicate in 96-well plates, in a final of volume of 125 μl per well in respective binding buffers, with hot ligand concentrations near Kd values. Total binding and nonspecific binding were determined in the absence and presence of 10 μM of reference compounds, respectively. Plates were incubated at RT in the dark for 90 min, and reactions were stopped by vacuum filtration onto 0.3% polyethyleneimine-soaked 96-well filter mats using a 96-well Filtermate harvester, followed by 3 washes with cold wash buffers. Scintillation cocktail was then melted onto the microwave-dried filters on a hot plate and radioactivity was counted in a Microbeta counter.

### MDR1-MDCKII permeability studies

Bidirectional permeability and efflux ratio was measured in MDR1-MDCKII cells by Eurofins Discovery (Catalog item g363). Compounds were tested at 10 µM, pH 7.4/7.4 in duplicate according to standard procedures^7^, with the addition of 2% BSA to increase compound percent recovery. A-B permeability was measured at 0 and 60 min, while B-A permeability was measured at 0 and 40 min, and both incubations were performed at 37°C. Colchicine, labetalol, propranolol, and ranitidine were used as reference compounds. Compounds were quantitated by HPLC-MS/MS.

### Organoid sectioning and immunofluorescence staining

Organoids were washed 1-2 times with PBS and fixed in cold 4% paraformaldehyde (PFA) for 30-60 min at RT and transferred to 30% sucrose solution at 4°C overnight for cryoprotection. Organoids were allowed to settle before sectioning. To prepare for freezing, organoids were placed into block molds and embedded in a tissue-freezing medium (15-183-13 General Data Healthcare). Prior to sectioning, blocks were allowed to equilibrate at the cryostat temperature (approximately 20-30 minutes). Sections were cut at 30 µm thickness using a cryostat, added to glass slides and stored at -20°C until used for staining. All washes were performed on a shaker and all incubations were carried out in a humidity chamber. Slides containing organoid sections were thawed at RT for 10 min and the borders surrounding the sections were outlined using a hydrophobic PAP pen. Samples were washed once with PBS for 5 min, followed by permeabilization in PBS containing 0.1% Triton X-100 for 10 min. After permeabilization, sections were blocked (5% normal donkey serum, 0.1% Triton X-100 in PBS) for 1 hour at RT. Primary antibodies diluted in the blocking solution were applied and incubated overnight at 4°C. Following incubation, slides were kept at RT for 5 min and then washed 3 times with PBT (PBS + Tween) for 5 min each. Secondary antibodies (Cy3, AF488, and AF647), diluted 1:200 in blocking solution, were applied and incubated for 1 hr at RT, protected from light. Post-incubation, slides were washed 3 times with PBT for 5 min each, followed by staining with DAPI (1 µg/mL) diluted 1:1000 in PBT for 5 min. Slides were then washed twice in PBS for 5 min each. Finally, sections were mounted using Immu-Mount (Epredia: Cat# 9990402) and no. 1.5 coverslips. Slides were allowed to dry for at least 48 hr at RT to optimize signal intensity before imaging at 20X magnification using a Zeiss Axioscan 7 Slide Scanner. Primary Antibodies used: Chicken anti-GFP (1:300; Aves Labs: GFP-1020), Goat polyclonal anti-hSox9 (1:250; R&D Biosystems: AF3075), Rabbit monoclonal anti-NF-κB p65/RelA(1:400; CTS: 8242), Mouse monoclonal anti-Phospho-RelA/NFkB p65 (S529) (1:50; R&D Systems: MAB7624), Rabbit polyclonal anti-CSNK2A2 (1:100; Proteintech: 10606-1-AP); Chicken polyclonal anti-NeuN (1:250; Millipore: ABN91).

### Tau and phospho-tau ELISA

Phospho-Tau (Thr205) and Total Tau levels were measured using the FastScan Tau ELISA kits (Cell Signaling Technology 51580, 57519). Organoid lysates (Buffer: 20 mM HEPES 7.4 pH, 150 mM NaCl, 1% Triton-X, 1 mM EDTA, 1 mM EGTA, Halt Protease/Phosphatase inhibitors) were diluted to 120 µg/mL (for phospho-tau) and 3 µg/mL (for total tau) in 1X cell extraction buffer (69905). Samples were incubated in duplicate with antibody cocktail for 1 hour at RT on a shaker. The plate contents were aspirated and washed 3x with wash buffer. After 15 min of TMB substrate incubation (protected from light), absorbance at 450 nm was measured within 30 min of the addition of stop solution (Promega Discovery GloMax). All experiments were performed within 4 weeks of antibody reconstitution.

### Amyloid β 42/40 ELISA

Amyloid β 40 (Aβ40, Invitrogen: KHB3481) and Aβ42 (Invitrogen: KHB3441) levels in organoid media were measured by ELISA. Organoid media samples were diluted 1:2 for Aβ42 and 1:5 for Aβ40 using respective diluent buffers with 1X Halt proteinase/phosphatase inhibitor cocktail (Thermo Scientific: 78430). Standards were reconstituted and serially diluted per manufacturer instructions. Samples were incubated in duplicate with the capture antibody detection system for 3 hr at RT, washed 4 times, and then incubated with IgG-HRP for 30 min. After additional washes, stabilized chromogen was added and incubated for 30 min. Absorbance at 450 nm was measured using the Promega Glomax Discover within 30 min of adding TMB stop solution. Aβ42/40 ratios were calculated in Excel and plotted in GraphPad Prism.

### Kinetic solubility assays

Compounds were shipped to Eurofins (St. Charles, MO) for testing. To determine aqueous solubility, the compound was incubated in PBS, pH 7.4, for 24 h using the shake-flask method. Chromatograms of the test compound in PBS (500 μM) or in organic solvent (500 μM, methanol/water, 60/40, v/v; calibration standard) along with UV/VIS spectra with labeled absorbance maxima were generated. Aqueous solubility (μM) was determined by comparing the peak area of the principal peak in the calibration standard (500 μM) with the peak area of the corresponding peak in the PBS sample. In addition, chromatographic purity (%) was defined as the peak area of the principal peak relative to the total integrated peak area in the HPLC chromatogram of the calibration standard.

### Human plasma and mouse brain tissue binding

Compounds were shipped to Cyprotex (Framingham, MA) for testing. For assessment of plasma binding, compound or reference standard (warfarin) was added at a final concentration of 5 µM in duplicate to human plasma (pH 7.4, ± 0.1). The mixture was dialyzed in a RED device (Rapid Equilibrium Dialysis, Pierce) per the manufacturers’ instructions against PBS and incubated on an orbital shaker. The system was equilibrated at 37°C for 4 hr. At the end of the incubation, aliquots from both plasma and PBS sides were collected and were matrix-matrix matched with an appropriate amount of PBS and blank plasma, respectively. Acetonitrile (3 volumes) containing an analytical internal standard (IS) was added to precipitate the proteins and release the test article. After centrifugation, the supernatant was transferred to a new plate and analyzed by LC-MS/MS to obtain peak area ratios (analyte/IS) for determining the fraction unbound.

For brain protein binding, mouse brain was homogenized in 3 mL of Dulbecco’s PBS (DPBS) per 1 gram of brain tissue. Compound or reference standard (diazepam) was added at a final concentration of 5 µM. The resulting mixture was dialyzed in a RED Device (Pierce) per the manufacturers’ instructions against PBS and incubated on an orbital shaker. The system was equilibrated at 37°C for 4 hr. After the end of the incubation, aliquots from both brain homogenate and PBS sides were collected. To the brain fraction, 50 μL of PBS was added to 20 μL of brain homogenate fraction. To the PBS side, 20 μL of brain homogenate was added to 50 μL of PBS fraction. Methanol containing internal standard was added to precipitate the proteins and release the agents. After centrifugation, the supernatant was transferred to a new plate and analyzed by LC-MS/MS to determine compound concentration.

Fraction unbound (fu) values in plasma were calculated using the equation fu= PF/PC, where:

PC = test compound concentration in protein-containing compartment

PF = test compound concentration in protein-free compartment

Fraction unbound (fu) values in diluted brain tissue were calculated as detailed below:

fumeas = PF/PC (fraction unbound using diluted brain tissue)

fubrain=1/D/[(1/fumeas)-1) +1/D]

fubrain = fraction unbound in brain

D = dilution factor of brain tissue

### Mouse cytokine multiplex assay

Snap-frozen dissected tissues were thawed into non-denaturing buffer (20 mM HEPES 7.4 pH, 150 mM NaCl, 1% Triton-X, 1 mM EDTA, 1 mM EGTA, Halt Protease/Phosphatase inhibitors) and homogenized using a Bullet Blender 24 Gold instrument (NextAdvance). Lysates were processed for Western blotting as described below and for cytokine expression using a multiplex bead-based assay according to manufacturer’s instructions (Mouse Inflammation 7-Plex Panel 1, GeniePlex MOAMPM019). Samples were analyzed using the HTS module of a FACSymphony A3 instrument (BD) and raw data were analyzed using FlowJo v10.8.0.

### Compounds

Phorbol 12-myristate 13-acetate was sourced from Fisher Scientific (NC9325685). CX-4945 was sourced from Sigma Aldrich (ADV465749205-1G), and SGC-CK2-1 was sourced from Tocris Biosciences (745010). Natural products were sourced as follows: chrysin (Sigma Aldrich); TMF, dTMF, chrysoeriol, and homoeryodictyol (Indofine Chemical). Apigenin was a kind gift from the Mars Corporation. Naringenin was a kind gift from James LeClair. dO-TMF was synthesized from TMF according to the following conditions: in a procedure adapted from Xiao et al^8^, a flame-dried 25-mL teardrop flask was charged with a Teflon stir bar and aluminum chloride (80 mg, 0.6 mmol, 3 eq.). The vessel was sealed with a rubber septum, evacuated via a needle inlet, and backfilled with argon. The vessel was charged with 1.1 mL dry tetrahydrofuran, and the resultant suspension was cooled with stirring in an ice-water bath. Lithium aluminum hydride solution (2.5 M in tetrahydrofuran, 0.12 mL, 1.5 eq.) was then added dropwise. Ten min later, a solution of trimethoxyflavone (TMF, 63 mg, 0.2 mmol, 1 eq) in dry tetrahydrofuran (4 mL) was added dropwise over 5 min. The resultant yellow mixture was stirred in an ice-water bath for another 125 min; it was then diluted with wet ethyl acetate followed by dropwise addition of deionized water, poured into a separatory funnel, and the layers were separated. The aqueous layer was extracted twice more with ethyl acetate. The combined organic fractions were washed with water and brine, dried over sodium sulfate, filtered, and concentrated at reduced pressure. Flash-column chromatography (6 by 4 cm SiO_2_, 10% ethyl acetate/hexanes (250 mL)) of the crude residue generated 14 20 mL fractions. Fractions 6-12 were combined and concentrated to give dO-TMF as a white solid (38 mg, 0.128 mmol, 64%). 1H NMR (500 MHz, CDCl3, δ): 7.59 (d, J = 9 Hz, 2H); 6.90 (d, J = 9 Hz, 2H); 6.21 (d, J = 1 Hz, 1H); 6.15 (d, J = 1 Hz, 1H); 5.39 (t, J = 4 Hz, 1H); 3.83 (s, 3H); 3.80 (s, 3H); 3.79 (s, 3H); 3.34 (d, J = 4 Hz, 2H). TAL606 was prepared according to Tucker et al.^8^ Compounds were stored at -20°C as 10-20 mM stock solution aliquots in DMSO.

### Differentiation of astrocytes from hESCs and hiPSCs

H1 hESCs (WiCell Research Institute) or AD and control hiPSCs (UCI ADRC iPS Cell Bank) were cultured on Matrigel-coated plates in mTeSR medium and differentiated into glial progenitor cells (GPCs) and mature astrocytes as previously described^9^. First, EBs were prepared by mechanical dissociation of H1 cultures with 1 mg/mL collagenase IV (Gibco), plated onto ultra-low attachment plates (Corning) in TeSR medium with 10 uM Y-27632 (ROCK inhibitor, StemCell Tech), and incubated overnight with rocking. For differentiation of GPCs from EBs, medium was changed to AM supplemented with 500 ng/mL Noggin (R&D Systems) and 10 ng/mL PDGFAA (Peprotech) for 2 weeks and then Noggin was withdrawn for another week. The EBs were dissociated with papain (Papain dissociation system, Worthington) and the GPCs were cultured and expanded in 10 mg/mL poly-L-ornithine (Sigma)/1 mg/mL laminin (Invitrogen)-coated plates in AM supplemented with 20 ng/mL fibroblast growth factor 2 (FGF-2, Joint Protein Central) and 20 ng/mL epidermal growth factor (EGF, Humanzyme). Astrocytes were differentiated from low-confluent GPC cultures in DMEM/F12 Glutamax supplemented with N2 and B27 and 10% FBS. After 2 weeks of differentiation, the cells were transferred to non-coated plates for another 2 weeks of maturation. Immunofluorescence staining was performed as described below to assess astrocytic marker expression with antibodies for Glast (EAAT1): mouse anti-EAAT1 (1:1,000; Santa Cruz, sc-15316); S100b: rabbit anti-S100b (1:1,000; Dako, Z031129-2); Vimentin: goat anti-Vimentin (1:500; EMD Millipore, AB1620).

### Differentiation of microglia-like cells from hESCs and hiPSCs

H1 hESC and EC11^10^ or Clue4-7^11^ hiPSCs were cultured on Matrigel-coated plates in mTeSR medium and differentiated into mature microglia-like cells (iMGLs) via an induced hematopoietic progenitor cell (iHPC) intermediate, as previously described.^12^ First, confluent stem cell cultures were dissociated into single-cell suspensions with TrypLE (Gibco) and plated onto untreated tissue-culture plates in TeSR with 10 uM Y-27632. The next day, medium was changed to iHPC basal medium supplemented with FGF2 50 ng/ml, BMP4 50 ng/ml (Peprotech), Activin-A 12.5 ng/ml (Proteintech), 1 uM Y-27632, and 2 mM LiCl (Sigma), and placed in a hypoxic incubator (5% O_2_, 5% CO_2_) for 4 days. iHPC basal medium was composed of 50% DMEM/F12, 50% IMDM, 2% ITS-X Insulin-Transferrin-Selenium-Ethanolamine (Thermo-Fisher Scientific), L-ascorbic acid 2-phosphate magnesium (64 µg/ml; Sigma), monothioglycerol (400 uM), PVA (10 µg/ml; Sigma), Glutamax (1X), chemically defined lipid concentrate (1X), non-essential amino acids (NEAA; 1X), and 1% Penicillin/Streptomycin. Next, cells were cultured in normoxic conditions for another 6-16 days with fresh medium supplemented with FGF2 50 ng/ml, VEGF 50 ng/mL (Proteintech), TPO 50 ng/ml (Proteintech), SCF 10 ng/ml (Proteintech), IL-6 50 ng/ml (Proteintech), and IL-3 10 ng/mL (Proteintech) added every 2 days. The resulting iHPCs were collected and sorted for CD43+ staining by FACS. Next, the pure iHPCs were differentiated into iMGLs by a 25-day maturation in iMGL medium [50% DMEM/F12, 50% IMDM, 2% ITS-G Insulin-Transferrin-Selenium (Thermo-Fisher Scientific), 2% B27, 0.5% N2, 200 uM monothioglycerol, 5 µg/mL insulin (Sigma), Glutamax (1X), NEAA (1X), 1% Penicillin/Streptomycin (ScienCell)] supplemented with cytokines. For the first 22 days, cytokines M-CSF 25 ng/ml, IL-34 100 ng/ml, and TGFb-1 50 ng/ml (Proteintech) were added. Following a 3-day maturation with the addition of CD200 100 ng/ml and CX3CL1 100 ng/ml, the iMGLs were FACS-sorted using a 6-antibody panel. CD45+, CD11b+, CD64+, CD14+, CX3CR1+ and HLA-DR+ cells were plated onto 96-, 24- or 6-well Primaria plates (Corning) for downstream bead phagocytosis and activation assays.

### Flow cytometry

For each well of HCAs in a 6-well plate, 2 mL of fresh medium with GolgiPLUG (BD #555029, 1:1000) and GolgiSTOP (BD # 554724, 1:1000) with 20 uM compound or vehicle (DMSO) and IL1-β 10 ng/mL or DPBS was added. The cells were incubated for 5 h at 37°C. Cells were collected with TrypLE and plated into a 96-well V-bottom plate for staining. The plate was centrifuged for 5 min at 1,300 rpm, 4°C, to pellet cells that were subsequently resuspended in 100 µL DPBS with 1 µL Zombie Violet (Biolegend #423113) and 5 µL Human Trustain FC block (Biolegend #422302) per well for 15 min at RT in the dark. The plate was centrifuged, supernatant was aspirated, and the cells were washed with 100 µL/well DPBS. Cells were fixed with 100 µL/well of Cytofix/Cytoperm (BD) for 20 min at 4°C. The plate was centrifuged, supernatant was aspirated, and cells were resuspended in 100 µL Permwash (BD) and incubated for 15 min at 4°C. The plate was centrifuged, supernatant was aspirated, and cells were incubated with 100 µL/well of Ab mix or isotype mix in Permwash (see below) for 20 min at 4°C, followed by centrifugation and 2x PermWash washes. The cells were resuspended in 100 µL DPBS and transferred to FACS tubes containing 200 µL of PBS (total volume per tube 300 µL) for analysis on FACS Analyzers (LSRII or Fortessa).

| Antigen | Fluorochrome | Volume | Concentration | Isotype | Clone | Catalog # | Company |
| --- | --- | --- | --- | --- | --- | --- | --- |
| IL-8 | PerCP | 5 μl | 100 μg/ml | Mouse IgG2b | BH0814 | 514606 | Biolegend |
| TNF-a | Pe-Cy7 | 5 μl | 200 μg/ml | Mouse IgG1, k | MAB11 | 502930 | Biolegend |
| IL-6 | APC | 5 μl | 50 μg/ml | Rat IgG1, k | MQ2-13A5 | 501112 | Biolegend |
| MCP-1 (CCL2) | PE | 3 μl | 200 μg/ml | Armenian Hamster IgG | 2H5 | 505904 | Biolegend |

### FACS sorting

For FACS sorting of iHPCs and iMGLs, cells were collected and filtered through a 70-µM mesh cell strainer attached to a 50-ml conical tube to remove large clumps and centrifuged at 1,000 rpm for 3 min. The pellet was washed with 3 mL of cold sterile filtered FACS buffer (1X DPBS, 2% BSA, and 0.05 mM EDTA pH 8.0) and transferred to a 15-ml conical tube centrifuge. The pellet was resuspended in 100-300 µL of FACS buffer, 5 µL/100 µL of Trustain was added, and the cells were incubated at RT for 5 min. Five µL of cells were removed from the sample and transferred to a new conical tube and brought to 100 µL of FACS buffer, and 5 µL of corresponding isotype control antibody (see below) was added along with 1 µL of Zombie Violet live/dead stain. For each sample, 5 µL of each antibody (see below) and 1-3 µL of Zombie Violet live/dead stain was added. Each sample and the isotype control were incubated on ice in the dark for 20 min. After staining, the samples were washed once with 1 ml cold FACS buffer, centrifuged and resuspended in 100-300 µl FACS buffer. The sample and the isotype control were transferred into separate FACS tubes with a 40-µM filter attached. For iHPC, all live and CD43+ cells were sorted into cold basal iHPC differentiation medium including Pen/Strep. For iMGL, all live and CD45+, CD11b+, CD64+, CD14+, CX3CR1+ and HLA-DR+ cells were sorted into cold basal iMGL differentiation medium.

| Color | Antibody | Catalog # | Volume (μL) | Concentration (μg/mL) |
| --- | --- | --- | --- | --- |
| APC | CD64 | 305014 | 5 | 200 |
| PerCP-Cy5.5 | CX3CR1 | 341614 | 5 | 400 |
| 488/FITC | CD14 | 325610 | 5 | 400 |
| PE | CD11b | 301306 | 5 | 200 |
| PE-Cy7 | HLA-DR | 307616 | 5 | 200 |
| APC-Cy7 | CD45 | 304014 | 5 | 100 |
| FITC | CD43 | 315204 | 5 | 400 |
| FITC | Isotype Ct | 400107 | 5 | 500 |
| APC | Isotype Ct | 400122 | 5 | 200 |
| PerCP-Cy5.5 | Isotype Ct | 401623 | 5 | 200 |
| PE | Isotype Ct | 400112 | 5 | 200 |
| PE-Cy7 | Isotype Ct | 400231 | 5 | 200 |
| APC-Cy7 | Isotype Ct | 400127 | 5 | 200 |

### IL-6 and TNFa secretion assays

HCAs were plated at 20,000 cells per well in white, clear bottom 96-well plates. Two to 3 days later, medium was removed and replaced with medium containing 10 ng/mL IL1-β and compound dilution series. Five hr later, plates were processed using the Lumit IL-6 Assay Kit (Promega W6030) or the Lumit TNFa Assay Kit (Promega W6050) according to manufacturer’s instructions.

### NF-κB-Luciferase reporter assay

THP-1 NF-κB-Lucia cells were plated at 10,000 cells per well with 10 ng/mL PMA. Three days later, medium was removed and replaced with medium containing lipopolysaccharide (LPS 1 µg/mL) and dilution series of compounds in flat, clear bottom 96-well plates for 24 hr. A day later, 20 µL of medium was transferred to a 96-well white plate and 50 µL of QuantiLuc Gold reagent (InvivoGen rep-qlcg5) was added per well; the plate was mixed in the plate reader and luminescence was read using a Promega GloMax Reader.

### Cytokine release assay

Microglia were treated in 6-well plates with fresh iMGL medium and either vehicle alone (DMSO) or 200 ng/mL LPS (Sigma) with 20 µM CHR or vehicle for 5 hr in duplicate. Subsequently, medium cleared of cells and debris was collected, snap-frozen and stored at -80°C until processing with the Human Cytokine Array kit (R&D Systems, Catalog # ARY005). Briefly, medium was diluted in Array Buffer 4 and incubated with a cocktail of biotinylated detection antibodies for 1 hr at RT with rocking. The resulting immunocomplexes were incubated with nitrocellulose membranes spotted with capture antibodies overnight at 4°C (see below for full list of targets). The membranes were washed 4 times with Wash Buffer and then incubated with IRDye 800CW Streptavidin for 30 min at RT (1:2000 in Array Buffer 5, LI-COR #926-32230). The membranes were washed 4x and imaged with an Odyssey CLx instrument. Intensity for each spot was quantified with ImageStudio software (LI-COR). Technical duplicate values for each target were averaged, and we determined differential cytokine release by microglia exposed to LPS + vehicle versus vehicle only (n = 2 biological replicates, two-tailed t-tests, P < 0.05). Then we determined whether the release of those cytokines was significantly affected by CHR treatment (n = 2 biological replicates, 2-tailed t-tests, P < 0.05).

Targets probed by Human Cytokine Array kit (R&D Systems, Catalog # ARY005):

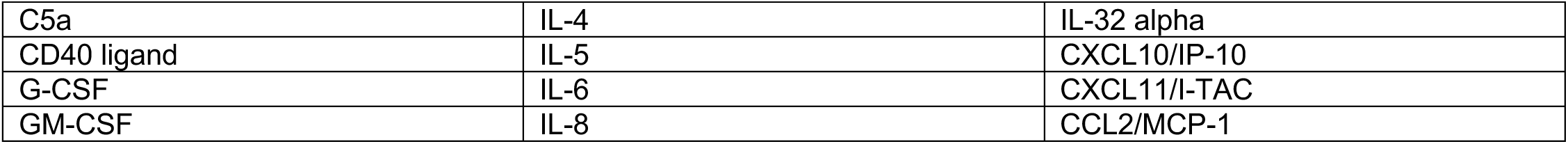
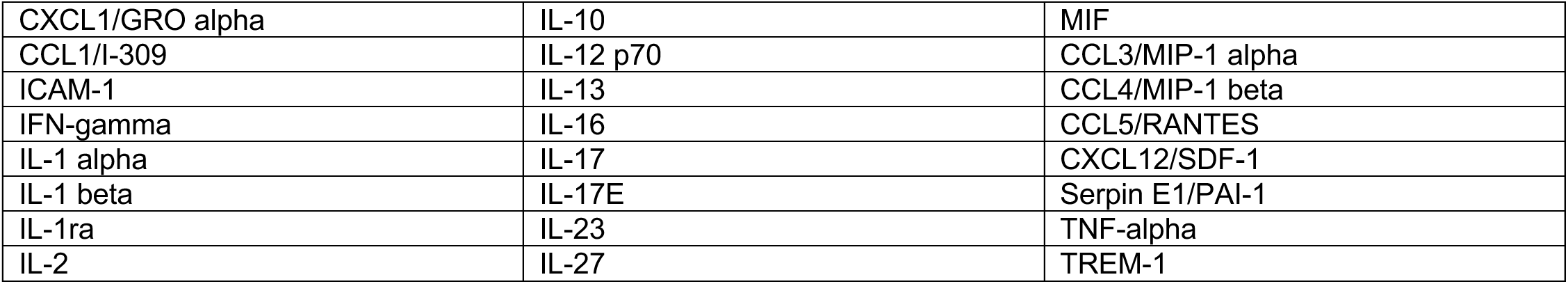

### Phagocytosis assay

Microglia were pre-treated in 12-well plates with fresh iMGL medium with vehicle (DMSO), 20 µM CHR, or 20 µg/mL cytochalasin D for 1 hr at 37 °C (2-3 biological replicates). Then, 0.5 µL of pHrodo E. Coli-FITC beads (Thermo Fisher, P35366) was added per well with gentle mixing and the cells were incubated for another 4 hr. pHrodo beads only fluoresce when they are internalized in cells. Cells were then collected by scraping, washed with FACS buffer (1X DPBS, 2% BSA, and 0.05 mM EDTA pH 8.0), and counter-stained with Zombie Violet to assess viability. Live cells were analyzed on the FACS Canto instrument for geometric mean fluorescence on the FITC channel. The background FITC fluorescence value (pHrodo beads only) was subtracted from each mean, and fold change was computed across experimental groups.

### Western blotting

For drug treatment assays, cells were treated with vehicle (DMSO) or 20 uM compound and stimulated with IL1-β 10 ng/mL for 1, 6, or 24 hr. Cells were dissociated with 1:1 Accutase:papain for 3 min at RT, pelleted for 5 min at 800 g, and washed twice with cold DPBS. Pellets were lysed with RIPA buffer (Sigma R0278) supplemented with Halt protease inhibitor cocktail (Thermo Fisher 78445) and clarified by centrifugation (16,000 g, 10 min, 4°C). Protein concentration was determined by BCA assay (Thermo Fisher). Proteins were denatured in 1X LDS Buffer and 2.5% beta-mercaptoethanol (Sigma) for 10 min at 70°C, separated on Bolt 4-12% Bis-Tris Plus polyacrylamide gels (Thermo Fisher), transferred to PVDF membranes using the iBlot 2 Dry Blotting System, and blocked with 0.1% casein-PBS (Bio-Rad) for 1 hr at RT. The blots were incubated with primary antibodies overnight at 4°C in PBS containing 0.1% casein and 0.2% Tween-20. Antibodies used were rabbit anti-CK2α1 (Bethyl #A300-198A, 1:5000), rabbit anti-CK2α2 (Bethyl #A300-199A, 1:5000), mouse anti-GAPDH (Fitzgerald #10-1501, 1:20000), rabbit anti-actin (Sigma #A2066, 1:20000), mouse anti-NF-κB (CST #6956, 1:1000), pNF-κB S529 (ThermoFisher #14-9864-82, 1:100), mouse anti-IkBa (CST #4814, 1:1000), mouse anti-IkBa pS32 (Santa Cruz #8404, 1:500), mouse anti-PTGR1 (Abnova #10563-436, 1:1000), and anti-HA-tag (Bethyl #A190-108A, 1:20000), rabbit anti-AKT pS129 (ab133458, 1:1000), mouse anti-AKT1 (Santa Cruz #sc-55523, 1:1000), mouse anti-synaptotagmin antibody (Abcam #ab13259, 1:1000) and rabbit anti-synaptophysin (Abcam #ab32594, 1:5000). The membranes were washed 5 times with 0.1% PBS-Tween20 (PBST) and incubated with secondary antibodies (1:20,000) for 1 hr at RT in PBS with 0.1% casein, 0.2% Tween20, and 0.1% SDS. Secondary antibodies used were goat anti-mouse IgG IRDye 800CW #925-32210, goat anti-rabbit mouse IgG IRDye 800CW #925-32211, goat anti-mouse IgG IRDye 680RD #926-68070, or goat anti-mouse IgG IRDye 680RD #926-68071. The blots were then washed 4 times with PBST and once with DPBS. Amide Black was used the stain the blots for total protein content following immunostaining. The blots were imaged using the Odyssey CLx imaging system and blots were analyzed on ImageStudio software (LI-COR).

### Immunofluorescence staining of primary cells

Primary human astrocytes (ScienCell) were plated on glass slides (EMD Millipore PEZGS0896) and treated with CHR, CX-4945/silmitasertib, or DMSO and stimulated with IL1-β for 1, 5 or 25 hr. After treatment, cells were fixed with 4% paraformaldehyde solution for 20 min at RT and washed 3 times with DPBS for 10 min each. The cells were permeabilized using 5% horse serum and 0.1% Triton X in DPBS for 15 min at RT and blocked with 5% horse serum in PBS for 30 min at RT. Cells were incubated with primary antibodies in blocking buffer either overnight at 4°C or for 2 hr at RT. After incubation, cells were washed 3 times with PBS, blocked using 5% horse serum (Sigma H1270) in PBS for 30 min at RT, and incubated with Cy3 red-labeled or AF488 green-labeled secondary antibodies diluted in blocking buffer for 1 hr at RT. The cells were washed once with PBS for 10 min and then counter-stained with DAPI. The cells were washed 3 times with PBS and mounted with glass coverslips and Immu-Mount solution. Images were obtained at 20X with a Zeiss fluorescent microscope (Axio Observer Z1), visualized using Zen Blue v3.3 and ImageJ v2.1.0, and analyzed using CellProfiler v3.1.8. The following primary antibodies were used: rabbit anti-NF-κB (1:400; CST #8242), mouse anti-phospho NF-κB S529 (1:25; R&D MAB7624), rabbit anti-CK2Α1 (1:500; Bethyl A300-198A), and rabbit anti-phospho CK2Α1 Y255 (1:25; Thermo Fisher PA5-40226).

### Activation assays and ddPCR

HCA or H1-derived astrocytes were treated with vehicle (DMSO) or 20 µM compound and stimulated with IL1-β 10 ng/mL for 5 hr. Cells were scraped into RNA-Bee and total RNA was purified using the Direct-zol RNA kit (Zymo Research). RNA concentration was measured using Nanodrop and RNA was reverse transcribed into cDNA using the High Capacity cDNA Reverse Transcriptase Kit (Applied Biosystems). Reaction mixes consisting of TaqMan FAM probe (IL6 probe Hs00174131_m1, IL8 probe Hs00174103_m1, CSNK2A1 probe Hs00751002_s1, CSNK2A2 probe Hs00176505_m1), control TaqMan ACTB-VIC probe (Thermo Fisher Scientific 4326315E), ddPCR Supermix for Probes (Bio-Rad 186-3024), and cDNA were formed into oil droplets using the QX200 Droplet Generator. After amplification (Bio-Rad C1000 Touch Thermal Cycler) according to manufacturer’s instructions, the plate was read in the QX200 Droplet Reader.

### Activation assays and qRT-PCR

AD or CT patient-derived astrocytes were treated and RNA was collected, purified, and reverse transcribed as described in the “Activation assays and ddPCR” section. Reaction mixes consisting of TaqMan FAM probe (IL6 probe Hs00174131_m1, TNFAIP2 probe Hs00969305_m1), control TaqMan ACTB-VIC probe (Thermo Fisher Scientific 4326315E), 2x TaqMan Fast Advanced Master Mix (Applied Biosystems), and cDNA were run in technical triplicate on a QuantStudio 3 or 5 Flex Real-Time PCR System (Thermo Fisher). Data was analyzed using the Realtime PCR Relative Quantification module (Thermo Fisher Cloud Software v1.1).

### Cloning and lentiviral transduction

pBOB-CAG-CK2α1-WT, pBOB-CAG-CK2α2-WT, pBOB-CAG-CK2α1-K68M, and pBOB-CAG-CK2α1-K69M plasmids were generated by cloning HA-tagged CK2 inserts from parent plasmids (Litchfield lab, Addgene 27086, 27090, 27089, 27087) into the pBOB backbone (Addgene #12337). pBOB-CAG-CK2α2-P2A-GFP-PGKpuro and pBOB-CAG-P2A-GFP-PGKpuro were generated by cloning in a P2A sequence and an eGFP-PGKpuro insert from a pCSC-hProx1-5-eGFP-PGKpuro construct (Gage lab) using In-Fusion HD Cloning Kit (Takara Bio 639642). Plasmids were transformed into TOP10 competent cells (Thermo Fisher) or Stellar competent cells (Takara Bio). H1-GPCs in 6-well plates were transduced with lentiviral particles (∼10^8^ particles/mL, Salk Virus Core) by incubating for 3 days. Medium was changed and cells were differentiated into mature astrocytes as described above after checking for HA-CK2 expression by Western blotting. Astrocytes were processed as described under “Activation assays, ddPCR.”

### siRNA knockdown

HCA were nucleofected with siRNAs against *CSNK2A1* (Ambion s2888), *CSNK2A2* (Ambion s7501), or scrambled control (Ambion 4390843) using the Amaxa Nucleofector II (Program T-019). After 48 hr, knockdown efficiency was assessed by ddPCR and cells were replated to assess inflammatory response to IL1-β as described under “Activation assays, ddPCR.” Results shown in Figure 2D represent multiple electroporations.

### Radiometric kinase assay

Compounds dTMF, dOTMF, and dCHR were tested for *in vitro* inhibition of CK2α1 (Eurofins Discovery Services, Catalog #14-197KP). CK2 (h) was incubated with 20 mM HEPES pH 7.6, 0.15 M NaCl, 0.1 mM EDTA, 5 mM DTT, 0.1% Triton X-100, 165 uM RRRDDDSDDD, 10 mM MgAcetate and 15 µM [gamma-33P]-ATP in the presence of DMSO or compound. Compounds were tested in 8-point dilution series starting at 10 µM (n = 2), starting from 50X stocks in DMSO. Each reaction was initiated by the addition of the Mg/ATP mix. After incubation for 40 min at RT, each reaction was stopped by the addition of phosphoric acid to a concentration of 0.5%. An aliquot of each reaction was then spotted onto a filter and washed 4 times for 4 min in 0.425% phosphoric acid and once in methanol prior to drying and scintillation counting.

### CRISPR-Cas9 editing

THP-1-NF-κB-Lucia cells were genome-edited using CRISPR-Cas9 ribonucleotide particles (RNP). Two million cells were washed with DPBS and resuspended in 100 µL of Ingenio Electroporation Buffer (Mirus Bio 10766-842). For the CK2 knockouts, custom-order TrueGuide synthetic gRNAs by Thermo-Fisher (A35533) were used: sgCK2Α1 (sequence: GTGAGGATAGCCAAGGTTCT) and sgCK2Α2 (sequence: ACGCCGAGGTGAACAGTCTG). For the NF-κB S529A-NeoR knockin, 2 custom-order TrueGuide synthetic gRNAs by Thermo-Fisher (A35533) were used: sgNF-κB-G526 (sequence: ATCTCCTGAAAGGAGGCCAT) and sgNF-κB-stopCodon (sequence: AGGGCAGGCGTCACCCCCTT). For each CRISPR-Cas9 editing experiment, 8 ug of Cas9 (TrueCut V2, Thermo-Fisher A36498) was mixed with 0.48 µL of each sgRNA (100 µM stock) and incubated 15 min at RT. The RNP complex was mixed with 0.48 µL electroporation enhancer (IDT 1075916) and added to cells in Ingenio Electroporation Buffer. For the knock-in experiment, 9 µL HDR donor (HDR-NF-κB-WT-NeoR-v2, 325 ng/µL) was also added at this step. HDR donor DNA was synthesized by Synbio Technologies. The cells were nucleofected using program V-001 on the Amaxa II Nucleofector and then added to 2 mL THP-1 medium (plus HDR enhancer v2 at 1 µM (IDT 10007910, 1:690) for the knockin experiment). After 3 weeks recovery, cells were selected with 100 µg/mL G418 (Gibco 10131027) for 1 week. For CK2Α1 and CK2Α2 knockouts, clones were isolated using FACS sorting into 96-well plates and validated by Western blotting, immunoprecipitation-Western, and/or Sanger sequencing.

### Thermal shift assay and thermal proteome profiling (TPP)

Experiments were performed as previously described.^13,14^ Forty-five million H1-derived astrocytes were activated with 10 ng/mL IL1-β for 6 hr, dissociated with 1:1 Accutase/papain (3 min RT), and centrifuged at 1,800 rpm for 2 min at 4°C. The pellet was washed twice with 10 ml cold DPBS, lysed in 2.25 mL DPBS and 0.4% v/v NP-40 by freeze-thawing 3x, and clarified by centrifugation (20,000*g*, 30 min, 4°C). Protein concentration was determined by BCA assay (Bio-Rad), and lysates were diluted to 2 mg/mL. First, compound or vehicle was added (10 µL DMSO, 10 µL of 20 mM TMF, or 10 µL of 20 mM CHR) to individual tubes, then 0.99 mL lysate was added to each tube and vortexed briefly. Final concentrations were 1% vehicle, 200 uM TMF, and 200 uM CHR. Reactions were incubated at RT for 30 min, after which each treated extract was divided into 10 aliquots of 95 µL in PCR strip tubes and stored on ice. Each group of treated samples was heated for 3 min at predesignated temperatures (3 x 10 temperatures) and then kept for 3 min at RT before ultracentrifugation (20 min at 4°C, 100,000*g*) in 200 µL polycarbonate tubes (Beckman Coulter #343775). Designated temperatures were 37, 41.1, 43.6, 46.9, 50, 53.7, 56, 59.5, 63, and 67°C. A small aliquot (5 µL) of supernatant was saved for Western blotting and the rest was snap frozen at -80°C and submitted for LC-MS/MS. For temperature gradient thermal shift experiments with TMF and dTMF (Extended Figure 2F), a similar procedure was followed except that samples were directly processed for Western blotting. For isothermal dose-response experiments, CHR was added to individual extracts at concentrations in a 3-fold dilution series ranging from 200 μM-0.1 μM, including one vehicle control, and the extracts were heated at 60°C and processed as described above.

### Mass spectrometry for TPP study

Samples were precipitated by methanolchloroform and redissolved in 8 M urea/100 mM TEAB, pH 8.5. Proteins were reduced with 5 mM tris(2-carboxyethyl)phosphine hydrochloride (TCEP, Sigma-Aldrich) and alkylated with 10 mM chloroacetamide (Sigma-Aldrich). Proteins were digested overnight at 37°C in 2 M urea/100 mM TEAB, pH 8.5, with trypsin (Promega). The digested peptides were labeled with 10-plex TMT (Thermo product 90309) and pooled samples were fractionated by basic reversed phase (Thermo 84868). The TMT-labeled samples were analyzed on a Fusion Lumos mass spectrometer (Thermo) controlled by Thermo Xcalibur version 4.0.27.10. Samples were injected directly onto a 25 cm, 100 μm ID column packed with BEH 1.7 μm C18 resin (Waters). Samples were separated at a flow rate of 300 nL/min on a nLC 1200 (Thermo). Buffer A and B were 0.1% formic acid in water and 90% acetonitrile, respectively. The following conditions were used for 240 min of total run time: a gradient of 1–25% B over 180 min, an increase to 40% B over 30 min, an increase to 100% B over 20 min and a hold at 90% B for 10 min. Peptides were eluted directly from the tip of the column and nanosprayed directly into the mass spectrometer by application of 2.8 kV voltage at the back of the column. The Lumos was operated in a data-dependent mode. Full MS1 scans were collected in the Orbitrap at 120k resolution. The cycle time was set to 3 sec, and within these 3 sec the most abundant ions per scan were selected for CID MS/MS in the ion trap. MS3 analysis with multinotch isolation (SPS3) was utilized for detection of TMT reporter ions at 60k resolution^15^. Monoisotopic precursor selection was enabled and dynamic exclusion was used with exclusion duration of 10 sec. Protein and peptide identification were done with Integrated Proteomics Pipeline – IP2 (Integrated Proteomics Applications, version 6.0.1). Tandem mass spectra were extracted from raw files using RawConverter^16^ version 1.1.0.18 and searched with ProLuCID^17^ version 1.4.2 against the Uniprot human database. The search space included all fully tryptic and half-tryptic peptide candidates. Carbamidomethylation on cysteine and TMT on lysine and peptide N-term were considered as static modifications. Data were searched with 50 ppm precursor ion tolerance and 600 ppm fragment ion tolerance. Identified proteins were filtered using DTASelect^18^ version 2.1.4 and utilizing a target-decoy database search strategy to control the false discovery rate to 1% at the protein level^19^. Quantitative analysis of TMT was done with Census^20^ version 2.51, filtering reporter ions with 20 ppm mass tolerance and 0.6 isobaric purity filter.

### TPP analysis

Mass spectrometry data were analyzed using the TPP-temperature range (TR) workflow of the R package *TPP* (version 3.10.0) as previously described.^13^ The package performs protein quantity normalization, fits melting curves, determines melting points, and identifies proteins that have a significant shift in thermal stability compared with controls. We used data from 2 independent TPP experiments to quantify significant thermal shifts between CHR and vehicle and TMF and vehicle. The algorithm was run with minor changes to default parameters (filtering requirements changed to include peptides with spectral counts ≥ 2 and fold-change thresholds from 0-1.5). Briefly, data for each protein and condition underwent curve-fitting analysis, and the significance of thermal shift was calculated only for proteins with R^2^ >0.8 and a plateau of <0.3 for the vehicle curve. Proteins that met all 4 of the following benchmarks for significance and had spectral counts ≥ 3 were considered “hits”: (a) P values for the 2 replicate experiments were <0.05 and <0.2, respectively; (b) the compound-vehicle melting point shifts in the 2 independent experiments had the same direction; (c) each compound-vehicle ΔT_m_ was greater than the ΔT_m_ between the 2 vehicle controls; and (d) the minimum curve slope in each experiment was <−0.06. Network analysis and visualization of each set of hits were performed with the GeneMania app v3.4.1 within Cytoscape (v. 3.5.1).

### Native kinase capture

Experiments were performed as previously described.^21^ Forty-five million H1-derived astrocytes were activated with 10 ng/mL IL1-β for 6 hr, dissociated with 1:1 Accutase/papain (3 min RT), and centrifuged at 1,800 rpm for 2 min at 4°C. The pellet was washed twice with 10 ml cold DPBS, lysed in 2.25 mL DPBS by repeated freeze-thawing, sonicated (2 × 10 s pulses with a 30-sec break, 4°C) and clarified by centrifugation (20,000*g*, 10 min, 4°C). The supernatant was desalted through a column (732-2010, Biorad) and then eluted with cold kinase buffer (20 mM HEPES, pH 7.4, 150 mM NaCl, 0.5% Triton X-100, with Halt Protease and Phosphatase inhibitor cocktail). For each treatment, 475 μl of the lysate (2 mg/mL) was pre-incubated with 10 μl MnCl_2_ (1 M) and 5 μl compound to the desired concentration at RT for 30 min. Uninhibited kinases were captured with 10 μl ActivX desthiobiotin-ATP probe (0.25 mM; 88311, Pierce) at RT for 10 min. Samples were mixed with 500 μl urea (8 M; 818710, Millipore) and 50 μl streptavidin agarose (20359, Thermo) for 60 min at RT on a nutator. Beads were washed twice with a 1:1 mixture of kinase buffer and 8 M urea and were collected by centrifugation (1,000*g*, 1 min). Proteins were eluted from the beads with 100 μl 2 × LDS sample buffer (NP0007, Life) at 95°C for 10 min. Samples were analyzed by standard immunoblotting and normalized to total protein following Amido Black staining. The experiment was performed twice.

### Mass spectrometry for CHR brain exposure study

Target compound was extracted from tissue lysate (brain, 25 mg of tissue) with ethyl acetate spiked with d5-apigenin as internal standard. Samples were vortexed and centrifuged (14,000 x g, 4°C, 10 min). The organic layer was transferred to a new tube and dried in speedvac. Samples were reconstituted in 50 µL methanol prior to injection. A compound standard curve was created in serum using the same extraction protocol. Samples were analyzed on a TSQ Quantiva mass spectrometer (Thermo), controlled by Thermo Xcalibur 3.0.63, coupled to a Dionex Ultimate 3000 LC system (Thermo) fitted with a Luna C18(2) column (3 µm, 250 x 2 mm i.d., Phenomenex). The following LC solvents were used: solution A, 0.1 % formic acid in water; solution B, 0.1 % formic acid in acetonitrile. At a flow rate of 0.4 mL/min, the following gradient was used: column equilibrated at 6 % B, 6-16.5 % B in 5 min, 16.5-18.5 % B in 7.5 min, 18.5-100 % B in 8.5 min, 100 % B for 2 min and reequilibrate at 6 % B for 2 min, for a total run time of 25 min. The sample injection volume was 10 µL, column oven temperature was set to 30 °C and the autosampler kept at 4 °C. MS analyses were performed using electrospray ionization in positive mode, spay voltages of 3.5 kV, ion transfer tube temperature of 325 °C, and vaporizer temperature of 275 °C. Multiple reaction monitoring (MRM) was performed by using precursor mass of the intact compound (301.1 m/z) and five transitions (135.1, 203.0, 229.1, 258.1 and 286.1 m/z). Peak areas at retention time of 18.2 min were measured using Skyline^22^ version 21.2.0.536 and normalized to the internal standard. Compound amount was calculated using the standard curve.

### Immunoprecipitation

HCAs were treated for 2 hr with vehicle or CHR 20 µM and IL1-β 10 ng/mL or vehicle only. Cells were dissociated with Accutase and centrifuged at 1,800 rpm for 5 min at 4°C. The pellet was washed twice with cold DPBS, lysed in IP Buffer (20 mM HEPES 7.4 pH, 150 mM NaCl, 1% Triton-X, 1 mM EDTA, 1 mM EGTA plus Halt Protease and Phosphatase Inhibitor Cocktail) with sonication (6 × 10 s pulses with 10-sec breaks, 4°C) and clarified by centrifugation (20,000*g*, 10 min, 4°C). Lysates were precleared with Protein A/G Magnetic Beads (Pierce) for 20 min at RT and then incubated with NF-κB primary antibody (Novus NB100-97831, Rabbit polyclonal) overnight with rocking at 4°C. After 3 washes in cold DPBS, proteins were eluted in 4X LDS/BME at RT for 10 min and then denatured at 70°C for 10 min, separated on NuPAGE gels at 1X LDS, and probed with NF-κB primary antibody (CST D14E12) and anti-rabbit Light Chain-specific IgG AlexaFluor 790 (Jackson Immuno NC0493011) secondary antibody for detection. THP-1 NF-κB Lucia CRISPR clones were collected, washed twice in cold DPBS, lysed in IP Buffer, and processed as above, except for pulldown and blotting using anti-CK2A2 primary antibody (Bethyl A300-199A).

### RNA-seq in primary and iPSC-derived astrocytes

RNA-sequencing analysis was performed on 5-week-old HCAs, either non-stimulated or stimulated with 10 ng/mL IL-1β, and further split into 2 groups, without or with apigenin (20 µM). Five hours later, the cells were harvested in RNABee solution (Tel test, Inc) and total RNA was extracted using the DNAase-Free RNA Kit (Zymo Research) according to the manufacturer’s instructions. iPSC-derived astrocytes were differentiated for 4 weeks from GPCs in SATO medium (50% DMEM Glutamax Medium, 50% Neurobasal Medium, SATO (100x), 1 mM Glutamine (200x), 1 mM Sodium Pyruvate (100x), 20 ng/ml CNTF, 20 ng/ml BMP4, 5 µg/ml insulin, 20 ng/ml EGF, 20 ng/ml FGF2) in 6-well plates (triplicate wells, n = 3 CT, n = 3 AD lines). SATO supplement (100x) was prepared by adding transferrin 100 µg/ml, BSA 100 µg/ml, putrescine 16 µg/ml, progesterone 60 ng/ml, sodium selenite 40 ng/mL into Neurobasal Medium. Medium was changed 2 days before harvesting RNA to the following: 50% DMEM (no phenol red), 50% Neurobasal (no phenol red), 20 ng/mL carrier-free CNTF, 20 ng/mL carrier-free BMP4, 1 mM sodium pyruvate, 1 mM Glutamax, and 100 µg/mL Penn/Strep. RNA quality was assayed using Agilent Technologies 2200 TapeStation and only samples with high RNA quality (RIN > 8) were used for library preparation. Stranded mRNA-Seq libraries were prepared using the Illumina TruSeq Stranded mRNA Library Prep Kit according to the manufacturer’s instructions and single-end reads were acquired on an Illumina HiSeq 2500 or 4000 instrument.

### RNA-seq analysis of primary astrocytes

Sequenced reads were quality-tested using FASTQC^23^ and aligned to the hg19^24^ human genome using the STAR^25^ aligner version 2.4.0k. Mapping was carried out using default parameters (up to 10 mismatches per read, and up to 9 multi-mapping locations per read). The genome index was constructed using the gene annotation supplied with the hg19 Illumina iGenomes^26^ collection and overhang value of 100. Raw gene expression was quantified across all gene exons (RNA-Seq) using the top-expressed isoform as proxy for gene expression. Differential expression was performed using DESeq2^27^ package version 1.20.0. Differentially expressed (DE) genes were defined as having a false discovery rate (FDR) <0.05 and a log_2_ fold change >1. Enrichment analysis was performed using WebGestalt^28^ R package version 0.1.1. Hierarchical clustering was performed using the R language (v.3.3.2) with Ward’s hierarchical agglomerative clustering method and 1-correlation as a distance metric. GO enrichment analysis was performed using DAVID v6.8 and Kinase Enrichment Analysis (KEA^29^) was performed using X2K Web^29,30^ and the output Expression2Kinases network of enriched kinases and TFs resulting from all DE genes were visualized using Cytoscape^31^. Gene expression analysis for astrocyte polarization was performed by first extracting genes induced >4X after MCAO in astrocytes (“Astro-Injury”) or LPS (“Astro-Inflammation”) from Table 3 published by Zamanian et al (2012)^32^ Then, “Astro-Injury” genes were annotated as genes included in the MCAO set but excluded from the LPS set, whereas “Astro-Inflammation” genes were annotated as genes included in the LPS set but excluded from the MCAO set, as described in Figure 7 from Zamanian et al (see Supplementary Figure 4). Figure 3H was generated by (a) plotting relative expression of all “Astro-Injury” genes significantly downregulated in the IL1-β group relative to the PBS group (FPKM > 1; expression for each gene normalized to mean of PBS group) or (b) plotting relative expression of all “Astro-Inflammation” genes (FPKM > 1) significantly upregulated in the IL1-β group relative to the PBS group (FPKM > 1; expression for each gene normalized to mean of PBS group).

### RNA-seq analysis of iPSC-derived astrocytes

Sequenced reads were quality tested using FASTQC, reads were trimmed using SolexaQA^33^ v3.1.7.1 and adapters were removed using cutadapt^34^ v2.10. Reads were aligned to the hg38 human genome and quantified using Salmon^35^ v1.2.1 using default parameters in mapping-based mode. The pre-computed genome index with partial decoy was downloaded from https://refgenomes.databio.org/v2/asset/hg38/salmon_partial_sa_index/archive?tag=default. Quantified gene expression data were loaded into R v4.0.0 and annotated to Ensembl identifiers using package tximeta package v1.8.3 and converted gene symbols with package org.Hs.eg.db v3.12.0. Differential expression was performed using the glm function of edgeR^36^ package version 3.11. AD-upregulated genes were defined as having a false discovery rate (FDR) <0.05 and a log_2_ fold change >0.69. Enrichment analysis was performed using DAVID v6.8 and ClueGO v2.5.7 (Cytoscape^31^ 3.5.1 plugin). KEA^29^ was performed using Expression2Kinases 1.6.1207^29^, the list of top 20 kinases enriched in AD-upregulated genes was input into STRING^37^ and the output network was visualized using Cytoscape.

### Cmap analysis

L1000 data were accessed using clue.io cloud-based software (cMAP v v1.0.2). The list of DE genes from API/IL1-β-treated astrocytes compared to IL1-β-treated astrocytes was uploaded to CLUE, then searched against overexpression and knockdown signatures present in L1000 data using the Morpheus module. The resulting signatures were rank-ordered and plotted in GraphPad Prism from least to most similar.

### Analysis of aging, dementia, and traumatic brain injury (TBI) RNA-seq data and UPP proteomics data

Normalized RNA-seq data (as z-scores) for *CSNK2A1*, *CSNK2A2*, *CSNK2A3* and *CSNK2B* expression, GFAP IHC expression, and sample metadata from parietal cortex tissues were downloaded from https://aging.brain-map.org/download/index. Data were filtered to retain only samples from donors who had never experienced TBI and who were classified as having “Dementia” (n = 21) or “No Dementia” (n = 27). Individual two-tailed t-tests were conducted to determine statistically significant differences; only *CSNK2A2* exhibited differential expression between dementia and controls. For the GFAP comparison, raw values of GFAP IHC staining were plotted against z-scores for *CSNK2A2* for each corresponding donor tissue. For the UPP Proteomics study, normalized and batch-corrected log2(abundance) data and sample metadata for CSNK2A1, CSNK2A2, CSNK2A3 and CSNK2B were downloaded from https://www.synapse.org/#!Synapse:syn17009177. Data were filtered to retain only samples from patients with AD (n = 44) or age-matched controls (n = 48). One-way ANOVA with Sidak’s post-hoc test was conducted to determine statistically significant differences; only CSNK2A2 exhibited differential expression between AD and controls, and CSNK2A2 elevation relative to CSNK2A1 was observed in AD but not in controls. For the Braak scores comparison, Braak scores were plotted against CSNK2A2 log2(abundance) values for each corresponding donor tissue.

### NanoBRET assays

Assays were run using the NanoBRET Intracellular TE Kinase Assay K-5 kit (Promega #NV1191) according to manufacturer’s instructions. Approximately 1M HEK293T cells were transfected with 2.5 ug of NanoLuc-CSNK2A2 plasmid (Promega #PRN2500) using Lipofectamine 2000 and incubated for 24 hr. Cells were dissociated with TrypLE 3 min at RT, inactivated by adding medium, and spun down at 300 g for 3 min. Twenty thousand cells were seeded per well in DMEM + 10% FBS medium into white, non-binding surface, polypropylene 96-well plates and left to attach overnight. Test compounds were prepared by diluting 1,000X stocks to 1X final concentration in Assay Medium and by performing 2-fold serial dilutions. NanoBRET tracer was added to 1X final concentration to all compound concentrations. Medium was aspirated and a 100 μl of 1X serially diluted test compound and K-5 Tracer were added per well of the plate. The plate was incubated at 37°C, 5% CO_2_ for 2 hours. 3X Complete Substrate plus Inhibitor Solution in Assay Medium (Opti-MEM I Reduced Serum Medium, no phenol red) was prepared by gently mixing a 1:166 dilution of NanoBRET Nano-Glo Substrate plus a 1:500 dilution of Extracellular NanoLuc Inhibitor in Assay Medium. Fifty μl of 3X Complete Substrate plus Inhibitor Solution was added to each well of the 96-well plate and incubated for 2–3 min at RT. Donor emission (450 nm BP) and acceptor emission (600 nm LP) wavelengths were measured using the GloMax Discover System.

### Isothermal dose response target engagement study

HEK293T cells transiently expressing CSNK2A1- or CSNK2A2-Nanoluc were prepared as described above and seeded in suspension in Opti-MEM with 1% FBS in nonbinding V-bottom 96-well plates (Greiner Bio-One, #89131-674). Equal volumes of DMSO stocks or compound dilution series (2X, in Opti-MEM with 1% FBS) were added in triplicate and plates were incubated for 1h in the incubator. Cells were collected by centrifugation at 340 g and 4 °C for 2 min in each orientation. Cell pellets were washed twice with cold DBPS, then resuspended in DPBS supplemented with protease inhibitors. Samples were transferred to 0.1 mL PCR plates, and the PCR plates were heated in a PCR block for 3 min to 50 °C (for CK2A2) or 54 °C (for CK2A1) and then incubated for 3 min at RT. Ice-cold 4X lysis buffer (DPBS, 3.2 % Igepal CA-630, 8 U/mL benzonase, 6 mM MgCl_2_, cOmplete Mini EDTA-free Protease Inhibitor Cocktail, Roche #11836170001) was added to samples and the plates were incubated for 1 hr at 4°C. After clarification (3 min at 500 g), the soluble protein fraction was retrieved by centrifugation of supernatant for 5 min at 500 g through a 96w-filter plate (MultiScreen HTS HV MSHVN4550). Samples were transferred to white 96-well plates and Nano-Glo Luciferase Assay master mix (Promega N1110) was added. Luminescence was measured after a 3-min incubation. Stabilization fold-change for each protein was determined by normalizing luminescence values to DMSO controls.

### Statistical analyses

In general, experiments were repeated independently at least 3 times unless otherwise specified. Student’s t-tests were performed to compare 2 groups, whereas multiple-group comparisons were performed using a one-way or mixed effects analysis of variance (ANOVA) test with a post-hoc multiple comparisons test. Simple linear regression was performed to determine correlation statistics for two-variable datasets. Non-linear regression ([inhibitor] vs. response or normalized response, variable slope) was used to fit dose-response curves. Hypergeometric tests were performed to compare overlaps between two gene sets. A p-value less than 0.05 was considered statistically significant. Analyses were done using GraphPad Prism.

---

**Extended Data Figure 1.**
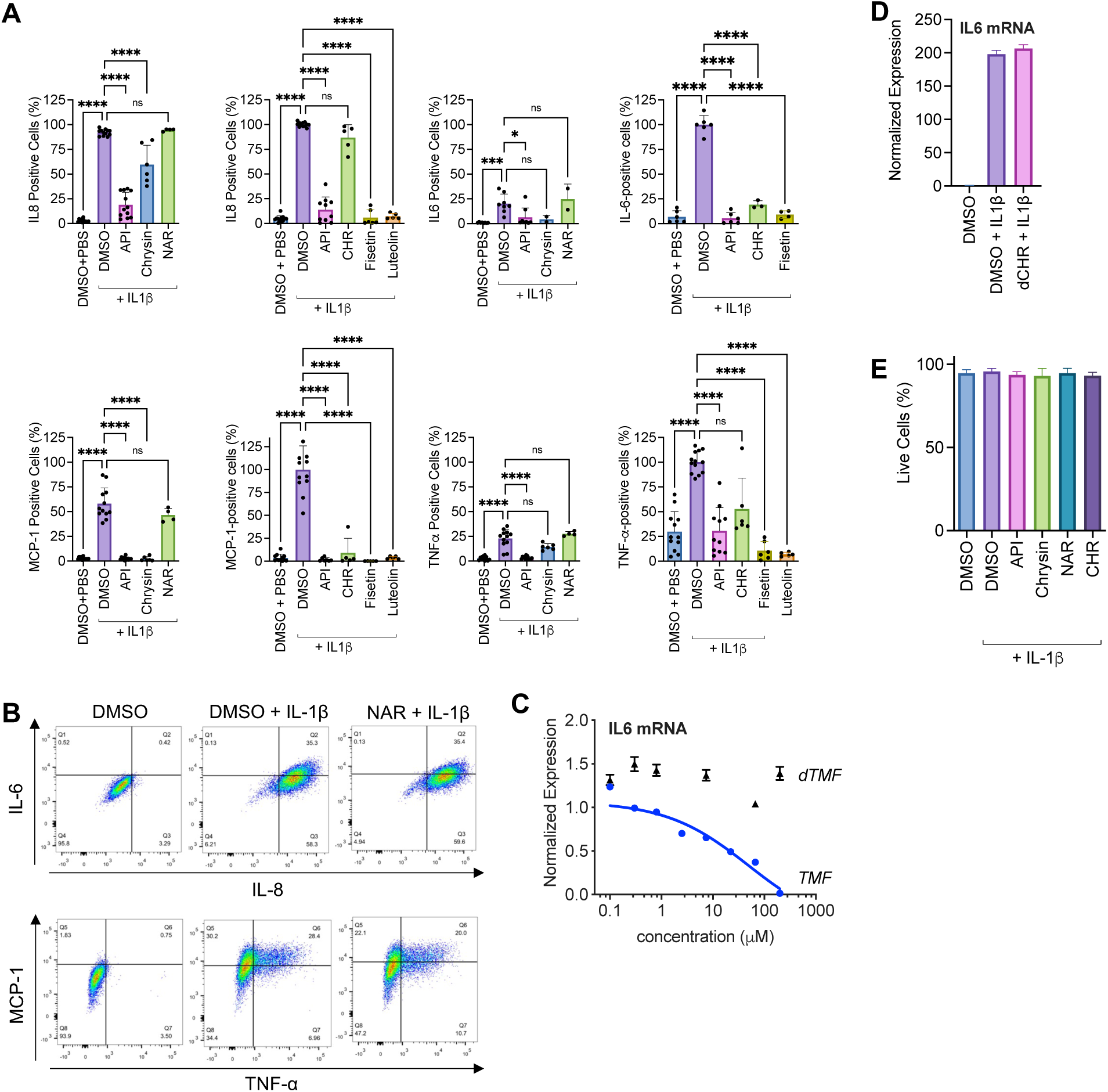
Additional characterization of flavones for anti-inflammatory activity and cytotoxicity in astrocytes (5-hour treatment). A. SAR studies show divergent activity of flavones and flavanones in a flow cytometry-based assay of cytokines production. Each dot represents 1 biological sample, and each bar graph is a pool of 2-5 independent experiments. For IL-8, MCP-1, and TNFα left graphs, n = 12 for DMSO + PBS, DMSO and API groups, n = 6 for chrysin group, n = 4 for NAR group; for IL-8 right graph, n = 12 for DMSO + PBS and DMSO groups, n = 10 for API, n = 6 for fisetin and n = 5 luteolin and CHR; for IL-6 left graph, n = 6 for all groups except n = 2 for NAR and chrysin groups; for IL-6 right graph, n = 6 for all groups except n = 3 for CHR and n = 4 for fisetin; for TNFα right graph, n = 12 for DMSO + PBS, n = 13 for DMSO, n = 11 for API, n = 6 for fisetin and CHR groups, n = 5 for luteolin; for MCP-1 right graph, n = 12 for DMSO + PBS and DMSO groups, n = 7 for API, n = 5 for fisetin, luteolin, and CHR groups; mean ± SD, one-way ANOVA with Bonferroni post-hoc test). Bar charts for NAR and chrysin in the IL6 graph represent 1 experiment. B. NAR does not block IL6, IL8, MCP-1, and TNFα cytokine production in HCA. Representative flow cytometry traces from 2 independent experiments in duplicate. C. Dose-response curve of *IL6* expression in TMF- or dTMF-treated astrocytes activated with IL1-β (n = 3 per dose normalized to n = 6 DMSO, mean ± SD), representative of 3 independent experiments. D. Normalized expression (IL6/GAPDH) relative to DMSO from a ddPCR experiment showing no effect of dCHR on *IL6* expression in IL1-β-activated astrocytes (n = 1 biological samples for DMSO and dCHR + IL1-β, n = 2 for DMSO + IL1-β, Poisson error of >15,000 droplets). E. Flavones do not reduce cell viability of primary human astrocytes at 20 µM. Each dot represents percentage of non-Zombie Violet-stained cells in 1 biological sample from 2-5 independent experiments (n = 2-10, mean ± SD). * P < 0.05; ** P < 0.01; *** P < 0.001; **** P < 0.0001

**Extended Data Figure 2.**
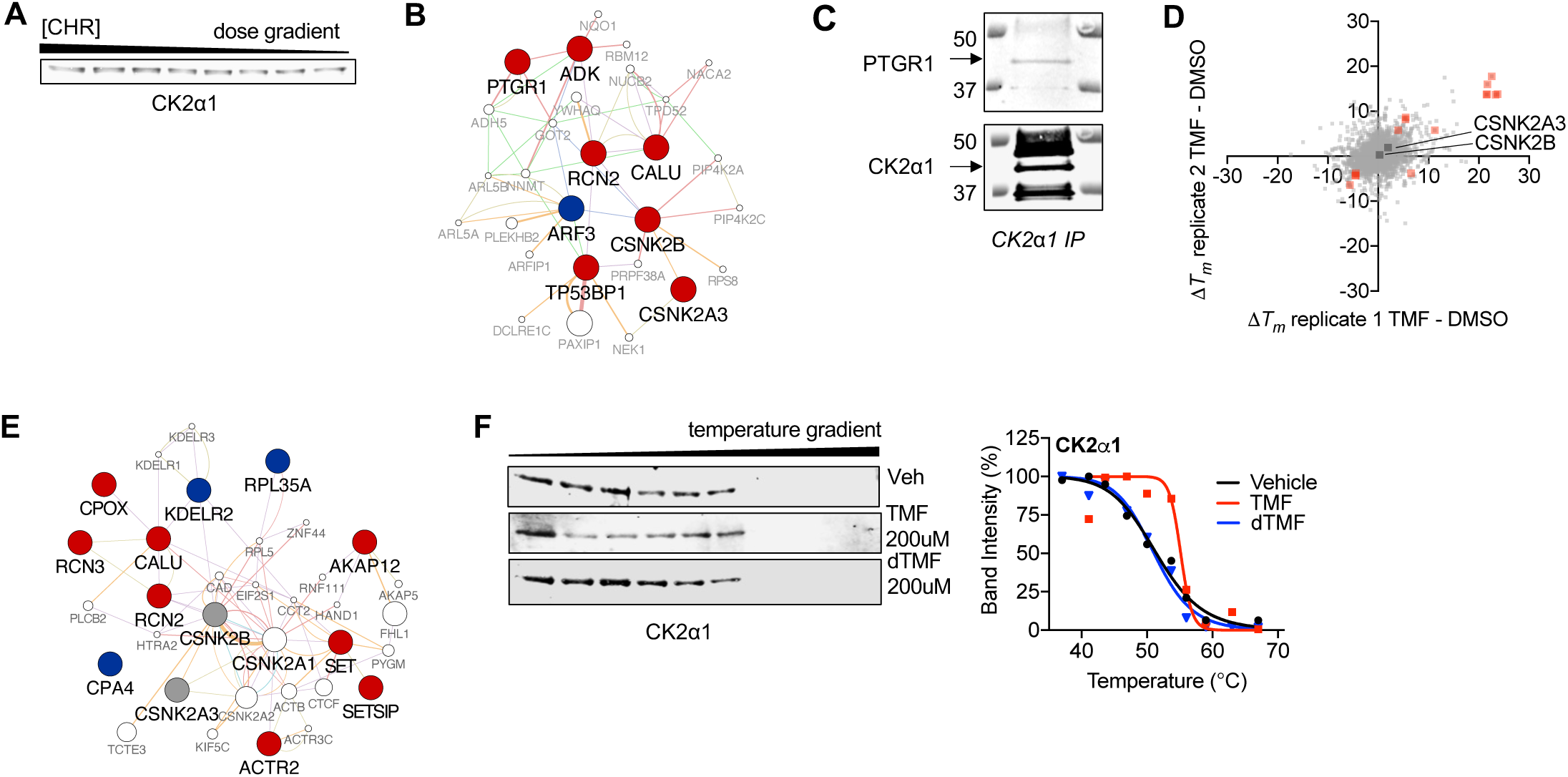
Thermal shift proteome profiling and Western blotting show stabilization of CK2 and CK2 interactors by flavones. A. TSA-WB isothermal dose response showing stabilization of CK2α1 by CHR (see Figure 1F). Full blots in Extended Data Figure 15. B. GeneMania network showing protein-protein interactions between hits from Figure 1E (red nodes: stabilized, blue nodes: destabilized) and related proteins (white nodes). C. Immunoblot showing physical interaction of PTGR1 and CK2α1 by co-immunoprecipitation. Full blots in Extended Data Figure 15. D. TPP in human iPSC-derived astrocytes. Red denotes proteins exhibiting significant and reproducible thermal shifts after TMF treatment in IL1-β-activated astrocytes. E. GeneMania network showing protein-protein interactions between hits from Extended Data Fig. 1C (red nodes: stabilized, blue nodes: destabilized, gray nodes: not significant) and related proteins (white nodes). F. TSA-WB (left) with TMF and dTMF, along with band intensity quantification (right). Full blots in Extended Data Figure 15.

**Extended Data Figure 3.**
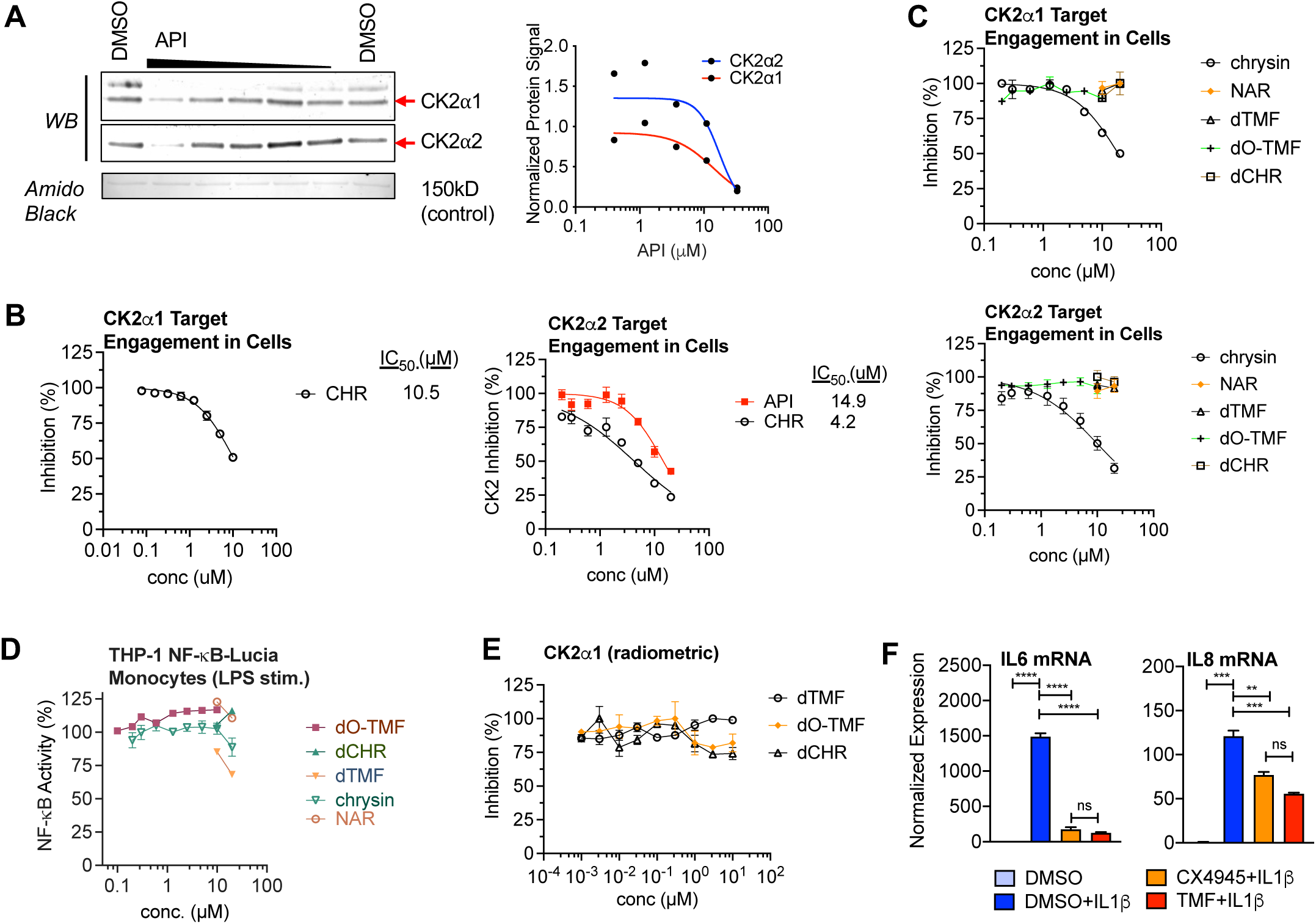
Additional characterization of flavones for *in vitro* and in-cell CK2 inhibition. A. Kinase capture shows apigenin is an ATP-competitive inhibitor of CK2α1 and CK2α2 in H1-derived astrocyte whole lysates. Graph shows band intensity normalizations to a 150 kD protein stained by Amido Black; lines represent nonlinear curve fits of dose-curve data. Representative of 2 independent experiments. Full blots in Extended Data Figure 15. B. NanoBRET assay shows dose-dependent CK2α1 and CK2α2 target engagement in HEK293T cells (each point mean ± sem, n = 3). Nonlinear fit was used to calculate IC_50_. C. NanoBRET assay shows dose-dependent CK2α1 and CK2α2 target engagement of chrysin in HEK293T cells, but not NAR, dCHR, dTMF, or dO-TMF (each point mean ± sem, n = 3). Nonlinear fit was used to calculate IC_50_. D. No activity of chrysin, NAR, dCHR, dTMF, or dO-TMF in THP-1-NF-κB-Lucia monocytes (each point mean ± sem, n = 3). Nonlinear fit was used to calculate IC_50_. E. Radiometric kinase assays using recombinant CK2α1 do not show inhibitory activity of NAR, dCHR, dTMF, or dO-TMF. F. Structurally unrelated CK2 inhibitor CX-4945 blocks IL1β-induced IL6 upregulation as well as TMF (mean ± SD, n = 2, one-way ANOVA with Tukey’s post-hoc test). Normalized to DMSO, data are representative of 3 independent experiments. ** P < 0.01; *** P <0.001; **** P < 0.0001

**Extended Data Figure 4.**
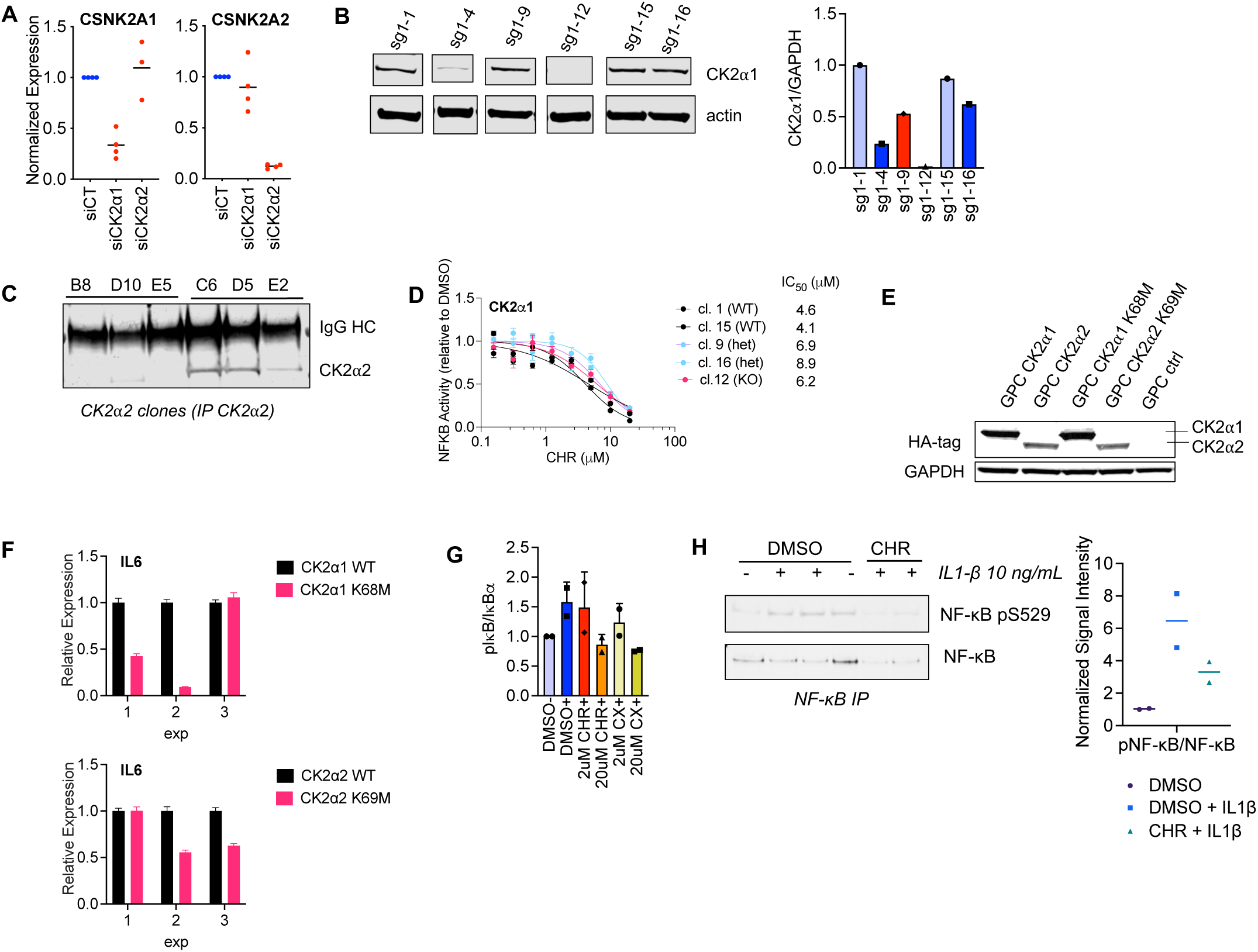
CK2 genetic perturbation assays and NF-κB immunoprecipitation. A. Knockdown efficiency of experiments shown in Figure 2B (mean expression shown with each dot representing an experiment). B. Western blotting showing expression of CK2Α1 and GAPDH in individual THP-1 NF-κB-Lucia clones, quantified as a relative ratio. Heterozygotes and homozygotes confirmed by Sanger sequencing. Full blots in Extended Data Figure 16. C. Western blotting showing CK2Α2 expression in individual WT or KO THP-1 NF-κB-Lucia clones after immunoprecipitation of CK2Α2. Full blots in Extended Data Figure 16. D. Quantification of NF-κB reporter luciferase activity in CK2Α1 heterozygous (het) or homozygous (KO) knockout THP-1 NF-κB Lucia cell clones stimulated with 20 ng/mL LPS and CHR dilution series (mean ± sem, n = 3). E. Expression of CK2α1-HA and CK2α2-HA in GPCs confirmed by Western blotting. Full blots in Extended Data Figure 15. F. CK2 kinase-dead mutants block IL1β-induced IL6 upregulation in HCA (3 independent experiments shown with Poisson error; relative expression normalized to each WT). G. Quantification showing that pIκB/IκBα levels are reduced with CK2 inhibitors (6-hour treatment, mean ± SD, n = 2, pooled independent experiments; related to Figure 3C). H. Immunoblot and quantification (n = 2, mean) of NF-κB IP showing NF-κB S529 phosphorylation is reduced with CHR (2-hour treatment). Full blots in Extended Data Figure 16.

**Extended Data Figure 5.**
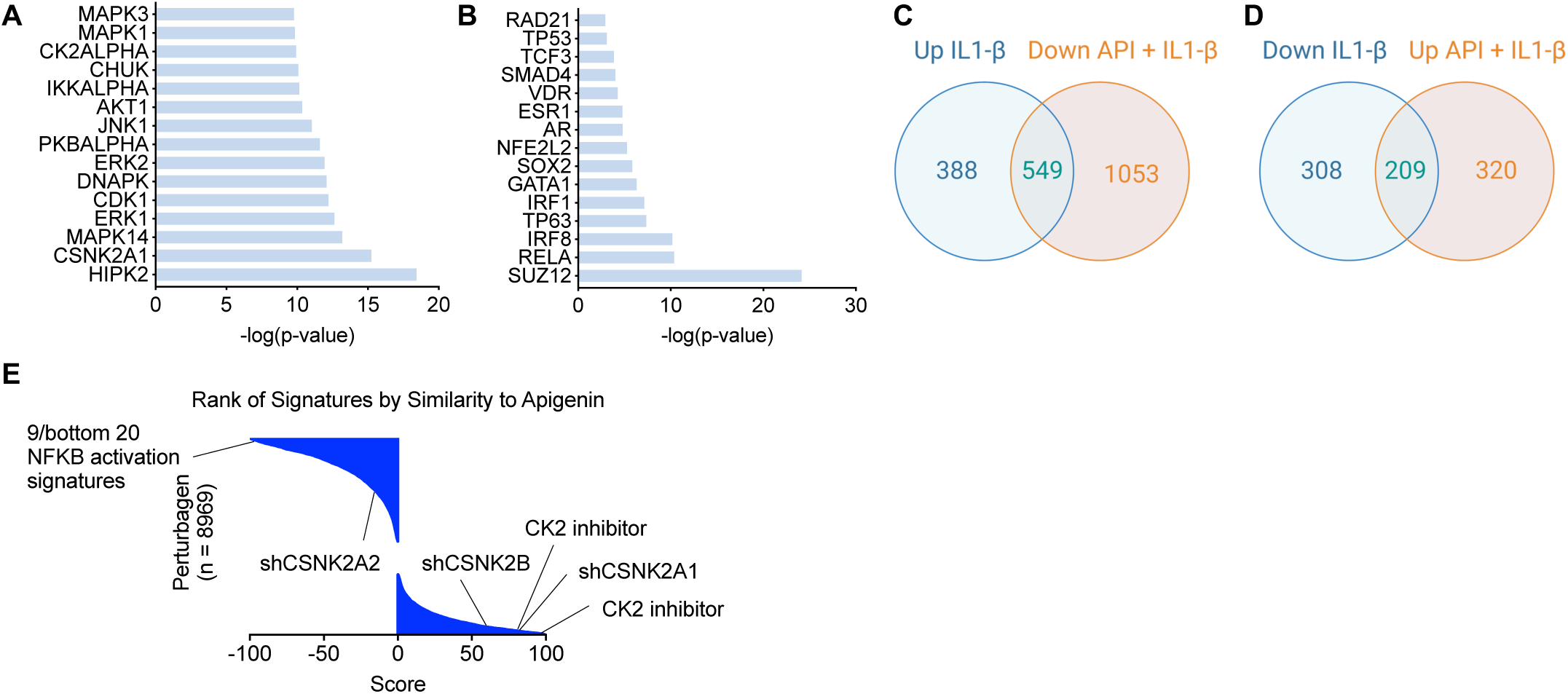
RNA-seq in IL1-β- and API-treated astrocytes. A. Bar graph depicting p-values associated with enriched kinases in IL1-β-treated astrocytes compared to controls (KEA). B. Bar graph depicting p-values associated with enrichment of TFs and other chromatin factors from transcription factor enrichment analysis (TFEA). C. Venn diagram showing overlap of significantly differentially expressed (DE) genes up in IL1-β and down in API + IL1-β-treated cells. D. Venn diagram showing overlap of significantly DE genes down in IL1-β and up in API + IL1-β-treated cells. E. Rainfall plot showing rank of CMap signatures by z-score in terms of similarity to API + IL1-β-treated HCAs (0 = no correlation; 100 = most positive correlation; -100 = most negative correlation). Of the 20 signatures with the most negative scores, 9 are NFKB activation signatures.

**Extended Data Figure 6.**
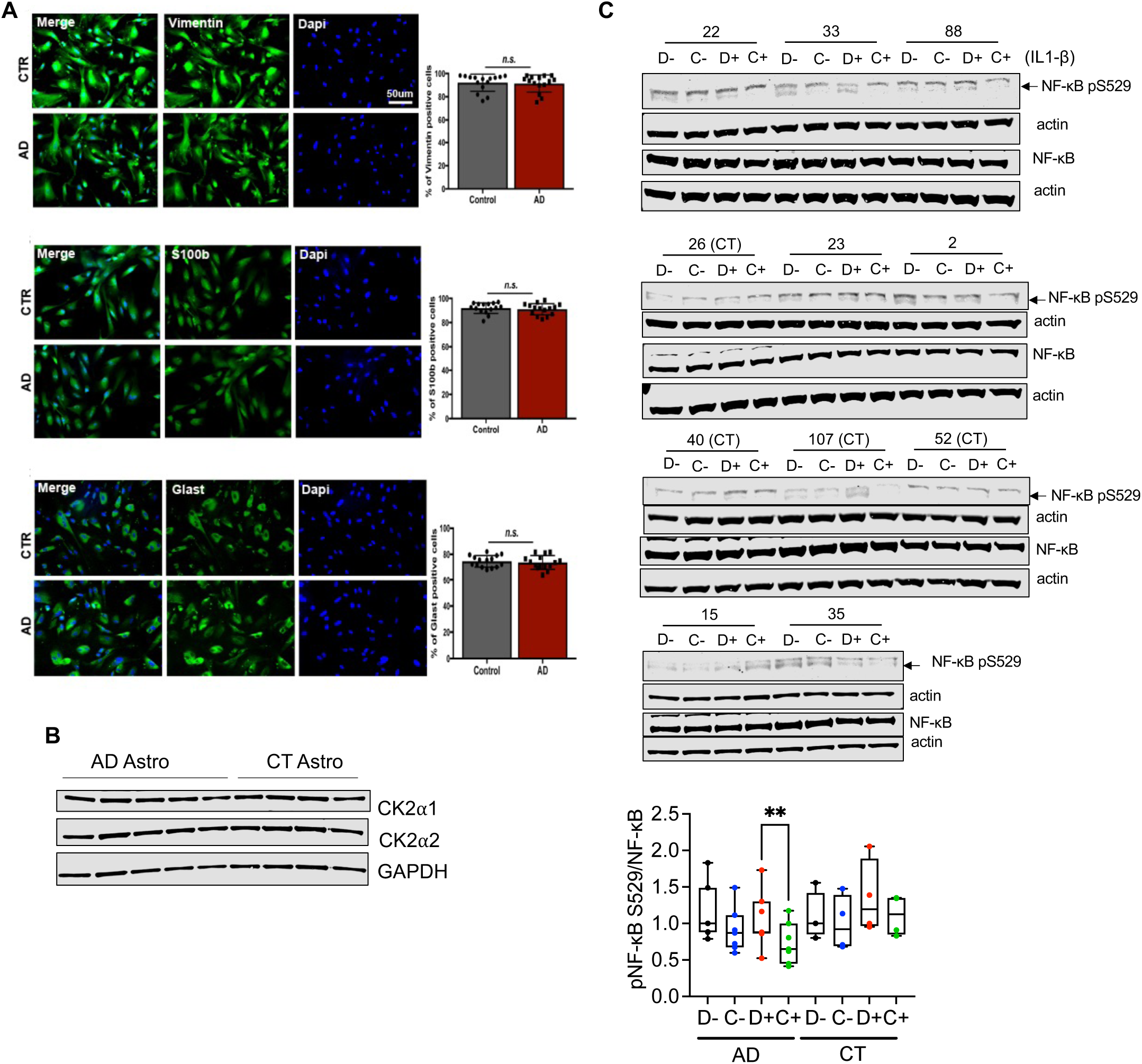
Characterization of CK2 levels in dementia or AD postmortem samples and astrocyte marker staining and pNF-κB/NF-κB levels in AD and CT patient-derived astrocytes. A. Representative images of immunostainings of astrocytes expressing astrocyte markers: Vimentin, S100B and GLAST. Quantification of the fraction of astrocytes expressing astrocyte markers reveals a homogeneous population of iPSC-derived astrocytes from both AD and controls. Data represent mean ± SEM for n=5 randomized images counted blindly for each line (total of 6 lines: 3 AD and 3 controls). B. Representative immunoblots associated with Figure 4D. Full blots in Extended Data Figure 16. C. Blots and quantification of pNF-κB/NF-κB levels (median and 25-75% CI of n = 7 AD and n = 4 CT samples; mixed-effects ANOVA with Sidak’s post-hoc test), associated with Figure 4F. Cell lines 40, 107, 52, and 26 are controls, whereas the rest of the cell lines are AD. D- = DMSO; D+ = DMSO + IL1β; C- = CHR, C+ = CHR + IL1β. Full blots in Extended Data Figure 17. Ns = not significant; * P < 0.05; ** P < 0.01

**Extended Data Figure 7.**
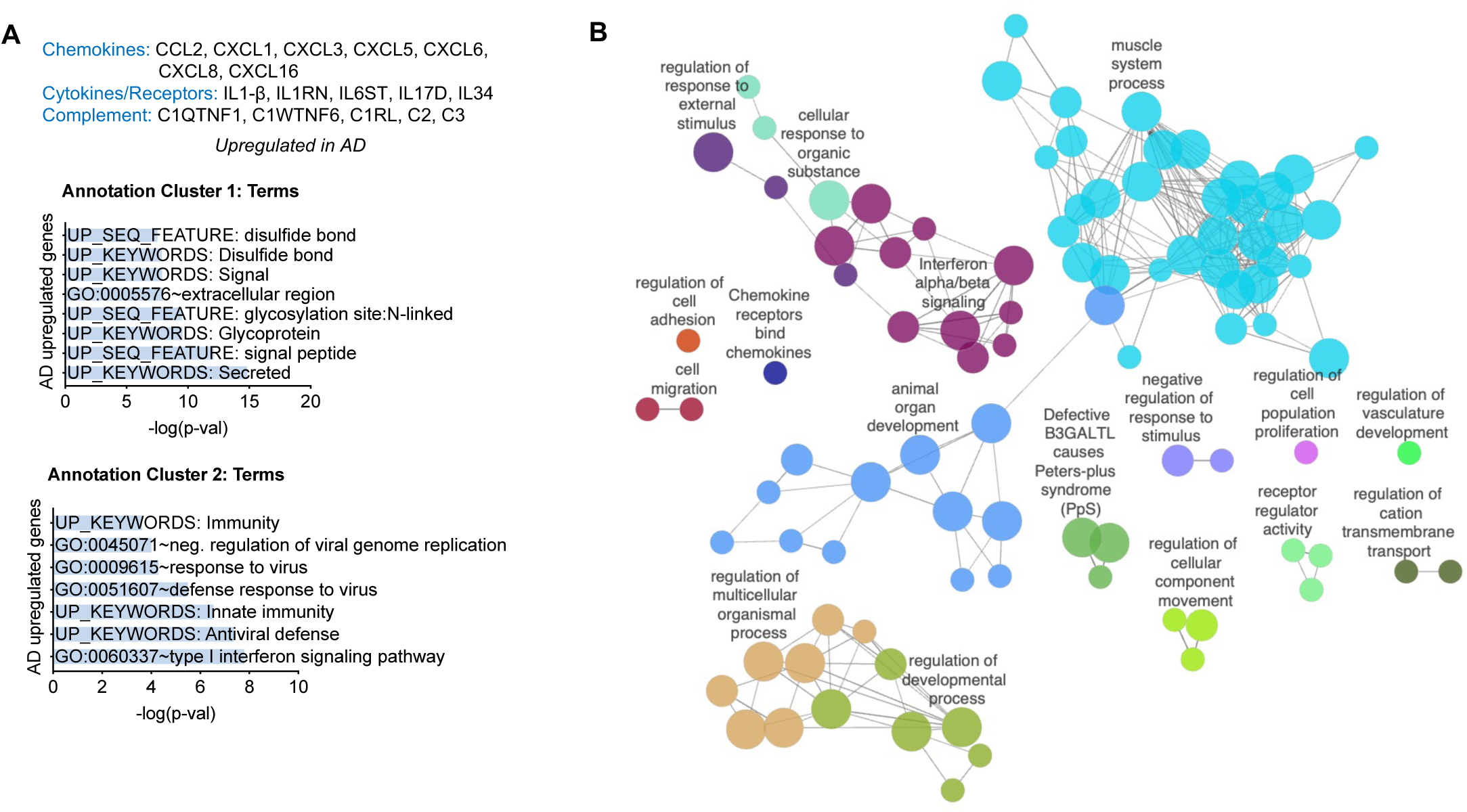
GO and network analysis of DE genes in AD vs. CT patient-derived astrocytes. A. Graphs depicting the top 2 annotation clusters and their associated terms and -log(p-values) from DAVID gene ontology analysis of significantly AD-upregulated genes. B. Network plot of ClueGO analysis using significantly upregulated genes in AD astrocytes versus CT (Cytoscape).

**Extended Data Figure 8.**
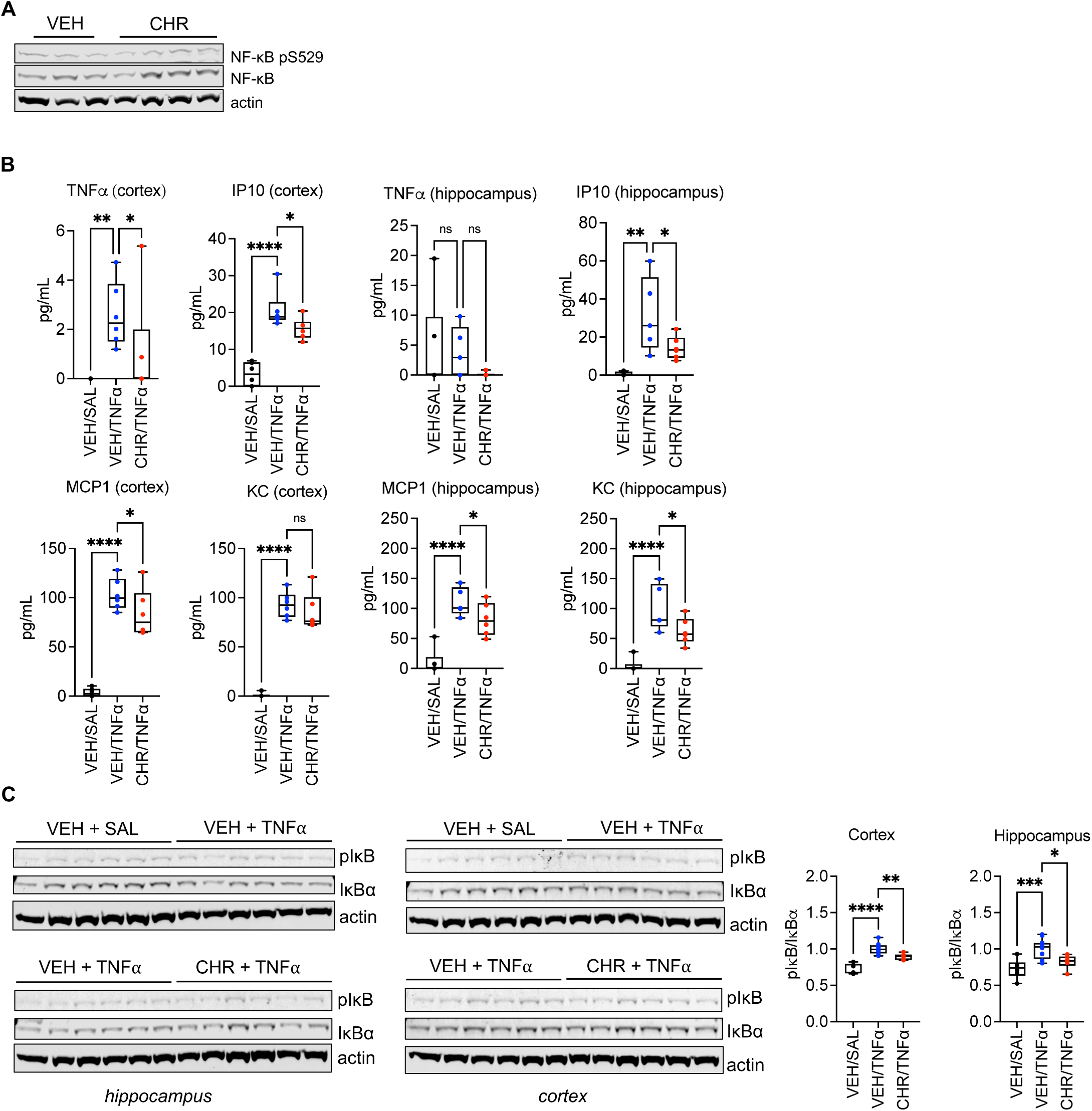
Efficacy of CHR in murine TNFα neuroinflammation model. A. Immunoblots associated with Figure 4I. Full blots in Extended Data Figure 18. B. Box plots showing median and 25-75% CI of MCP-1, IP10, KC, and TNFα cytokine expression in the cortex and hippocampus (n = 6 mice per group, one-way ANOVA with adjusted p-values via Benjamini, Krieger, Yekutieli FDR method). C. Immunoblots and quantification of pIκB S32/IκBα signal in cortical and hippocampal tissues from experiment described in (B) above (mean of n = 6 animals per group except for one VEH/SAL sample for cortex omitted in quantification due to artifact in lane, with points from VEH/TNFα groups plotted in duplicate (n = 12); one-way ANOVA with Dunnett’s post-hoc test). Full blots in Extended Data Figure 18. ns = not significant; * P < 0.05; ** P < 0.01; *** P < 0.001; **** P < 0.0001

**Extended Data Figure 9.**
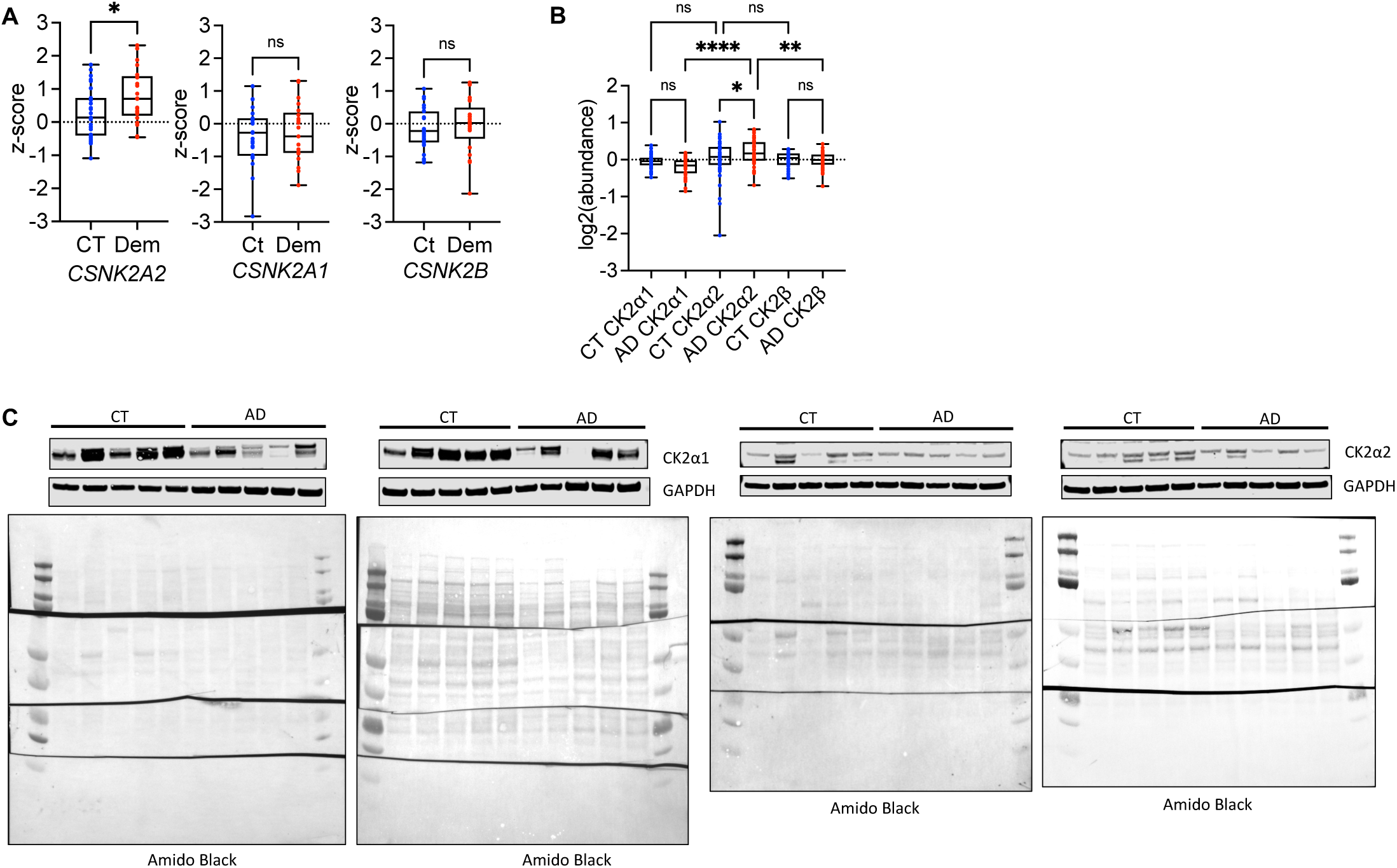
Analysis of CK2 isoform expression in published postmortem datasets and fresh frozen NIH Neurobiobank samples. A. Expression of *CSNK2A2, CSNK2A1* or *CSNK2B* in postmortem brains (parietal cortex) from patients with dementia (Dem, n = 21) or controls (CT, n = 27) from the Aging, Dementia, and TBI RNA-seq study^62^. Box plots depicting mean z-score (two-tailed t-test). B. CK2α2 protein levels in AD are higher than controls (CK2α1 and CK2β not significant) and relative levels of CK2α2 are higher than CK2α1 and CK2β in AD patients, but not in controls. Data from the UPP Proteomics Study^63^. Box plot depicting mean log2(abundance) (n = 47 for CT CK2α1 and CT CK2β, n = 44 for CT CK2α2, n = 49 for AD CK2α1 and AD CK2β, n = 48 for AD CK2α2, one-way ANOVA with Sidak’s post-hoc test). C. Representative immunoblots and Amido Black stains of postmortem cortical brain samples from CT and AD patients related to Figure 5A. * P < 0.05; ** P < 0.01; *** P < 0.001; **** P < 0.0001

**Extended Data Figure 10.**
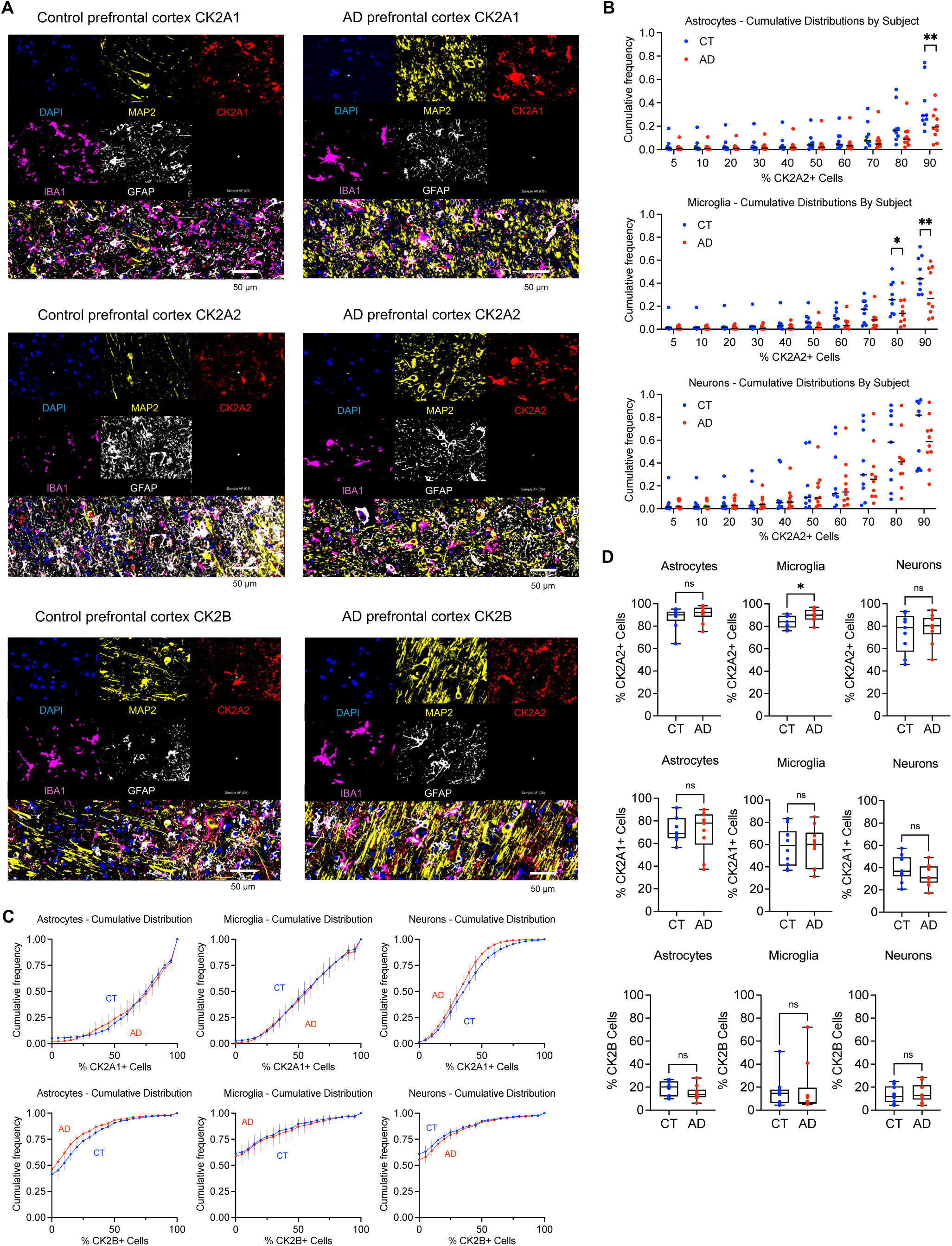
CK2 isoform staining in AD postmortem frontal cortex. A. Representative immunofluorescence images showing expression of CK2 isoforms in neurons (MAP2), astrocytes (GFAP), and microglia (IBA1), counterstained with DAPI, from the prefrontal cortex of an age-matched cognitively normal individual (left) or an AD patient (right). B. Cumulative frequencies per subject at CK2A2 positivity levels representative of the whole distribution extracted from data in Figure 5B (n = 9 subjects CT, n = 10 subjects AD; two-tailed t-tests with p-values adjusted for multiple comparisons by two-stage linear step-up method by Benjamini, Krieger and Yekuteili). C. Cumulative distribution plots of the area covered by each respective CK2A1+ or CK2B+ cell population across image tiles in control or AD groups (mean ± sem shown, n = 10 subjects per group). D. Box plots showing mean percentage of area covered by CK2A1+ cells by cell type in postmortem cortical brain AD or control samples from the UCSD ADRC as measured by immunofluorescence staining (median and 25-75% CI, n = 10 individuals per group, two-tailed t-test) or similar analyses for CK2B (median and 25-75% CI, n = 10 individuals per group, two-tailed t-test). * P < 0.05, ** P < 0.01

**Extended Data Figure 11.**
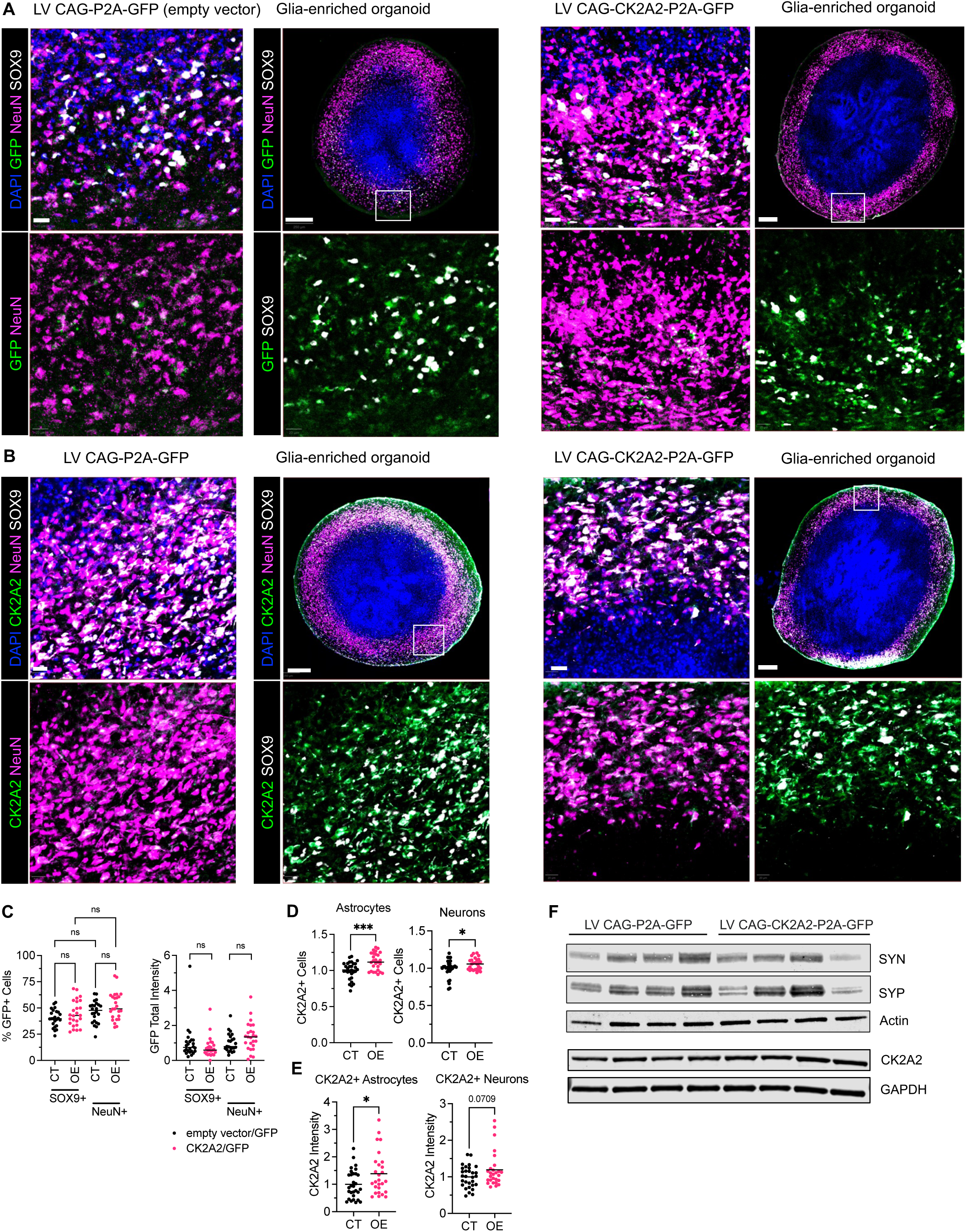
Additional characterization of CK2α2 overexpression in glia-enriched organoids. For Figures A-H, CT = empty vector and OE = CK2α2 overexpression. A. Representative immunofluorescence images showing GFP expression of astrocytes (SOX9) and neurons (NeuN), counterstained with DAPI, from glia-enriched organoids transduced with LV CAG-P2A-GFP (empty vector) or LV CAG-CK2A2-P2A-GFP. See Extended Data Figure 11A for quantification. Scale bar = 250 µm and 20 µm (inset). B. Representative immunofluorescence images showing CK2A2 expression of astrocytes (SOX9) and neurons (NeuN), counterstained with DAPI, from glia-enriched organoids transduced with LV CAG-P2A-GFP or LV CAG-CK2A2-P2A-GFP. Scale bar = 250 µm and 20 µm (insets). C. Quantification of percentage of GFP+ cells (one-way ANOVA with Sidak’s multiple comparisons test) and normalized proportion GFP total intensity (two tailed t-tests) by cell type (SOX9+ = astrocytes; NeuN+ = neurons); related to Figure 5C. Median of pooled data from 2 independent experiments shown (n = 24 CT and n = 25 OE organoids; two-tailed t-test). D-E. Quantification of normalized proportion of CK2A2+ astrocytes or neurons (D) and CK2A2 intensity in CK2A2+ astrocytes or neurons (E) (n = 30 CT and n = 27 OE organoids; two-tailed t-test). F. Representative immunoblots related to Figure 5F. Full blots in Extended Data Figure 19. * P < 0.05; ** P < 0.01; *** P < 0.001

**Extended Data Figure 12.**
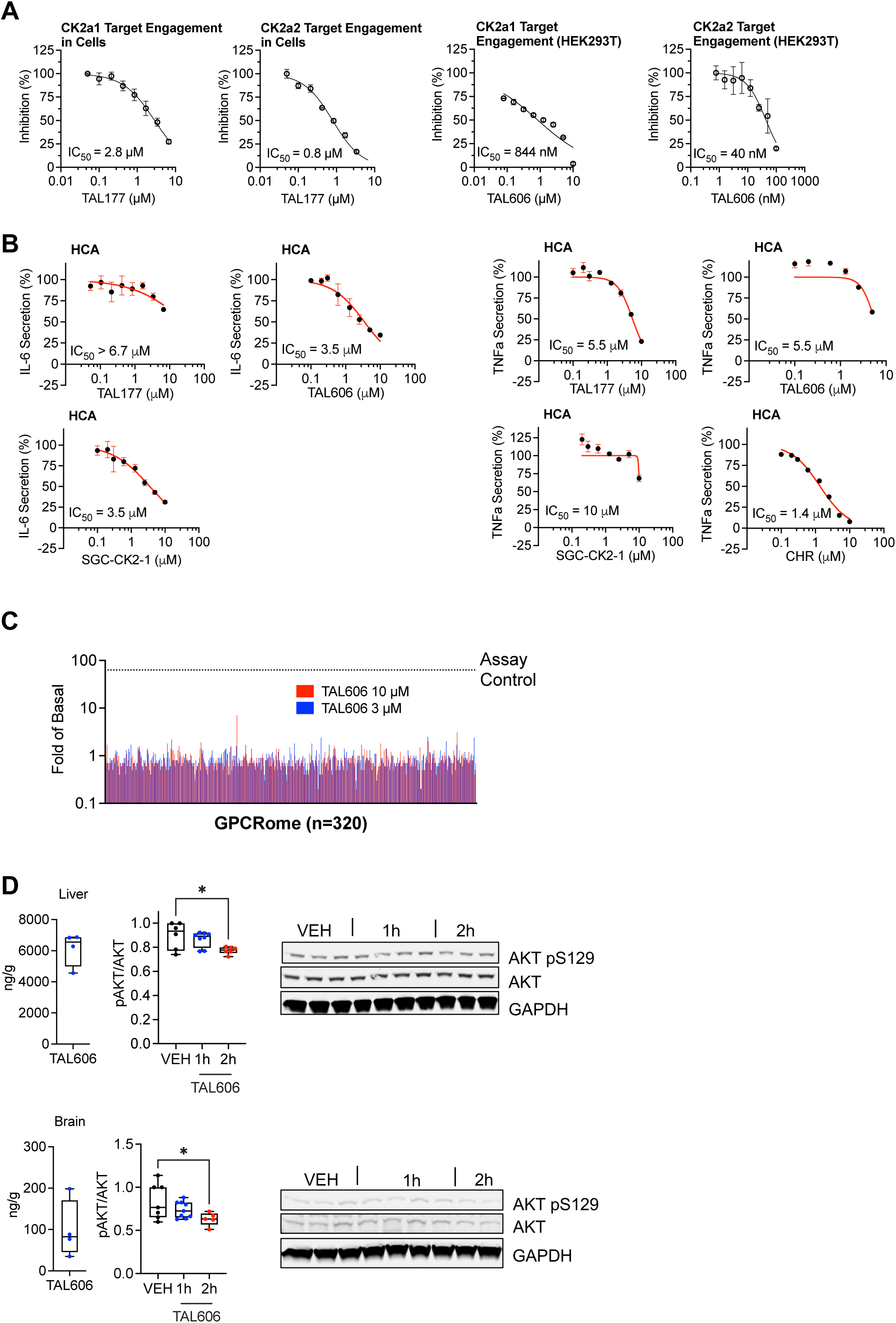
Cellular activity, selectivity, and PK/PD characterization assays of synthetic CK2 inhibitors. A. NanoBRET assay shows dose-dependent CK2α1 and CK2α2 target engagement of TAL177 and TAL606 in HEK293T cells (each point mean ± sem, n = 3). Nonlinear fit was used to calculate IC50. B. Dose-dependent anti-inflammatory activity of CHR, TAL177, TAL606, and SGC-CK2-1 in human astrocytes via block of IL1-β-induced IL-6 or TNFa secretion, respectively (n = 3, mean ± s.e.m.). Normalized to DMSO vehicle ± stimulation; representative of 2 independent experiments. C. Bar plots of TAL606 tested at 10 µM and 3 µM in the NIH PDSP GPCRome panel showing average fold-change of activity for each GPCR (n = 320 GPCRs). Dotted line represents the average fold-change in activity of D2 receptors by quinpirole 100 nM (Assay Control). Values ≥ 3.0 represent potential agonist activity. D. Concentrations of TAL606 in liver and brain at Cmax (1h; median and 25-75% CI, n = 4 mice) and immunoblots and quantification of AKT pS129 relative to total AKT in brain or liver lysates of mice treated with one dose of TAL606 at 62.5 mpk IP for specified times (mean and 25-75% CI, n = 3-4 mice in duplicate, one-way ANOVA). * P < 0.05

**Extended Data Figure 13.**
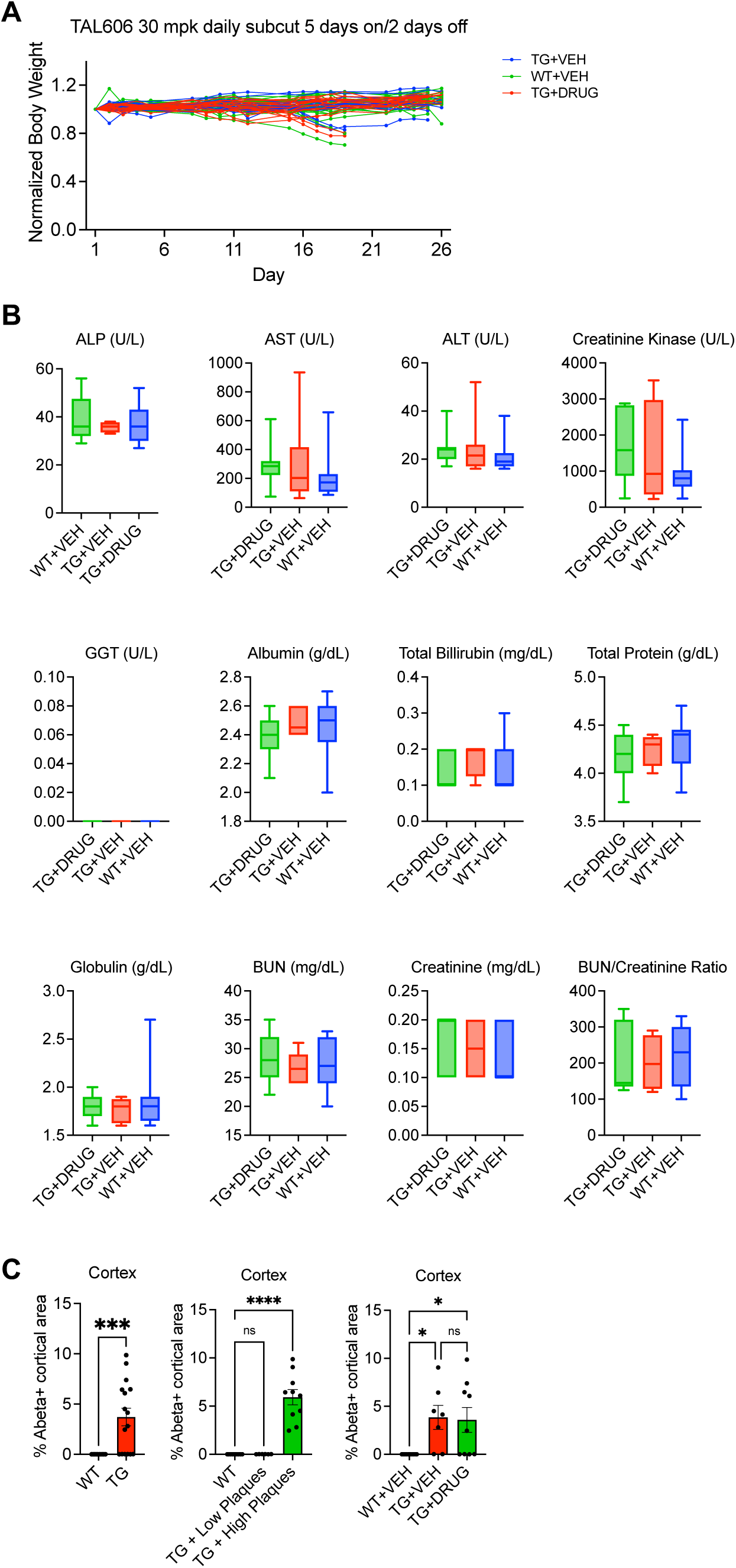
Tolerability studies of TAL606 in APP/PS1 mice. A. Individual body weights over time as fraction of starting weight for mice treated with 30 mpk TAL606 5 days on/2 days off for 26 days (n = 26 mice WT+VEH, n = 18 TG+VEH, n = 19 TG+TAL606). B. Box plots showing serum clinical chemistry measurements of 12 analytes indicative of kidney and liver health in the IDEXX Standard Tox Mouse panel (median and 25-75% CI, n = 13 WT+VEH, n = 8 TG+VEH, n = 7 TG+TAL606). C. Bar graphs showing percent cortical area covered by plaques, with each dot representing the average over 3-4 images per brain (mean ± sem, n = 10 WT, n = 16 TG mice, of which n = 7 TG + VEH and n = 9 TG +TAL606). Representative of 2 independent experiments. * P < 0.05; ** P < 0.01; *** P < 0.001; **** P < 0.0001

**Extended Data Figure 14.**
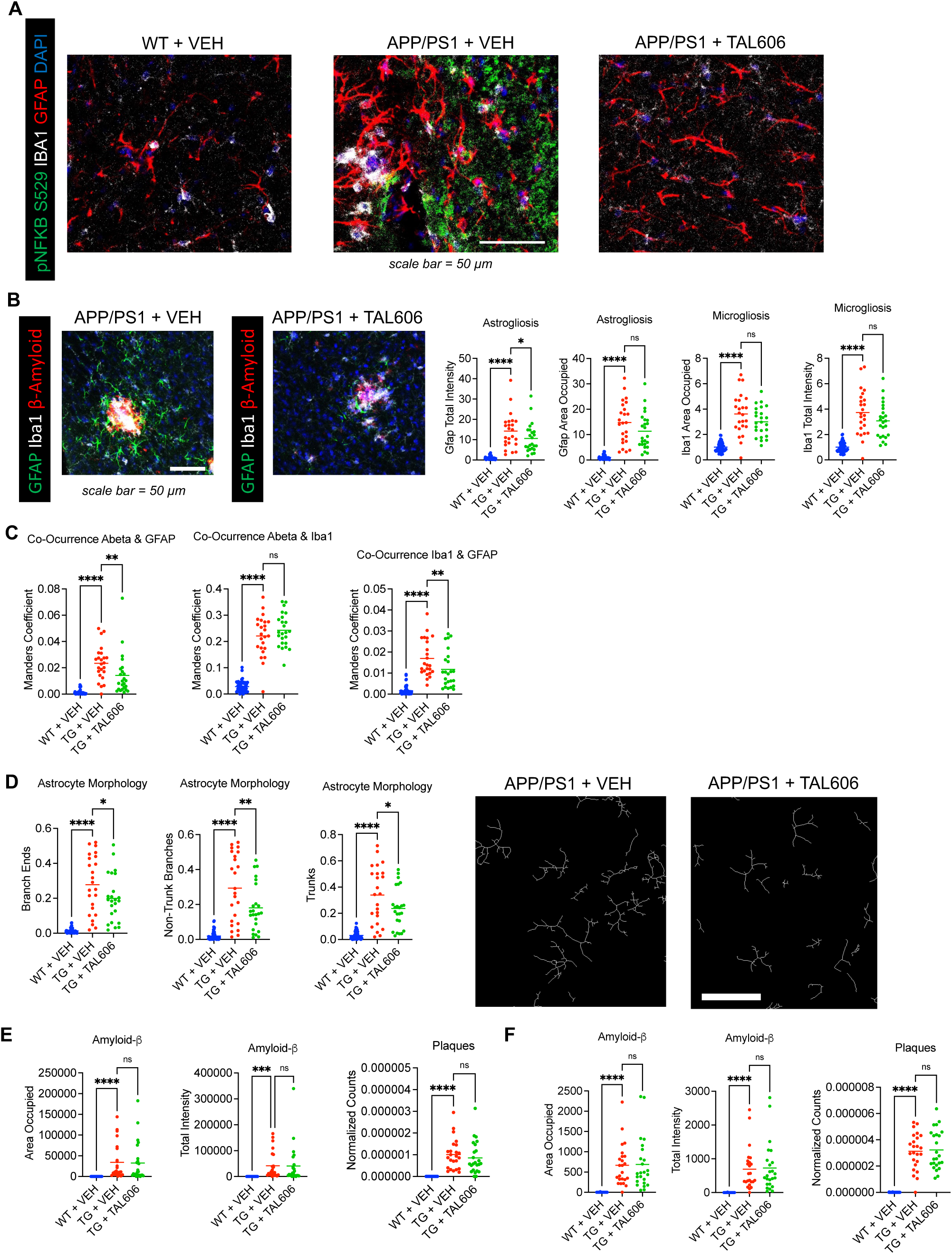
Additional characterization of TAL606 activity in APP/PS 1 mice. A. Representative immunofluorescence images showing pNF-κB expression in astrocytes (GFAP+) or microglia (Iba1+) within hippocampal CA1 regions; related to Figure 6H. B. (Left) Representative immunofluorescence images and (Right) quantification of astrogliosis (GFAP area occupied and total intensity) and microgliosis (Iba1 area occupied and total intensity) in the cortices of WT + VEH (N=18), TG + VEH (N=8), and TG + TAL606 (N=8) mice pooled from 2 independent cohorts, imaged in triplicate, and normalized to image stack volumes. C. Co-occurrence of amyloid-beta with astrocytes (GFAP) or microglia (Iba1) or astrocytes and microglia with each other were analyzed within cortex ROIs by calculating Manders coefficients in CellProfiler within maximum projection images of each image tile. Each point = 1 quantified image tile (850 µm x 850 µm area) from N=8 mice per group TG + VEH and TG + TAL606, N=18 mice WT + VEH, pooled from 2 independent cohorts. For each mouse, 3 total images of the cortex were quantified. D. Astrocyte morphology was analyzed within cortex ROIs by CellProfiler using MeasureObjectSkeleton in GFAP binary images (filtered for min and max sizes to remove artifacts). Representative skeletons shown. GFAP binary images were created from the maximum projection images of each image tile within ROIs. Each point = 1 quantified image tile from N=8 mice per group TG + VEH and TG + TAL606, N=18 mice WT + VEH, pooled from 2 independent cohorts. For each mouse, 3 total images of cortex were quantified. Related to Figure 6H. Scale bar = 50 µm. E. Area occupied, total intensity, and plaque counts were analyzed within hippocampus ROIs by CellProfiler after thresholding the Abeta channel across maximum projection images of each image tile. Each point = 1 quantified image tile (850 µm x 850 µm area) from N=8 mice per group TG + VEH and TG + TAL606, N=18 mice WT + VEH, pooled from 2 independent cohorts. For each mouse, 3 total images of the CA1, CA3, and DG regions of the hippocampus were quantified. Raw values were normalized to image stack volumes. F. Area occupied, total intensity, and plaque counts were analyzed within cortex ROIs by CellProfiler after thresholding the Abeta channel across maximum projection images of each image tile. Each point = 1 quantified image tile (850 µm x 850 µm area) from N=8 mice per group TG + VEH and TG + TAL606, N=18 mice WT + VEH, pooled from 2 independent cohorts. For each mouse, 3 total images of the cortex were quantified. Raw values were normalized to image stack volumes. For panels B and E-F, area or intensity data in each cohort were normalized to their WT group average. * P < 0.05; ** P < 0.01; *** P < 0.001; **** P < 0.0001

**Extended Data Figure 15.**
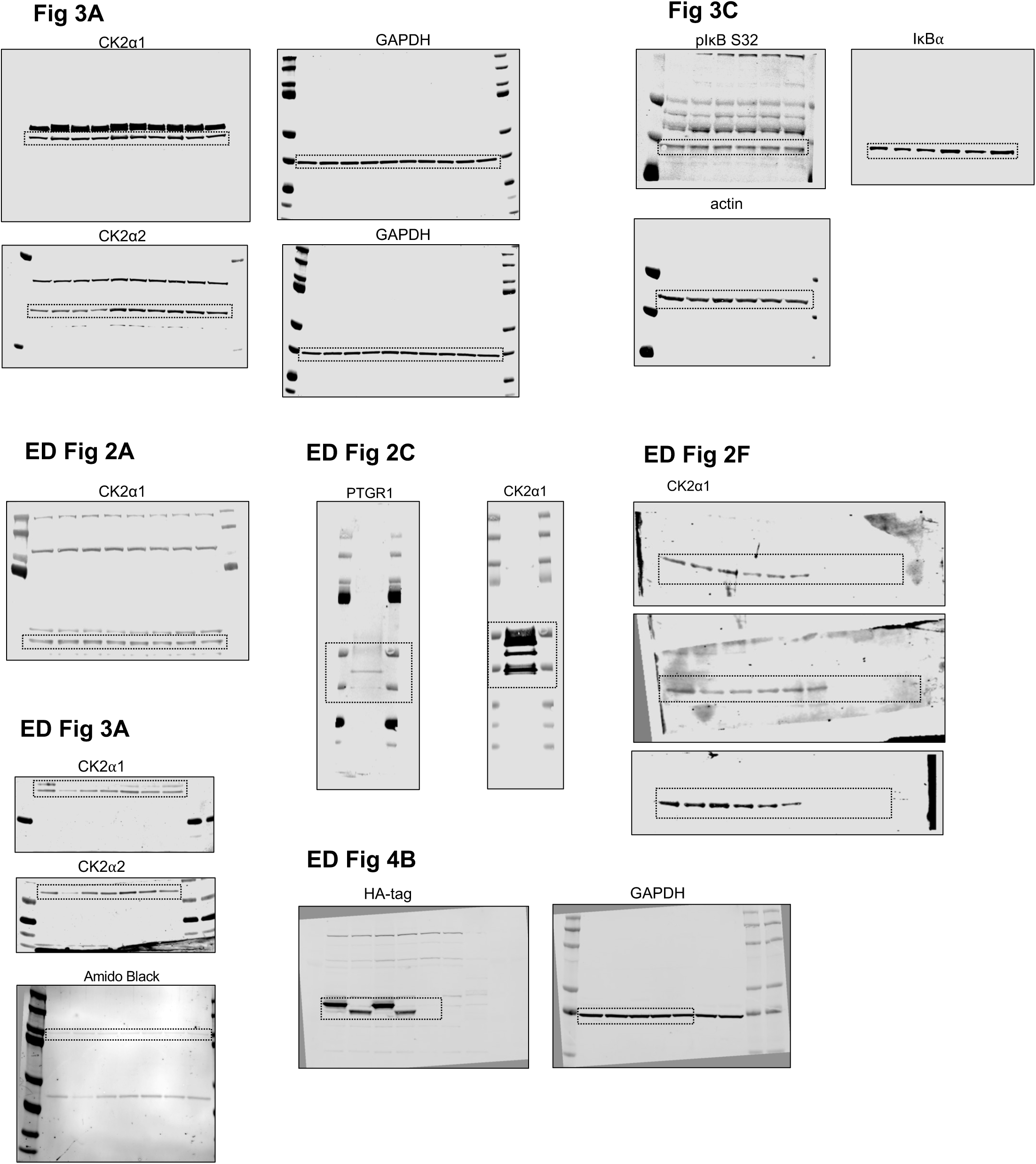
Original full blot images from Odyssey scans (Part I).

**Extended Data Figure 16.**
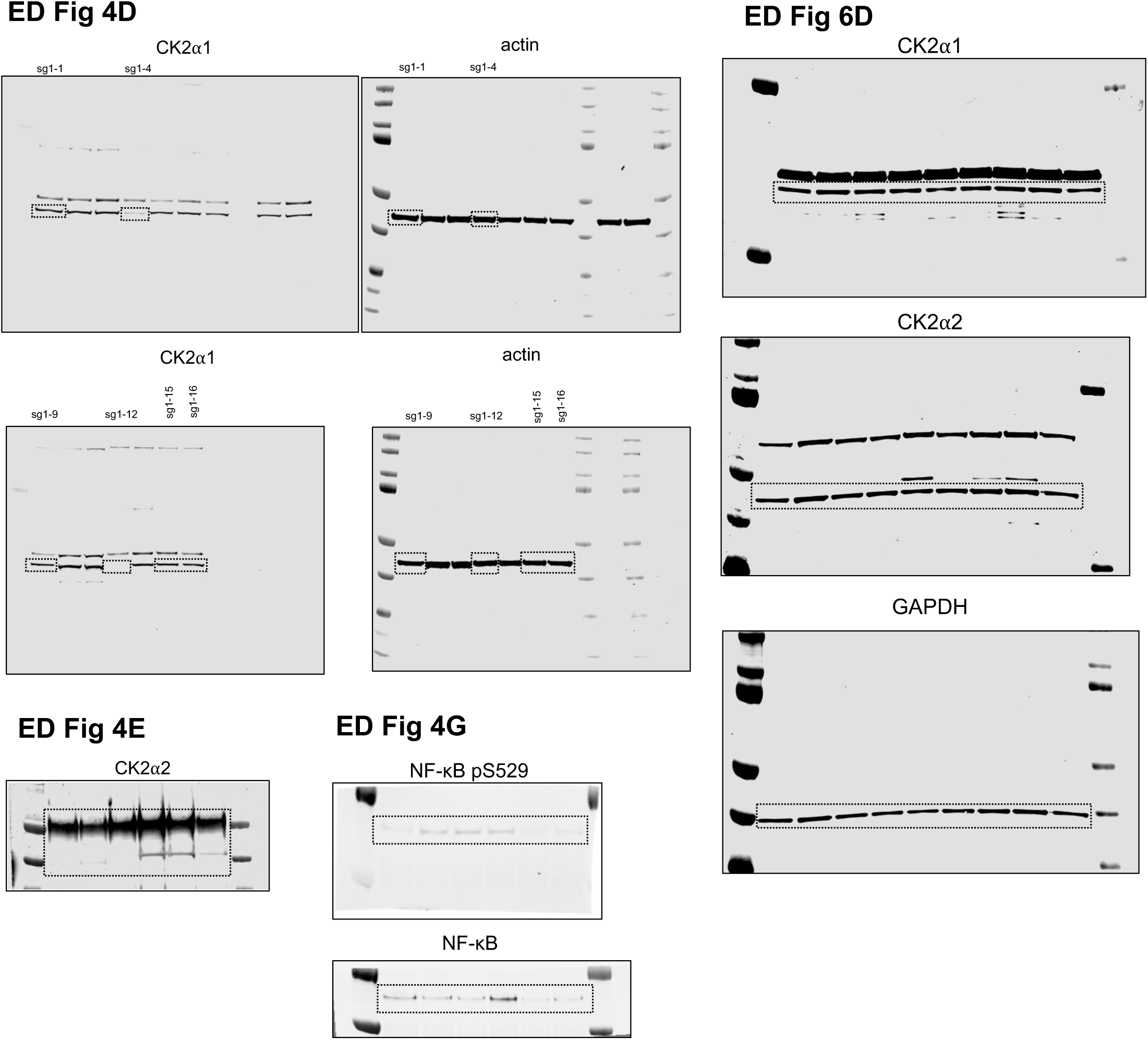
Original full blot images from Odyssey scans (part II).

**Extended Data Figure 17.**
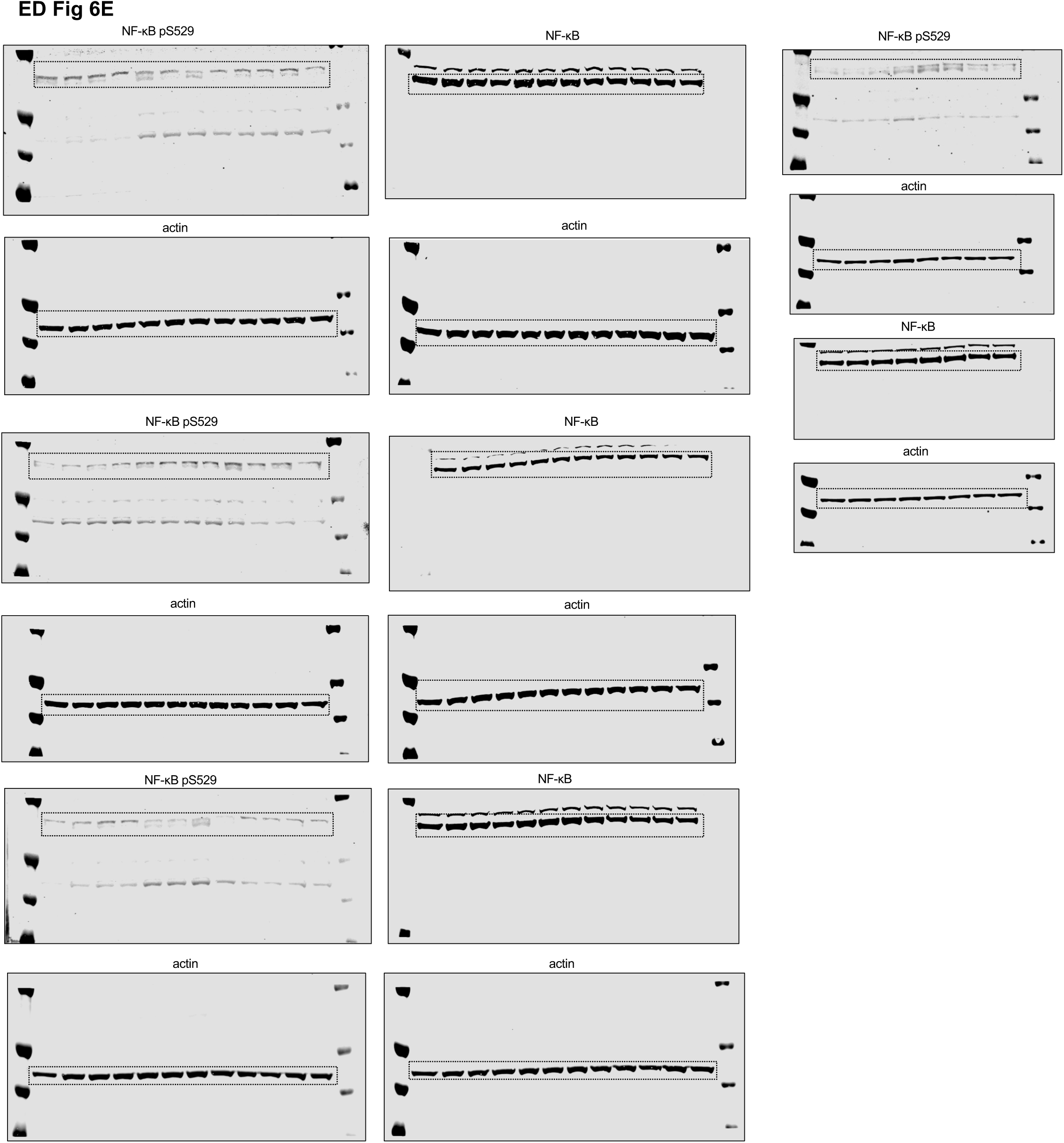
Original full blot images from Odyssey scans (part III).

**Extended Data Figure 18.**
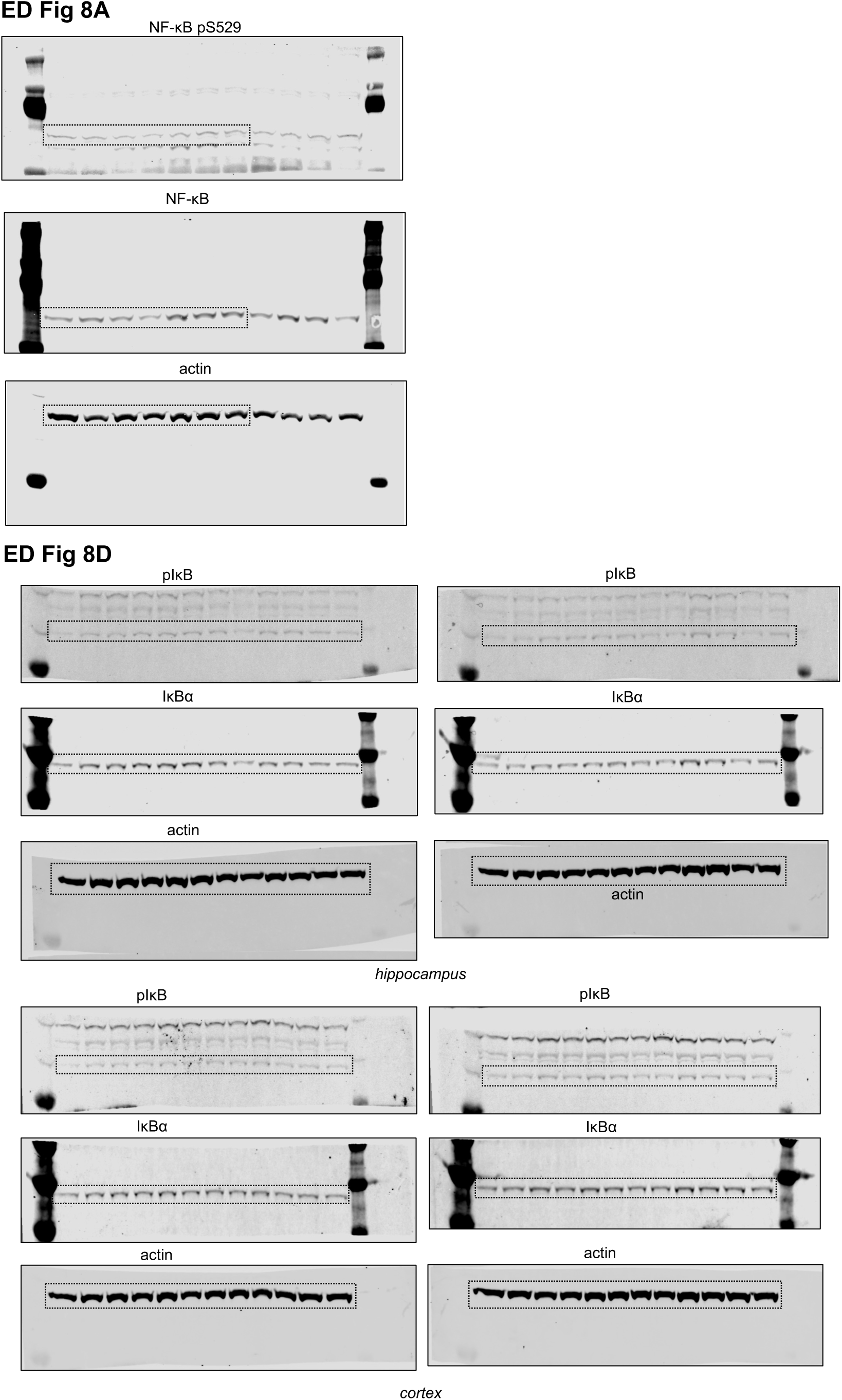
Original full blot images from Odyssey scans (part IV).

**Extended Data Figure 19.**
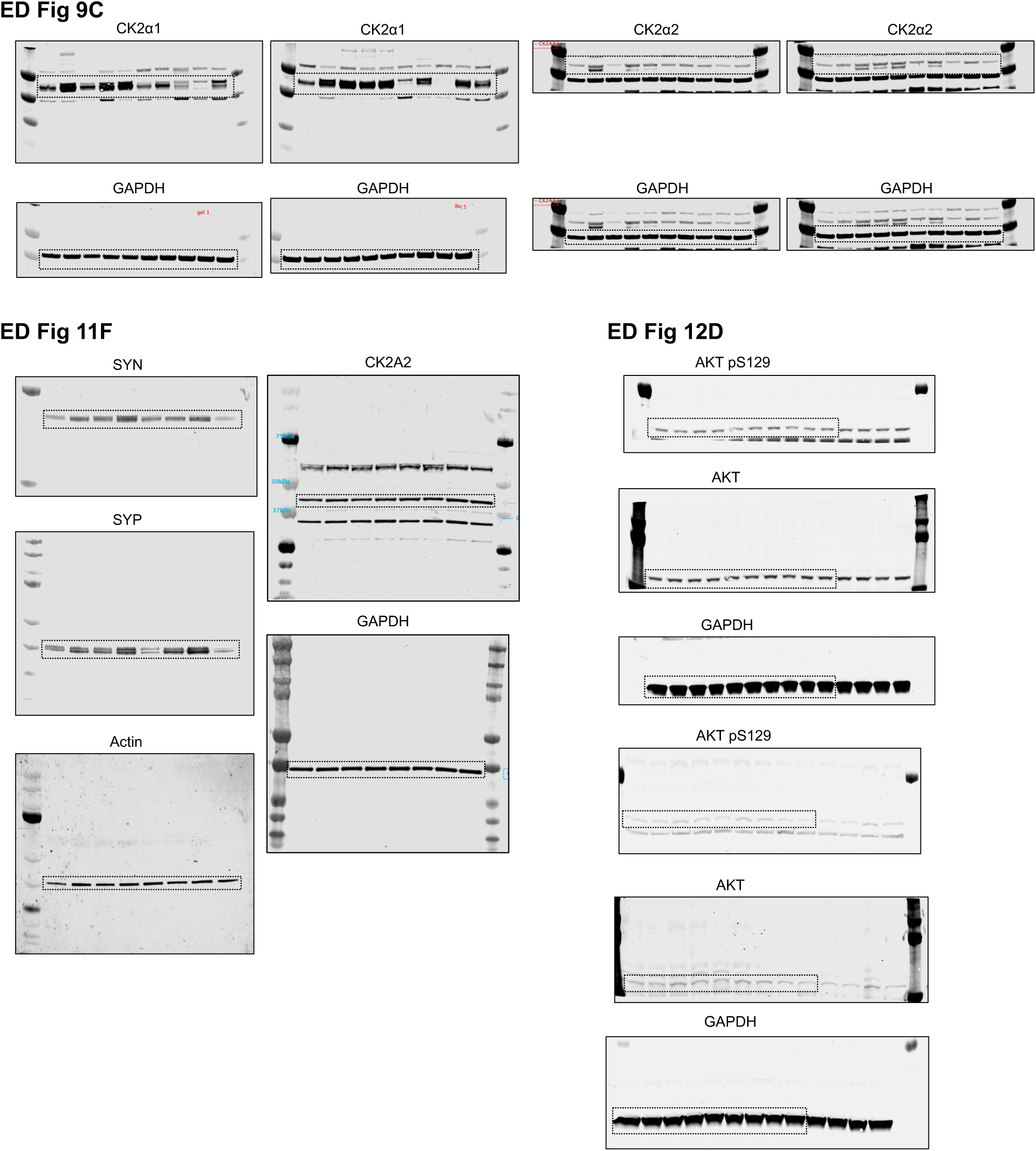
Original full blot images from Odyssey scans (part V). **Extended Data Figures 15-19.** Original full blot images from Odyssey scans. Dotted lines show where blots were cropped to create figures within the main Figures and Extended Data Figures.

